# Global kinome silencing combined with 3D invasion screening of the tumor microenvironment identifies fibroblast-expressed PIK3Cδ involvement in triple-negative breast cancer progression

**DOI:** 10.1101/822049

**Authors:** Teresa Gagliano, Kalpit Shah, Sofia Gargani, Liyan Lao, Mansour Alsaleem, Jianing Chen, Vasileios Ntafis, Penghan Huang, Angeliki Ditsiou, Viviana Vella, Kritika Yadav, Kamila Bienkowska, Giulia Bresciani, Kai Kang, Leping Li, Philip Carter, Graeme Benstead-Hum, Timothy O’Hanlon, Michael Dean, Frances M.G. Pearl, Soo-Chin Lee, Emad A Rakha, Andrew R Green, Dimitris L. Kontoyiannis, Erwei Song, Justin Stebbing, Georgios Giamas

## Abstract

As there is growing evidence for the tumor microenvironment’s (TME) role in tumorigenesis, we sought to investigate the role of fibroblast-expressed kinases in triple negative breast cancer (TNBC). Using a high-throughput kinome screen combined with 3D invasion assays, we identified fibroblast-expressed PIK3Cδ (f-PIK3Cδ) as a key regulator of progression. Although PIK3Cδ has been mainly described in leucocytes, we detected high expression in primary fibroblasts derived from TNBC patients, while PIK3Cδ was undetectable in cancer epithelial cell lines. Genetic and pharmacologic gain- and loss-of functions experiments verified the contribution of f-PIK3Cδ in TNBC cell invasion. By employing an integrated secretomics and transcriptomics analysis, we revealed a paracrine mechanism via which f-PIK3Cδ confers its pro-tumorigenic effects. Inhibition of f-PIK3Cδ promoted the secretion of factors, including PLGF and BDNF, which subsequently led to upregulation of NR4A1 in TNBC cells where it acts as a tumor suppressor. Inhibition of PIK3Cδ in an orthotopic BC mouse model reduced tumor growth only after inoculation with fibroblasts, indicating a role of f-PIK3Cδ in cancer progression. Similar results were observed in the MMTV-PyMT transgenic BC mouse model, in addition to a decrease on tumor metastasis emphasizing the potential immune-independent effects of PIK3Cδ inhibition. Finally, analysis of BC patient cohorts and TCGA datasets identified f-PIK3Cδ (protein and mRNA levels) as an independent prognostic factor for overall and disease free survival, highlighting it as a therapeutic target for TNBC.

## Introduction

Triple negative breast cancer (TNBC; ER^-^, PR^-^, HER2^-^) represents a molecularly diverse and highly heterogeneous subtype of BC (15-20%) with a poor prognosis, high rates of recurrence and metastasis. Treatment largely relies on chemotherapy which remains toxic and often ineffective (1, 2).

Despite mounting evidence for the role of the tumor microenvironment (TME) in affecting surrounding cells and its involvement in metastatic progression, little is known about how stromal cells can influence the behavior of cancer epithelial cells and how they affect their response to target therapy (3–5). Under normal physiological conditions, the stroma serves as a barrier to epithelial cell transformation, while the interplay between epithelial cells and the microenvironment can maintain epithelial polarity and modulate growth inhibition (6). In BC, gene expression analysis of the tumor stroma has led to identification of clusters that can predict clinical outcome (7). In patients with TNBC, infiltration of inflammatory cells or the presence of a stroma with reactive, invasive properties, has been associated with a poor prognosis (8, 9).

Fibroblasts are the most prominent cells in the TME and are thought to induce both beneficial and adverse effects in pre-metastatic progression (4, 10). The important functions of fibroblasts include the deposition of extracellular matrix (ECM), regulation of epithelial differentiation, regulation of inflammation and involvement in cell migration (11, 12). Fibroblast-secreted ECM proteins play a vital role in BC onset and progression (13), while cancer-associated fibroblasts (CAFs) have been shown to promotes resistance to cytotoxic and target therapy by secreting protective factors (14). Further understanding the involvement of stromal cells in TNBC, in particular the elucidation of the cross-talk between fibroblasts and BC cells, might eventually lead to the design of new therapeutic strategies and more effective tailored treatment options for TNBC patients.

Finak et al. have reported that functional inactivation of phosphatase and tensin homolog (PTEN) that leads to phosphoinositide 3-kinase (PI3K) activation, in fibroblasts within the breast TME contributes to cancer development and progression (7). We hypothesized that PI3K activity may be a regulator of the tumor-stroma interactions (15) and inhibition of PI3K signaling in fibroblasts could impede its tumor-promoting activity. PI3Ks phosphorylate inositol lipids and are involved in immune response (16–18). Whereas PIK3Cα (p110α) and PIK3Cβ (p110β) are ubiquitously expressed, PIK3Cδ (p100δ) is predominantly expressed in white blood cells (19). However, an unexpected role of PIK3Cδ in oncogenesis of non-hematopoietic cells was observed in avian fibroblasts where overexpression of wild-type PIK3Cδ induced oncogenic transformation (20, 21).Another report has demonstrated the involvement of PI3K isoforms (including PIK3Cδ) in the differentiation of human lung fibroblasts into myofibroblasts (22). PIK3Cδ contributes to neutrophil accumulation in inflamed tissue by impeding chemoattractant-directed migration as well as adhesive interactions between neutrophils and cytokine-stimulated endothelium (23). Although hampering PI3Ks’ activity in fibroblasts would be expected to inhibit stroma mediated tumor-promoting activity, a direct effect of PI3K inhibitors on these cells has not been tested to date.

Herein, using a high throughput siRNA kinome screening we identify fibroblast-expressed PIK3Cδ as a mediator of TNBC development *in vitro* and *in vivo* and we show the mechanism via which fibroblast PIK3Cδ modulates TNBC progression. Our work reveals a previously uncharacterized yet significant role of fibroblast-expressed PIK3Cδ, which supports the rationale for clinical use of PIK3Cδ inhibitors for the treatment of TNBC.

## Results

### Examination of cancer associated markers in HMF and MRC5 fibroblast cell lines

The primary aim of this work was to examine how normal fibroblasts (NFs) within the TME affect TNBC progression. This reflects several controversial issues that have been raised about the genomic landscape of CAFs and the identification of specific markers that differentiate CAFs. According to recent published evidence (8), CAFs in TNBC should be characterized by the combined expression of fibroblast activated protein (FAP), integrin β1 (ITGβ1; CD29) S100A4, PDGFRβ, α-smooth muscle actin (α-SMA) and Caveolin. Therefore, the expression of CAF markers was evaluated in the fibroblast cell lines used in this study (HMF and MRC5) and were also compared to primary fibroblasts (NFs and CAFs) obtained from four TNBC patients (**Supplementary Table 1**). More specifically, the separation of primary CAFs from NFs was based on the distance from the site of the primary tumor (CAFs<5cm; NFs >5cm).

As shown in **Supplementary Figure 1A**, PDGFRβ was more abundant in CAFs as compared to NFs while Caveolin appeared to be downregulated in CAFs. Overall expression and changes in FAP levels were related to patient’s variability rather than the fibroblast site of origin. ITGβ1 and α-SMA were widely expressed in all samples analyzed, while S100A4 was hardly detectable in primary fibroblasts. The expression levels of the various CAF markers in HMF and MRC5 were comparable to those in the primary fibroblast cell lines. PDGFRβ, ITGβ1, FAP, Caveolin and α-SMA levels were equally detected in both HMF and MRC5 cells while S100A4 was solely present in HMF.

Although the expression of these markers in HMF and MRC5 were comparable to those found in primary fibroblasts of TNBC tumors, there was no clear distinction between NFs and CAFs based on these proteins, which supports the aforementioned controversy. On the other hand, the similarity between HMF/MRC5 and Primary fibroblast in expression of CAF-markers support the use of these cells as a model to study cancer cells-fibroblast interactions. Nevertheless, since our goal is not restricted to a specific subtype of fibroblasts, we proceeded using both cell lines / models (HMF and MRC5) in our experiments.

### High-throughput (HT) RNAi screening identifies fibroblast-expressed kinases involved in TNBC cell invasion

Based on the established role of protein kinases (PKs) as drug targets and taking into consideration the fact that intra- and extra-cellular signaling is mainly transmitted through PKs, we investigated the role of fibroblast-expressed kinases on TNBC progression. Therefore, we established an experimental pipeline as shown in **Figure 1A**, broadly applicable to different systems that consisted of a 3D co-culturing model (cancer and stromal cells) linked to an invasion assay as a readout tool.

**Figure 1:**
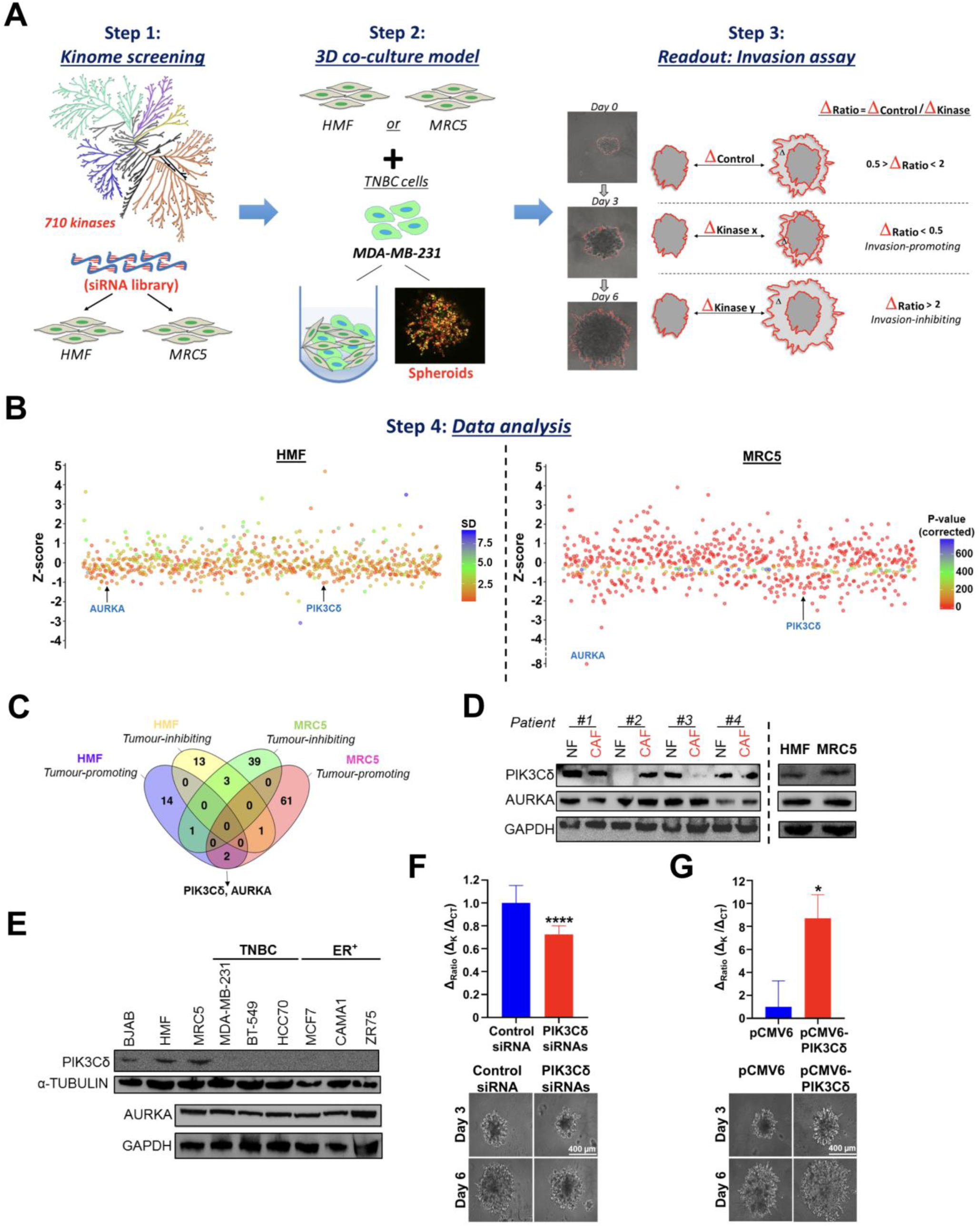
Experimental design of siRNA Kinome screening and identification of fibroblast-expressed kinases affecting TNBC invasion. (**A**) *<u>Step 1</u>*: Silencing of 710 kinases in HMF and MRC5 cells using a siRNA kinome library. <u>*Step 2*</u>: 3D co-culture of HMF or MRC5 with MDA-MB-231 in the presence of Matrigel® and chemoattractants to promote invasion. A representative image of MDA-MB-231 (red) and MRC5 (green) cells stained with different fluorescent lipophilic tracers is shown (Red: DiRDiIC18; Green: DiOC6). *<u>Step 3</u>*: The invasive potential of MDA-MB-231 cells was used as a readout tool. Results are expressed as changes in spheroid surface between Day 6 and Day 3 (Δ_Ratio_= Δ_CT_/Δ_K_). The Δ_Ratio_ values were used to calculate the Z-Scores based on the formula: **z = (x – μ)/σ** (**μ**: Δ_Ratio_ mean of 710 kinases; **σ**: standard deviation (SD); **x**: Δ_Ratio_ value for each kinase). For HMF, the Δ_Ratio_ Z-Score colour code refers to SD, as the screening was performed twice, while for MRC-5 the Δ_Ratio_ Z-Score colour code refers to p-Value. (**B**) *<u>Step 4</u>*: The Z-Score graphs for HMF and MRC5 are shown. Kinases were divided depending on their effects on MDA-MB-231 invasion. ‘Invasion-promoting’: Δ_Ratio_ ≤ 0.5, P < 0.01 (as well as SD < 0.5 for HMF). ‘Invasion inhibiting’: Δ_Ratio_ > 2, P > 0.05 (as well as SD >0.5 for HMF). **(C)** Venn diagrams comparing the number of invasion-promoting and invasion-inhibiting kinases in HMF and MRC5 cells. PIK3Cδ and AURKA were identified as main hits (invasion-promoting) in screenings experiments (Δ_Ratio_ < 0.5; p < 0.01). (**D**) Western blotting of PIK3Cδ and AURKA in HMF, MRC5 and primary fibroblasts obtained from TNBC patients (CAFs<5cm; NFs >5cm distance from the tumor site). GADPH was used as loading control. (**E**) Western blotting of PIK3Cδ and AURKA in BC and fibroblast cell lines (BJAB B-cell-line were used as positive control for PIK3Cδ expression). GADPH and α-tubulin were used as loading controls. (**F**) Validation of effects of PIK3Cδ knockdown in MRC5 (pool of 3 siRNAs), on MDA-MB-231 invasion following the experimental procedure described above (*n*=3 independent experiments, with at least 3 technical replicates). Results are expressed as mean ± SEM. Significance was calculated using Unpaired t-test w; **** P < 0.0001 vs Control siRNA. (**G**). Effects of PIK3Cδ overexpression in MRC5, using pCMV6-AC-PIK3Cδ-GFP plasmid, on MDA-MB-231 invasion following the experimental procedure described above (*n*=3 independent experiments, with at least 3 technical replicates). Results are expressed as mean ± SEM. Significance was calculated using Unpaired t-test; * P < 0.05 vs Control siRNA vs. pCMV6 transfected fibroblasts.

The primary screening was performed in duplicate in HMF and once in MRC5 cells. Fibroblast cell lines were transfected with a pool of 3 siRNAs/gene targeting each of the 710 human kinases (**Figure 1A**; *Step 1*). 24 hours after transfection, HMF or MRC5 were co-cultured in 3D with MDA-MB-231 and after three days required for spheroids formation, Matrigel and chemoattractants were added to the wells to promote invasion (**Figure 1A, Supplementary Figure 2 and Supplementary Videos 1 & 2**; *Step 2*). Pictures of all spheroids taken after 3 and 6 days were analyzed and the results were expressed as changes in spheroid surface (Δ= Surface_Day6_ - Surface_Day3_). The Δ-value of each silenced kinase (Δ_K_) was compared with the Δ- value of the control (Δ_CT_), at different time-points, to obtain a Δ_Ratio_ (Δ_Ratio_= Δ_CT_/Δ_K_) (**Figure 1A**; *Step 3*). Kinases were divided depending on their effects on MDA-MB-231 invasion and those with a ΔRatio ≤ 0.5 (50% less invasion vs CT) and p<0.01 (as well as SD < 0.5 for HMF), were considered as ‘invasion-promoting’, while kinases with a ΔRatio > 2 (100% more invasion vs CT), P > 0.05 (as well as SD > 0.5 for HMF) were considered as ‘invasion inhibiting’ ones. The Δ_Ratio_ values were used to calculate the Z-Scores and all hits were plotted for both cell lines, revealing new potential fibroblast-expressed kinases that are able to modulate TNBC cell invasion (**Figure 1B and Supplementary Figure 3**; *Step 4*). All siRNA screening data are presented in **Supplementary Table 2**.

Based on pre-specified cut-off criteria, we identified 17 kinases in HMF and 64 kinases in MRC5 whose silencing decreased the rate of TNBC invasion (∼40%-90%), suggesting a pro-invasive role of these proteins (**Figure 1C**). Under these conditions, there were two shared targets amongst HMF and MRC5, namely PIK3Cδ and AURKA. Using a panel of fibroblasts and breast cancer (BC) cells, we analyzed the levels of PIK3Cδ and AURKA and discovered a variability in their expression amongst the primary and immortalized fibroblast cell lines (**Figure 1D**, **Supplementary Figure 1B**). PIK3Cδ protein levels in fibroblast cells were comparable to those in the BJAB B-cell line (used as a positive control) (24) while intriguingly PIK3Cδ was hardly detectable or totally absent in most of the BC cells, as opposed to AURKA, which was ubiquitously expressed (**Figure 1E, Supplementary Figure 1C and Supplementary Figure 4D-*upper panel***). qRT-PCR analysis of PIK3Cδ revealed a similar trend for most of the cell lines tested (**Supplementary Figure 4A**), though it is well-known that protein and mRNA abundances do not always correlate (25, 26). Moreover, RNA sequencing (RNAseq) in a variety of different organs obtained from the Human Protein Atlas (27) revealed that apart from myeloid and lymphoid cells, fibroblast cell lines express moderate/high PIK3Cδ mRNA levels in contrast to BC cell lines that have low/negligible mRNA transcripts (**Supplementary Figure 4B**). We also investigated whether fibroblast-PIK3Cδ can induce the expression of PIK3Cδ in TNBC following extended co-culturing between the different cell types. As shown in **Supplementary Figures 4C and 4D**, by using both fibroblast cell lines and primary CAFs derived from MMTV-PyMT tumors, there were no changes of PIK3Cδ in TNBC cells, maintaining their low/undetectable protein levels.

Altogether, these results suggest that PIK3Cδ could not have been identified if we were solely studying BC cells instead of examining their interactions with the surrounding stroma, further supporting the setup of our experimental approach by regarding cancer as a systemic multi-cell lineage dependent disease. We further validated the involvement of fibroblast PIK3Cδ-mediated TNBC 3D invasion by repeating the experiment following either silencing (**Figure 1F**) or overexpressing (**Figure 1G**) PIK3Cδ. Similar data were obtained using MDA-MB-231, BT-549 and fibroblast cell lines (**Supplementary Figures 5A-5F**). Finally, we also determined that treatment of TNBC cells with conditioned media (CM) derived from genetically modified fibroblasts (PIK3Cδ-silenced or PIK3Cδ-overexpressed) has no significant effect on TNBC cell proliferation (**Supplementary Figure 5G**). Taking everything into consideration and bearing in mind the potential implication in BC we focused on PIK3Cδ and investigate its stromal (fibroblast) involvement.

### Confirmation of HT-RNAi screening results

The accuracy and validity of our experimental pipeline/screening was supported by identification of kinases (positive hits) whose involvement in stromal-mediated cancer invasion has been previously reported. Amongst these results were FLT4 (28) and EGFR (29) (invasion-promoting) as well as ACVR1B (30) and ITPKB (31) (invasion-inhibiting).

To further validate the of HT-RNAi screenings results, we performed the aforementioned experimental pipeline, this time by using shRNA plasmids targeting randomly selected kinases. As shown in **Supplementary Figure 6A**, the effects of shRNA-mediated silencing of the tested kinases in MRC5 led to similar effects on the invasion of MDA-MB-231 as observed in the primary screening (using specific siRNAs). The gene knockdown efficiency of the used shRNAs was confirmed by real-time qRT-PCR (**Supplementary Figure 6B**).

Next, we examined the effects of certain kinase inhibitors and repeated the experiment. As anticipated, similar results were observed, although in some cases (i.e. AZD4547) the results were not identical to the primary screening (**Supplementary Figures 6C**). This could be due: (i) to potential off-target effects of some inhibitors, (ii) to the fact that some of these inhibitors can target other isoforms of a specific kinase and/or (iii) because the genomic vs the chemical/pharmacological inhibition of a kinase does not necessarily have the same outcome. In relation to the overlapping hits from our screening (PIK3Cδ and AURKA), we verified that the observed effects on MDA-MB-231 cell invasion, following genomic inhibition (siRNA), is not based on a reduction (**Supplementary Figure 6D**) or an increase (in the case of PIK3Cδ overexpression; **Supplementary Figure 6E**) of cell viability.

### Genomic or chemical inhibition of PIK3Cδ in fibroblasts reduces TNBC cell invasion as a result of paracrine signaling

To clarify whether the catalytic activity of PIK3Cδ is required for its effect on TNBC progression, we repeated our 3D spheroid invasion assay following chemical inhibition of PIK3Cδ in fibroblast cells, using CAL-101 (Idelalisib; a highly selective and potent PIK3Cδ inhibitor) (32). HMF or MRC5 were initially treated with different concentrations of CAL-101 for 24h, while the efficacy of CAL-101 inhibition on downstream targets of PIK3Cδ (mTOR/Ser2448, PDPK1/Ser281 and AKT/Thr308) was validated (**Supplementary Figure 7A**). Moreover, treatment with CAL-101 had limited or no effects on fibroblasts’ cell viability for the 24h period of treatment (**Supplementary Figure 7B**). Nevertheless, to avoid any misinterpretations, fibroblasts were washed with PBS, counted with trypan blue, and only viable cells were used in co-cultures with TNBC cells (MDA-MB-231 or BT-549) at a 1:1 ratio. As shown in **Figure 2A and Supplementary Figure 8A**, CAL-101 pre-treated fibroblasts showed a decrease in 3D-spheroid invasion, suggesting that the kinase activity of PIK3Cδ is, to a great extent, responsible for the observed results.

**Figure 2:**
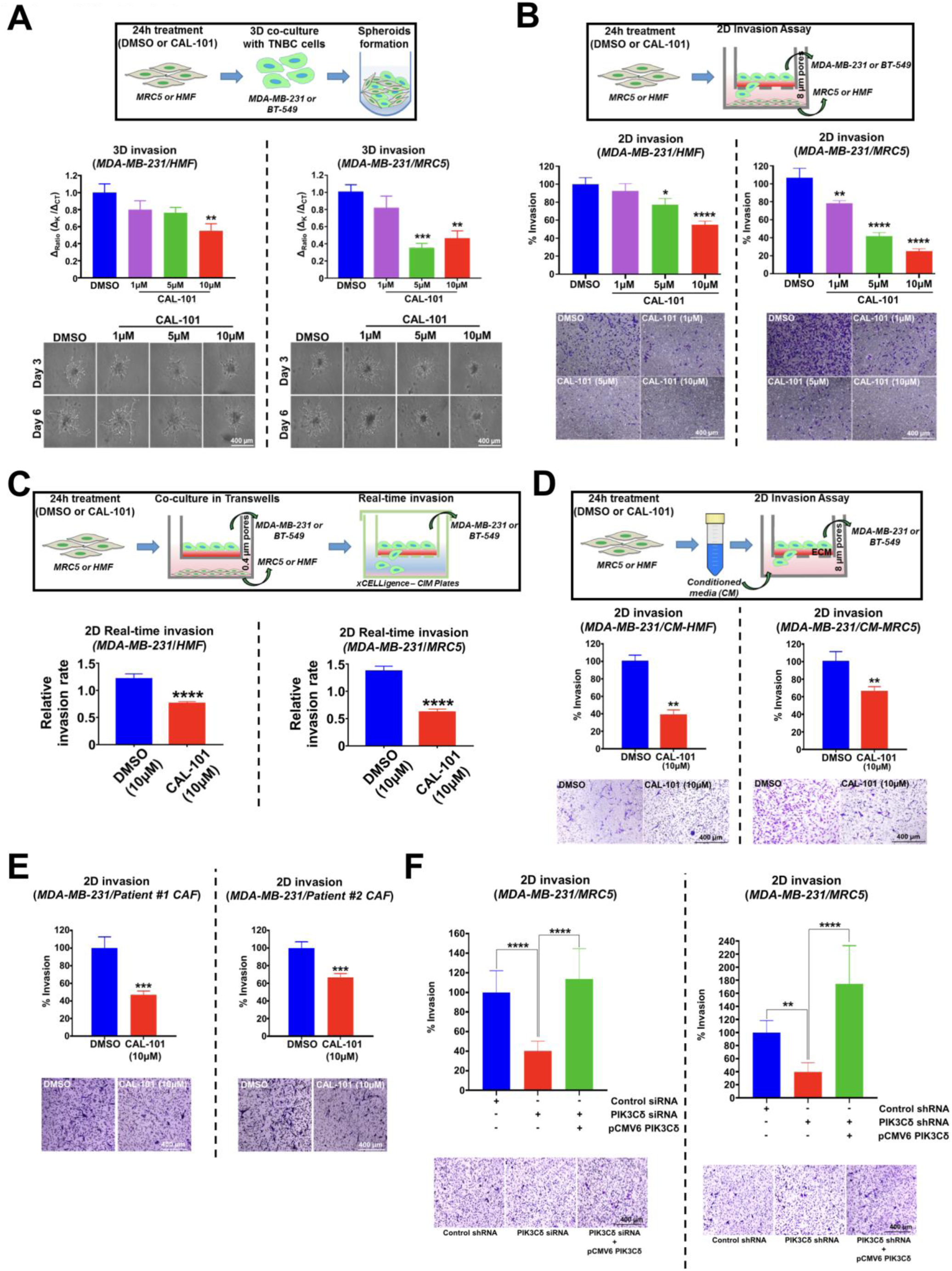
Effects of chemical inhibition of PIK3Cδ on TNBC 2D and 3D invasion. (**A**) 3D invasion assay: HMF (left panel) and MRC5 (right panel) were pre-treated with either vehicle (DMSO) or with 1, 5 and 10 µM of CAL-101 for 24h. Following, fibroblasts were co-cultured with MDA-MB-231 as 3D spheroids and the invasion potential was measured. Representative pictures of the 3D invasion assay at different time points are shown (*n*=3 independent experiments, with at least 4 technical replicates). Significance was calculated using one-way Anova and Tukey’s multiple comparisons tests. Results are expressed as mean ± SEM; ** P < 0.01, *** P < 0.001 vs. DMSO treated fibroblasts. (**B**) 2D invasion assay: HMF (left panel) and MRC5 (right panel) were pre-treated with either vehicle (DMSO) or with 1, 5 and 10 µM of CAL-101 for 24h and were then seeded on the lower chamber of a transwell insert. MDA-MB-231 cells were seeded on the Matrigel-coated upper chamber of the transwell insert (pore size: 8 μm allowing cell-crossing) and were co-cultured with the fibroblasts. After 24h of co-culture, migrated MDA-MB-231 cells were fixed and stained with crystal violet and counted using an inverted microscope (n=3 independent experiments, with at least 3 technical replicates) Significance was calculated using one-way Anova and Tukey’s multiple comparisons tests. Data are expressed as mean ± SEM; * P < 0.05, ** P < 0.01, **** P < 0.0001 vs. DMSO treated fibroblasts. (**C**) 2D real-time invasion assay: HMF (left panel) and MRC5 (right panel) were pre-treated with either vehicle (DMSO) or 1, 5 and 10 µM of CAL-101 for 24h and were then seeded on the lower chamber of a transwell insert. MDA-MB-231 cells were seeded on the upper chamber of the transwell insert (pore size: 0.4 μm not allowing cell-crossing) and were co-cultured with the fibroblasts. After 24h of co-culture, MDA-MB-231 were moved to CIM-Plates and their relative invasion rate was monitored using the xCELLigence Real-Time Cell Analysis (RTCA) instrument Significance was calculated using Unpaired t-test. Results are expressed as mean ± SEM; **** P < 0.0001. **(D)** 2D CM invasion assay: HMF (left panel) and MRC5 (right panel) were treated with vehicle only or with 10 µM CAL-101 in serum free for 24h in order to obtain the conditioned medium (CM). MDA-MB-231 were then incubated with HMF or MRC5 CM for 2D invasion assays (n=3 independent experiments, with at least 10 technical replicates). Significance was calculated using unpaired t-test. Data are expressed as mean ± SEM; ** P < 0.01 vs DMSO-treated fibroblasts’ CM. (**E**) 2D invasion assay: TNBC patients’ CAFs were pre-treated with either vehicle (DMSO) or with 10 µM of CAL-101 for 24h and were then seeded on the lower chamber of a transwell insert. MDA-MB-231 cells were seeded on the Matrigel-coated upper chamber of the transwell insert (pore size: 8 μm allowing cell-crossing) and were co-cultured with the TNBC CAFs. After 24h of co-culture, migrated MDA-MB-231 cells were fixed and stained with crystal violet and counted using an inverted microscope (n=2 independent experiments, with at least 5 technical replicates). Significance was calculated using unpaired t-test. Data are expressed as mean ± SEM; **** P< 0.0001 vs DMSO treated fibroblasts. (**F**) MRC5 were transfected with control or PIK3Cδ siRNA (left panel) or control or PIK3Cδ shRNA (right panel). MDA-MB-231 cells were incubated with MRC5 cells and 2D invasion assays were performed. Silenced MRC5 were transfected with either pCMV6-AC-GFP vector or with pCMV6-AC-PIK3Cδ-GFP plasmid. MDA-MB-231 cells were then incubated with MRC5 cells and 2D invasion assays were performed (n=2 independent experiments, with at least 5 technical replicates). Significance was calculated using one-way Anova and Tukey’s multiple comparisons tests. Data are expressed as mean ± SEM; ** P < 0.01 ****P < 0.0001 vs samples indicated in the graph.

Intercellular communication sets the pace for transformed cells to survive and to thrive. Based on the initial setup of our assay (3D-spheroids/cells co-culture), we could not be certain whether the involvement of stromal PIK3Cδ on TNBC progression is a result of juxtacrine signaling (cell-to-cell contact-dependent) or a consequence of paracrine signaling due to secreted factors derived from fibroblasts that can alter the behavior of TNBC cells. Hence, we implemented a transwell assay, where HMF or MRC5 cells pre-treated with CAL-101, were seeded on the lower chamber of the transwell inserts and 24h later co-cultured with TNBC cells (MDA-MB-231 or BT-549) platted on a Matrigel-coated top chamber to assess the 2D invasion potential of TNBC cells (**Figure 2B and Supplementary Figure 8B**). Using a similar experimental principle, we employed the xCELLigence Real-Time Cell Analysis (RTCA) instrument (33), to monitor the real-time invasion rate of TNBC cells following co-culturing with fibroblasts pre-treated with CAL-101 (**Figure 2C and Supplementary Figure 8C**). Both assays revealed a reduction in invasiveness after inhibition of fibroblast-PIK3Cδ activity. Similar results when observed when we implemented another method for indirect contact co-cultures using the CM of CAL-101 treated HMF or MRC5 and examining their effects on TNBC invasion (**Figure 2D and Supplementary Figure 8D**). Moreover, 2D invasion assays of either MDA-MB-231 or BT-549 cells co-cultured with primary TNBC CAFs showed comparable results further highlighting the role of fibroblast-PIK3Cδ (**Figures 2E and Supplementary Figure 8E**).

To further demonstrate the contribution of PIK3Cδ in the observed phenotype and rule out any off-target effects, we repeated the 2D invasion assays by initially silencing PIK3Cδ (siRNA or shRNA) and then performed a recovery experiment by re-introducing (overexpressing) PIK3Cδ. As anticipated, by recovering PIK3Cδ levels (**Supplementary Figure 9**), we were able to reverse the decrease in invasion that was induced by genetic inhibition of PIK3Cδ (**Figure 2F**).

Next, to examine the possibility of other fibroblast-expressed PI3K isoforms contributing to the decrease of TNBC cell invasion, in particular PI3KCγ that can also be inhibited by CAL-101, we repeated the 2D invasion assays using various PI3K inhibitors (**Supplementary Figure 10A**). Treatment with AS252424 (PIK3Cγ/α inhibitor) had minor effects on the invasion of TNBC cells, as compared to either CAL-101 or Leniolisib (PIK3Cδ inhibitor). Furthermore, use of the pan-PIK3C inhibitors, PI-103 and NVP-BEZ235 had analogous effects with the PIK3Cδ inhibitors, further supporting the importance of PIK3Cδ in the observed phenotype (**Supplementary Figure 10B**). Finally, we also verified that the observed effects on MDA-MB-231 cell invasion, following treatment with the various PIK3 inhibitors, is not based on a reduction of cell viability (**Supplementary Figure 10C**).

Taken together all combinations of cell lines and assays used, our results demonstrated that fibroblast-PIK3Cδ is able to promote TNBC progression via paracrine regulatory mechanisms.

### Integrated secretome and transcriptomic analyses reveal fibroblast PIK3Cδ-mediated paracrine mechanisms that promote TNBC progression

We and others (34, 35) have shown that co-culture of stromal with BC cells can lead to changes in expression of proteins supporting the hypothesis of cross-talk between different cell types. Changes of PIK3Cδ activity in fibroblasts can alter the intercellular communication between stromal and cancer cells in turn affecting their biological properties. To gain insights into the paracrine mechanisms employed by fibroblasts to promote invasion in TNBC cells, we performed an integrated analysis of proteins secreted by fibroblasts (HMF and MRC5) treated with CAL-101 and the transcriptome of MDA-MB-231 cells which were grown in a transwell setup with CAL-101 treated MRC5 cells.

Based on our 2D/3D co-culture results, we analyzed the PIK3Cδ-regulated secretome, using the Human L1000 Array, which consists of 1000 proteins including cytokines, chemokines, adipokines, growth and angiogenic factors, proteases, soluble receptors and adhesion molecules. HMF and MRC5 cells were treated with either vehicle (DMSO) or 10 µM CAL-101 for 24h and cell culture supernatants were isolated and processed according to the manufacturer’s instructions. By employing differential expression analysis, we identified a total of 206 and 377 secreted proteins that were significantly regulated by CAL-101 treatment of HMF and MRC5 respectively at the Log2 fold difference of |0.5| and a P-value of ≤ 0.05. We found that 73 secreted proteins were common between CAL-101 treated HMF and MRC5 cells, providing evidence for a mechanism of fibroblast-mediated regulation of TNBC aggressiveness (**Figure 3A**, **Supplementary Table 3**). To gain additional insights into the similarities and differences in CAL-101 mediated effects on the secretome, we generated an upset plot of differentially expressed secreted proteins from CAL-101 treated HMF and MRC5 cells. As shown in **Figures 3B and 3C**, CAL-101 upregulated a common set of 40 proteins and downregulated a set of 5 proteins in both HMF and MRC5 cells, while 28 proteins were differentially regulated by CAL-101 in HMF and MRC5.

**Figure 3:**
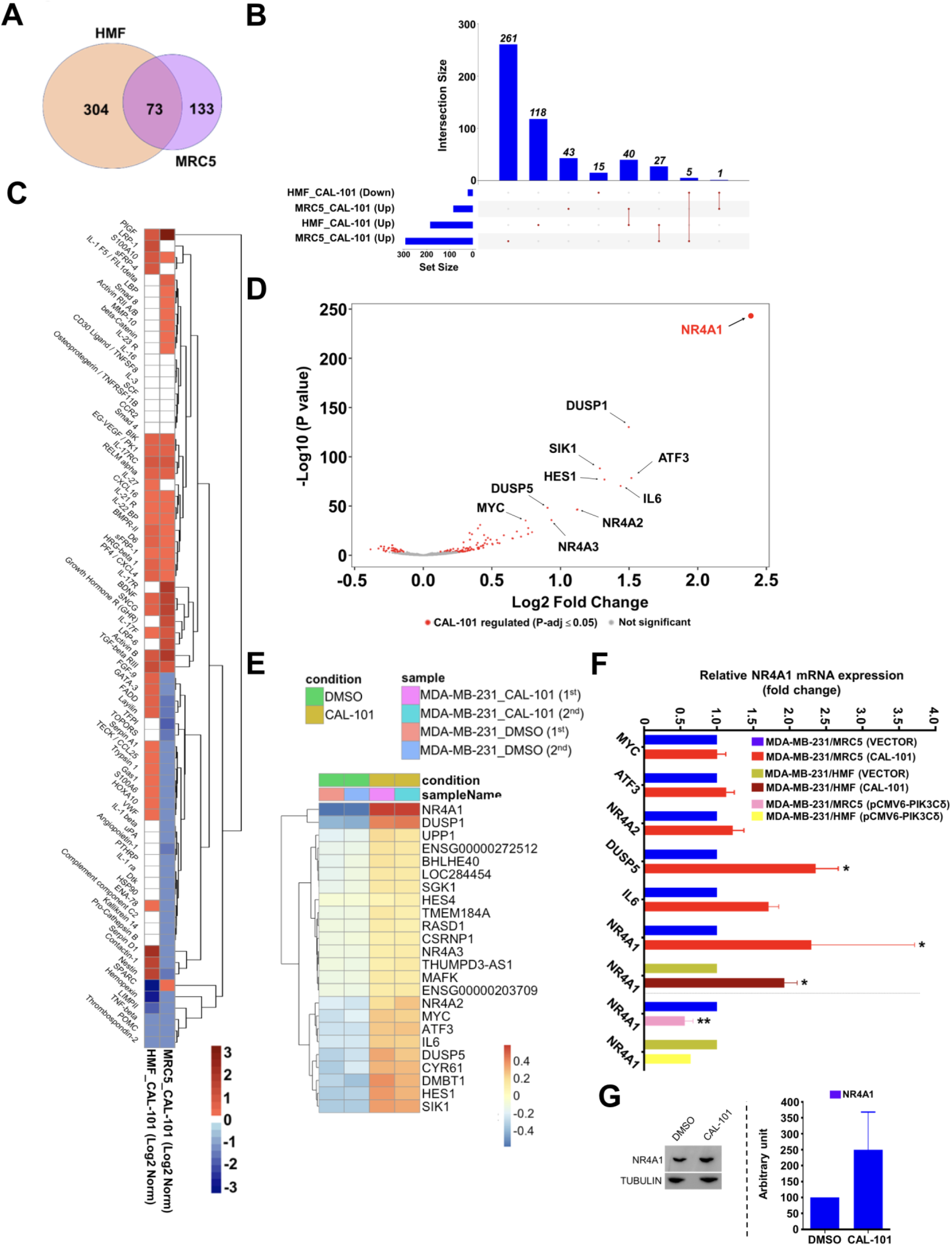
Global secretome analysis of CAL-101 treated fibroblasts and transcriptomics analysis of MDA-MB-231 cells. (**A**) To obtain CM from HMF and MRC5, cells were treated with vehicle only or with 10 µM CAL-101 in serum free medium for 24h. CM was used to perform secretome analysis using the Human L1000 array. Venn diagram showing an overlap as well as differences in the secreted proteins significantly regulated by CAL-101 in HMF and MRC5 cells (Padj < 0.05 and Log2 fold difference of ≥|0.5|) (**B**) The UpSet plot showing common and unique CAL-101 regulated proteins significantly up (Up) or down-regulated (Down) in each dataset (MRC5 and HMF). (**C**) Heatmap comparing Log2 fold change of secreted proteins common between CAL-101 treated HMF and MRC5 cells. (**D**) MRC5 were treated with either vehicle (DMSO) or with 10 µM CAL-101 for 24h hours. Following, cells were washed with PBS to remove the treatment and complete fresh medium was added to each well. 5 µm inserts containing MDA-MB-231 were then added in the well containing previously treated MRC5. 24h after co-culture, cancer cells were collected for RNA extraction and subsequent RNA sequencing. Volcano plot showing the Log2 fold change of genes in MDA-MB-231 cells that responded differently to CAL-101 treatment on MRC5 (DMSO:CAL-101). The Log10 of P value, for significance in fold change, is plotted on the y-axis. (**E**) Heatmap showing amounts by which the read counts of the top-24 (ordered based on Log2 fold change ≥ |0.5| and Padj value ≤ 0.05) regulated genes deviates from the genes’ average across all the samples. (**F**) qRT-PCR validation of genes identified via the RNAseq and DESeq2 analysis. Significance was calculated using unpaired t-tests. Results are expressed as mean ± SEM; *P < 0.05 vs. vector. (**G**) Western blotting of NR4A1 in MDA-MB-231 cells following co-culture with CAL-101 treated MRC5 cells. α-tubulin was used as loading control. Densitometry analysis of the blot is displayed as a ratio between CAL-101-treated vs. DMSO-treated cells.

With comprehensive profiling of CAL-101 mediated changes in the secretome of fibroblast cell lines established, next we investigated how these secreted proteins altered the transcriptional state of MDA-MB-231 cells. We cultured MDA-MB-231 in a transwell along with CAL-101 or vehicle-treated MRC5 cells for 24 hours and total RNA was extracted from MDA-MB-231 and processed as described earlier (36). As anticipated, whole transcriptome data showed a high degree of similarity between replicate samples and most significant variations between MDA-MB-231 cells co-cultured with CAL-101 treated or untreated MRC5 cells (**Supplementary Figure 11**). The principal component analysis (PCA) support our hypothesis that inhibition of PIK3Cδ in fibroblasts can have a paracrine effect on TNBC cells. We next employed differential gene expression analysis using the DESeq2 pipeline to identify genes dysregulated in MDA-MB-231 cells as a consequence of inhibiting PIK3Cδ in MRC5 cells. We found 137 genes here at the false discovery rate Padj ≤ 0.05 (**Figure 3D and Supplementary Table 4**). Surprisingly, of these 137 genes, only 24 genes were found to be significantly dysregulated at the false discovery rate Padj ≤ 0.05 and Log2 fold difference of ≥ |0.5| (**Figure 3E**). We validated the RNASeq analysis using an orthogonal approach of real-time qRT-PCR in independent experiments using a separate cohort including the effects of overexpression of PIK3Cδ on NR4A1 mRNA levels (**Figure 3F, and Supplementary Table 5**). Amongst the most significantly modulated genes was NR4A1 transcription factor, which was recently reported to be implicated in TNBC invasion (37). The increase on NR4A1 protein levels in MDA-MB-231 cells following co-culture with CAL-101 treated MRC5 cells was also confirmed (**Figure 3G**).

Overall, these results show that pharmacological inhibition of PIK3Cδ in fibroblast cells not only alters its secretome, but also has a subtle paracrine effect on the gene expression of epithelial cancer cells.

### Promotion of TNBC invasion via the fibroblast/epithelial-mediated PIK3Cδ-PLGF/BDNF-NR4A1 signaling pathway

Following confirmation of NR4A1 protein expression in a panel of BC cell lines (**Supplementary Figure 12A**) and to further demonstrate its involvement in TNBC, we treated cells with cytosporone B, an agonist of NR4A1 (38) and examined its effects on the invasiveness of MDA-MB-231 and BT-549 cells. As expected, cytosporone B significantly decreased the invasiveness of TNBC cells (**Figure 4A and Supplementary Figure 12B**). On the contrary, silencing of NR4A1 increased the invasive ability of MDA-MB-231 cells (**Figure 4B**). In addition, we investigated the effects of PIK3Cδ overexpression in MRC5 cells on the invasiveness of MDA-MB-231 cells that were pre-treated with cytosporone B in order to increase their NR4A1 levels. As shown in **Figure 4C**, PIK3Cδ partly rescued the inhibitory effects of NR4A1 activation, demonstrating the PIK3Cδ-NR4A1 paracrine signaling axis between fibroblast and TNBC epithelial cells. In order to examine if the effects of PIK3Cδ inhibition (CAL-101) on TNBC cell invasion are related to NR4A1 expression, we performed a 2D-invasion assay in which MRC5 (or HMF) were treated with CAL-101 (or DMSO), while NR4A1 was silenced in MDA-MB-231 cells. As shown in **Figure 4D and Supplementary Figure 12C**, silencing of NR4A1 completely abrogated the effects of CAL-101 on TNBC cell invasion. Moreover, the NR4A1 silencing-mediated induction of invasion was abolished when fibroblasts were pre-treated with CAL-101 (**Figure 4D**), which is due to the paracrine induction of NR4A1 expression caused by CAL-101 (**Figure 3F**) that balances the NR4A1 siRNA knock-down. The mRNA levels of NR4A1 were indeed rescued after co-culture with CAL-101 treated MRC5 (**Figure 4D**; right panel). Taken together, these results suggest the existence of a direct association between fibroblast PIK3Cδ-mediated reduction of invasion and TNBC cells’ NR4A1 levels.

**Figure 4:**
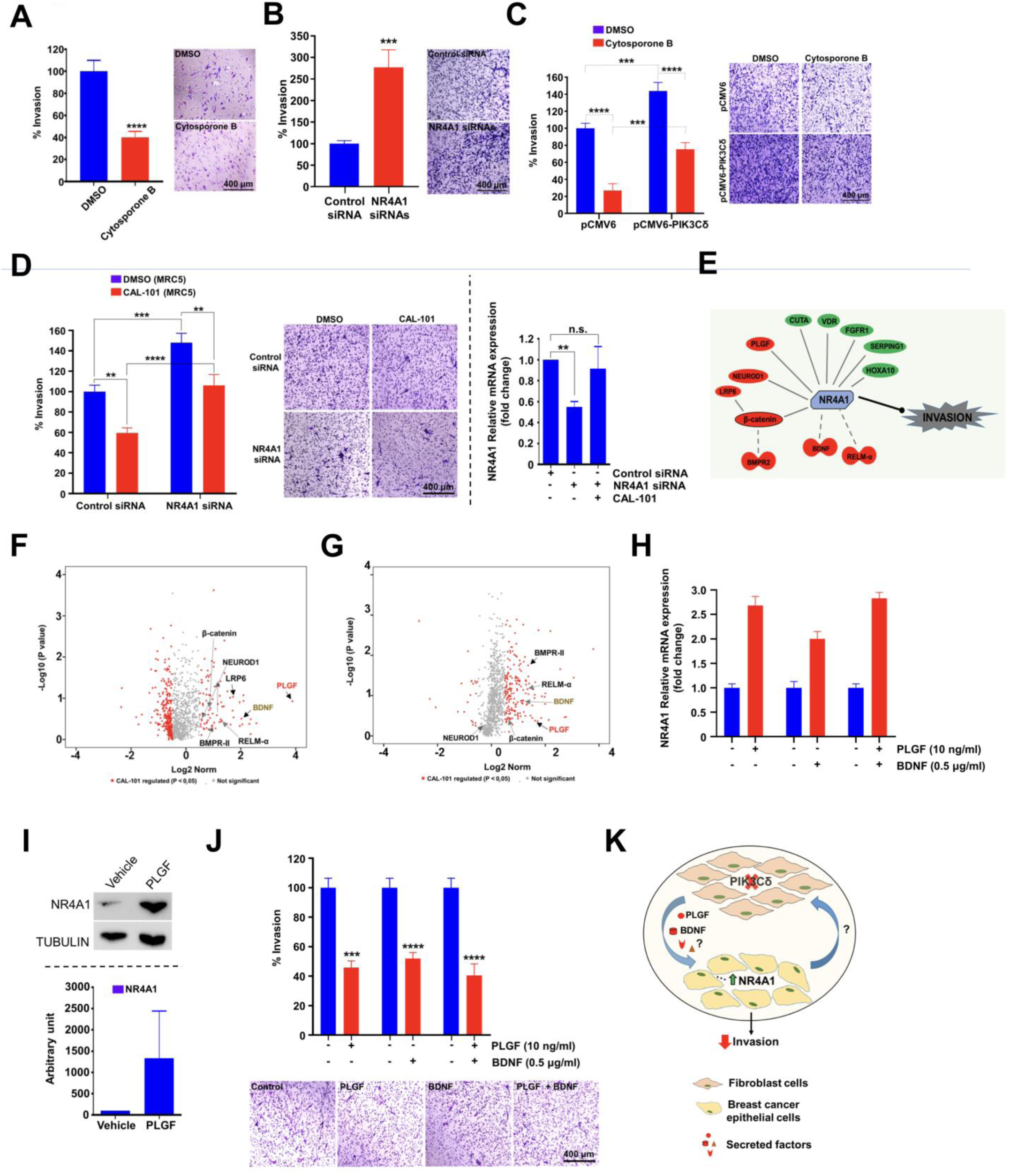
*In silico* prediction and validation of the fibroblast PIK3Cδ-PLGF/BDNF paracrine pathway altering NR4A1-mediated invasion of TNBC cells. (**A**) 2D invasion assay: MDA-MB-231 cells were seeded on the Matrigel-coated upper chamber of the transwell insert (pore size: 8 μm allowing cell-crossing) and were treated with either vehicle (DMSO) or Cytosporone B (5 µM). After 24h, migrated MDA-MB-231 cells were fixed and stained with crystal violet and counted using an inverted microscope (n=2 independent experiments, with at least 9 technical replicates). A representative experiment is displayed. Significance was calculated using unpaired t-test. Results are expressed as mean ± SEM; **** P < 0.0001 vs. DMSO treated cells. (**B**) 2D invasion assay: MDA-MB-231 cells were seeded on the Matrigel-coated upper chamber of the transwell insert (pore size: 8 μm allowing cell-crossing) and were treated with either control or NR4A1 siRNAs (pool of 2 siRNAs). After 24h, migrated MDA-MB-231 cells were fixed and stained with crystal violet and counted using an inverted microscope (n=2 independent experiments, with at least 9 technical replicates). Significance was calculated using unpaired t-test. Results are expressed as mean ± SEM; *** P < 0.001 vs. siRNA control transfected cells. (**C**) Effects of PIK3Cδ overexpression in MRC5, using pCMV6-AC-PIK3Cδ-GFP plasmid, on MDA-MB-231 invasion following pre-treatment of MDA-MB-231 with 5 µM Cytosporone B. (n=2 independent experiments, with at least 9 technical replicates). Significance was calculated using two-way Anova and Tukey’s multiple comparisons tests. Results are expressed as mean ± SEM; *** P < 0.001, **** P < 0.0001 vs the samples indicated in the graph. (**D**) Left/middle panels: MRC5 cells were treated with CAL-101 or DMSO. NR4A1-silenced (siRNA) MDA-MB-231 or control-siRNA cells were seeded on the Matrigel-coated upper chamber of a transwell insert (pore size: 8 μm allowing cell-crossing) and were co-cultured with fibroblasts. After 24h of co-culture, migrated MDA-MB-231 cells were fixed and stained with crystal violet and counted using an inverted microscope (n=3 independent experiments, with at least 6 technical replicates). Significance was calculated using two-way Anova and Tukey’s multiple comparisons tests. Results are expressed as mean ± SEM; ** P < 0.01, *** P < 0.001, **** P < 0.0001 vs the samples indicated in the graph. Right panel: NR4A1 levels were evaluated in siRNA transfected MDA-MB-231 before and after co-culture with CAL-101 treated MRC5. Significance was calculated using unpaired t-tests. Results are expressed as mean ± SEM; **P < 0.01 vs. control. (**E**) Custom pathway built using the IPA showing how secreted proteins from CAL-101 treated MRC5 cells may regulate cell invasion of MDA-MB-231 cells via upregulation of NR4A1 pathway. The red and green colour of the secreted factors indicates increased and decreased expression respectively while the red arrow signifies activation. (**F**) Volcano plot showing the Log2 fold change of secreted proteins in MRC5 cells that responded differentially to the CAL-101 treatment. The Log10 of P value, for significance in fold change, is plotted on the y-axis. **(G)** Volcano plot showing the Log2 fold change of secreted proteins in HMF cells that responded differentially to the CAL-101 treatment. The Log10 of P value, for significance in fold change, is plotted on the y-axis. (**H**) qRT-PCR of *NR4A1* expression levels in MDA-MB-231 cells following treatment with PLGF and BDNF (or control PBS/BSA (0.1%)). (**I**) Western blotting of NR4A1 in MDA-MB-231 cells following treatment with PLGF (10ng/ml) for 24h. GAPDH was used as loading control. Densitometry analysis of the blot is displayed as a ratio between PLGF-treated vs. vehicle-treated cells. (**J**) 2D invasion assay: MDA-MB-231 cells were seeded on the Matrigel-coated upper chamber of the transwell insert (pore size: 8 μm allowing cell-crossing) and were treated with PLGF or BDNF (or control PBS/BSA (0.1%)). After 24h, migrated MDA-MB-231 cells were fixed and stained with crystal violet and counted using an inverted microscope (n=3 independent experiments, with at least 4 technical replicates). Significance was calculated using unpaired t-test. Results are expressed as mean ± SEM; *** P < 0.001, **** P < 0.0001 vs. vehicle treated cells. (**K**) Schematic model depicting the paracrine signaling pathway between fibroblasts and TNBC cells. Inhibition of PIK3Cδ in fibroblasts leads to the secretion of different factors, including PLGF and BDNF, which promote the overexpression of *NR4A1* in epithelial cancer cells. NR4A1 acts as a tumor suppressor inhibiting the invasiveness of TNBC cells.

Having built a comprehensive dataset of secreted proteins from CAL-101 treated MRC5 cells and the transcriptome of MDA-MB-231 co-cultured with CAL-101 treated MRC5 cells, we performed an integrated transcriptomics-proteomics analysis to unravel mechanisms employed by stromal cells to promote invasion in malignant cells.

We hypothesized that secreted factors derived from CAL-101 treated MRC5 and HMF cells may alter cell-cell communication pathways and regulate the invasion of MDA-MB-231 cells by modulating the expression of invasion related genes including *NR4A1*, which was shown to be overexpressed in our transcriptomic analysis (**Supplementary Table 4**). To test this hypothesis, we employed the Ingenuity Pathway Analysis (IPA) software and literature mining to curate a list of potential PIK3Cδ-regulated secreted proteins that are enriched in cell migration/invasion pathways and are also known to modulate expression of NR4A1. Our analytical method is described in **Supplementary Figure 13**. Our approach identified several secreted factors (n=40) that were directly involved in pathways regulating cellular movement or cell-to-cell signaling mechanisms (**Supplementary Table 6**), amongst which certain of them have been previously reported to have an association with *NR4A1* expression, including placental growth factor (PLGF) (39, 40) and brain-derived neurotrophic factor (BDNF) (41) (**Figure 4E**). Scatter plots displaying all secreted proteins from CAL-101 treated MRC5 and HMF while highlighting the list of potential candidates regulating NR4A1 expression in MDA-MB-231 are shown in **Figures 4F and 4G**.

Previous studies have demonstrated that induction of NR4A1 by PLGF inhibits endothelial cell proliferation (42, 43), while PLGF can also impede tumor growth and metastasis (44). Moreover, BDNF has been described to have a role in BC progression (45), even though its exact role has not been completely clarified. Treatment of MDA-MB-231 or BT-549 cells with PLGF or BDNF confirmed its positive effects on NR4A1 supporting their contribution in the paracrine upregulation of NR4A1 mRNA/protein levels (**Figures 4H, 4I and Supplementary Figure 14A**). Moreover, as anticipated, PLGF and BDNF also led to a significant decrease in TNBC invasion (**Figure 4J and Supplementary Figure 14B**). Finally, we confirmed that silencing of either BDNF or PLGF in fibroblasts attenuated the CAL-101-mediated reduction of TNBC cell invasion, further supporting the involvement of this pathway, while suggesting the existence of additional mechanisms implicated in the described phenotype (**Supplementary Figure 15**). Interestingly, silencing of BDNF and/or PLGF did not affect TNBC invasion, suggesting that the PIK3Cδ effects are exerted during membrane trafficking and/or secretion of these factors and not at the gene/protein expression level. This is in accordance with previous reports describing a role of CAL-101 in down-regulating secretion, rather than expression, of chemokines in stromal co-cultures (32). In addition, silencing of BDNF and/or PLGF abolished the CAL-101 inhibitory effects on fibroblast-mediated invasion, without affecting basal invasion levels. This may be due to the experimental design, since silencing of a gene does not have an immediate effect on the respective total and/or secreted protein levels. Finally, it is worth mentioning that other mechanisms / factors could also contribute to the fibroblast-mediated invasion.

In summary, our results reveal a novel paracrine signal transduction pathway between fibroblasts and TNBC cells, encompassing PIK3Cδ-PLGF/BDNF-NR4A1 (**Figure 4K**), without ruling out the existence of other mechanisms contributing to the observed phenotype.

### Pharmacological inhibition of fibroblast PIK3Cδ reduces BC tumor growth *in vivo*

Next, we used an orthotopic BC xenograft model where MDA-MB-231 or MDA-MB-231 with MRC5 were co-injected in the mammary fat pads of NOD SCID mice in order to examine the effects of PIK3Cδ inhibition. After tumor formation and mice randomization, perioral (PO) administration of CAL-101 or vehicle (30% PEG 400 +0.5% Tween 80) was initiated for both groups according to the plan depicted in **Figure 5A**.

**Figure 5:**
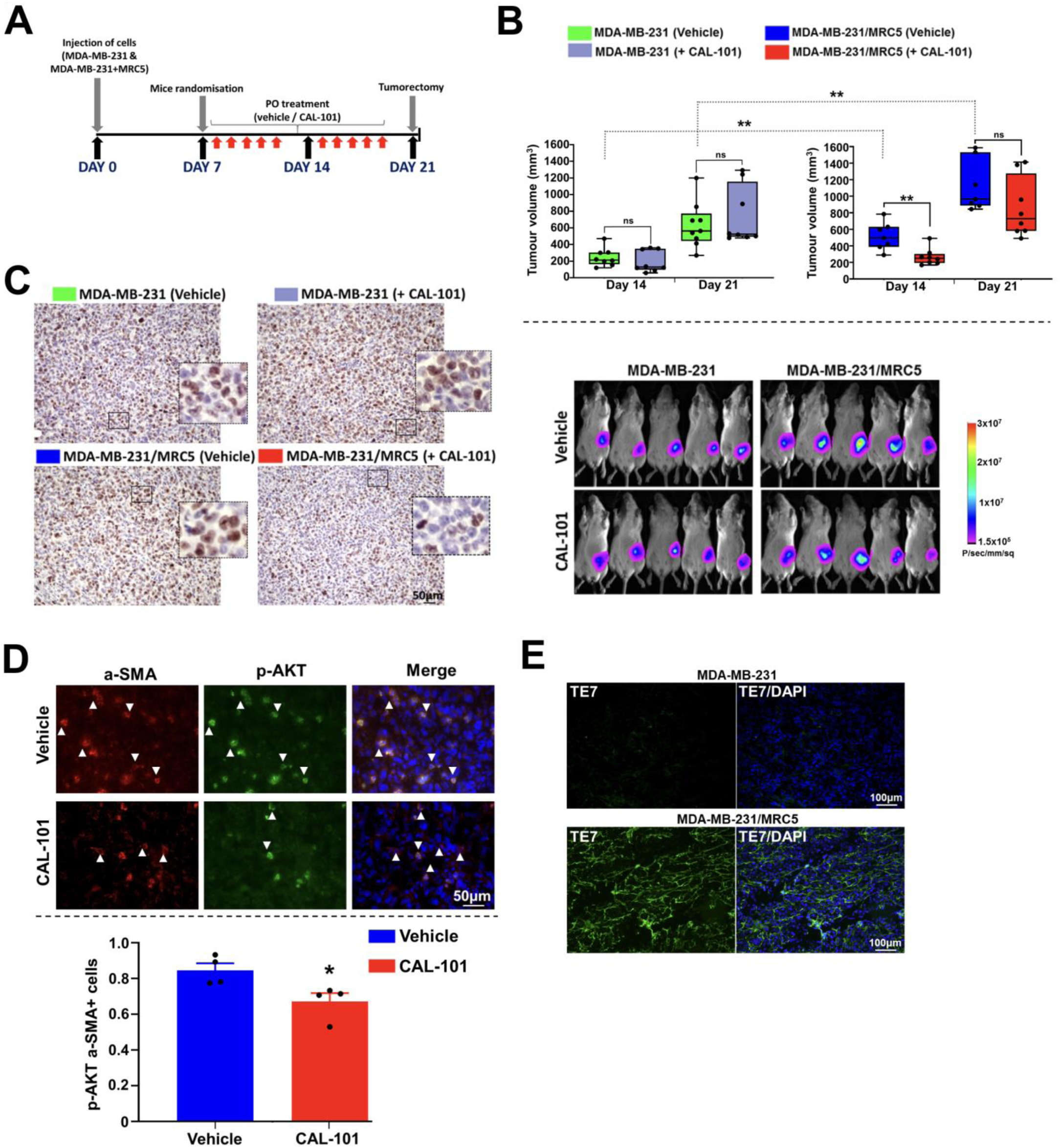
Effects of fibroblast-PIK3Cδ inhibition on TNBC tumor growth *in vivo*. (**A**) Schematic representation of the *in vivo* experiment using NOD CB17 PRKDC/J mice. MDA-MB-231 (Groups 1-2) and MDA-MB-231/MRC5 (Groups 3-4) tumor cells were implanted s.c. on day 0. After randomization on day 7, treatment with CAL-101 was initiated in Groups 2 and 4, whereas Groups 1 and 3 were administrated with vehicle. During the course of the study, the growth of the subcutaneously implanted primary tumors was determined twice weekly by luminescence and caliper measurement. (**B**) Upper panel: Box and whisker plots comparing different groups at Day14 and Day 21. Significance was calculated using unpaired t-test. Results are expressed as mean ± SEM; ** P < 0.01, ns: not significant. Lower panel: Representative *in vivo* images of different groups, treated with vehicle or CAL-101. (**D**) Histological analysis of Ki67 expression in representative tumor tissue sections of different groups. Original magnification, 20×. Scale bar, 50μm. (**D**) Representative images of immunofluorescent staining of MDA-MB-231/MRC5 tumor cryosections for α-SMA and p-AKT (Ser473) after vehicle or CAL-101 treatment. Significance was calculated using unpaired t-test. Results are expressed as mean ± SEM; * P < 0.05 vs. vehicle treated tumors. Original magnification, 40×. Scale bar, 50 μm. (**E**) Representative images of immunofluorescence staining of tumor cryosections using TE-7 anti-human fibroblast antibody. Original magnification, 20×. Scale bar, 100 μm.

Consistent with previous reports (46), cancer cells in the co-injected tumors (MDA-MB-231+MRC5) exhibited larger tumor volumes (**Figure 5B**) and a higher proliferative rate based on Ki-67 labelling (**Figure 5C**). Moreover, CAL-101 treatment of MDA-MB-231 tumors did not significantly affect their *in vivo* growth (**Figure 5B**), in agreement with our cell-based data that demonstrated only a marginal inhibitory effect of CAL-101 on MDA-MB-231 proliferation (**Supplementary Figure 16**). Interestingly, MDA-MB-231+MRC5 tumors were reduced following treatment with CAL-101 (**Figure 5B**; Day 14: 48.28% average reduction; Day 21: 23.65% average reduction). The efficacy of CAL-101 on PIK3Cδ activity was validated by assessing the expression of pAKT via immunohistochemistry analysis of tissue samples (**Figure 5D**). Moreover, the presence of 14human fibroblasts in MDA-MB-231+MRC5 tumors was confirmed in cryosection slides using a human anti-fibroblast antibody (**Figure 5E**). We also checked for CD68^+^ (monocytes / macrophages) cells, however we only detected a small population of these tumor infiltrating cells, which is consistent with the immunocompromised background of this mouse strain (**Supplementary Figure 17**). Finally, animals treated with CAL-101 did not display any significant changes in weight nor gross phenotypic changes (eg. mortality) indicating that that the treatment did not cause any adverse or toxic effects (data not shown).

As a proof of principal study of the potential use of PIK3Cδ inhibitors for treatment of BC, we also employed the MMTV-PyMT transgenic model (47), which is driven by the mammary gland-specific expression of polyoma middle T antigen under the control of mouse mammary tumor virus promoter/enhancer. The MMTV-PyMT mice received daily oral administration of either control vehicle or CAL-101 (10 mg/kg) for a period of six weeks. Our results revealed a significant reduction in tumor growth following CAL-101 treatment (**Figures 6A, 6B and Supplementary Figures 18A, 18B**). In addition, the number of lung metastasis nodules was significantly reduced in the CAL-101 group, compared with the control group, as evidenced by H&E staining and macroscopic observation of lung specimens (**Figure 6C and Supplementary Figures 18B and 18C**). Furthermore, following CAL-101 treatment, the expression of p-AKT was significantly decreased in tumor infiltrating fibroblasts (a-SMA^+^) (**Figures 6D**), besides -as expected-in macrophages (F4/80^+^) (**Figure 6E**), while no changes were observed in the total PIK3Cδ levels of fibroblasts or macrophages (**Supplementary Figures 18D and 18E**). In addition, as demonstrated in the co-cultures between immortalized TNBC and fibroblast cell lines (**Supplementary Figure 4C**), PIK3Cδ was exclusively expressed in CAFs isolated from MMTV-PyMT tumors, while cancer cells had low/undetectable levels of PIK3Cδ, the expression of which did not change following co-culturing with CM isolated from CAFs (**Supplementary Figure 18F**), further supporting the *in vitro* evidence that the effects of CAL-101 are cancer cells-independent. Finally, IHC analysis of MMTV tumors revealed an increased expression of NR4A1 as well as PLGF and BDNF following CAL-101 treatment (**Figure 6F**) supporting our cell-based data. Taken together, these results support the involvement of fibroblast-expressed PIK3Cδ in promoting BC growth *in vivo*.

**Figure 6:**
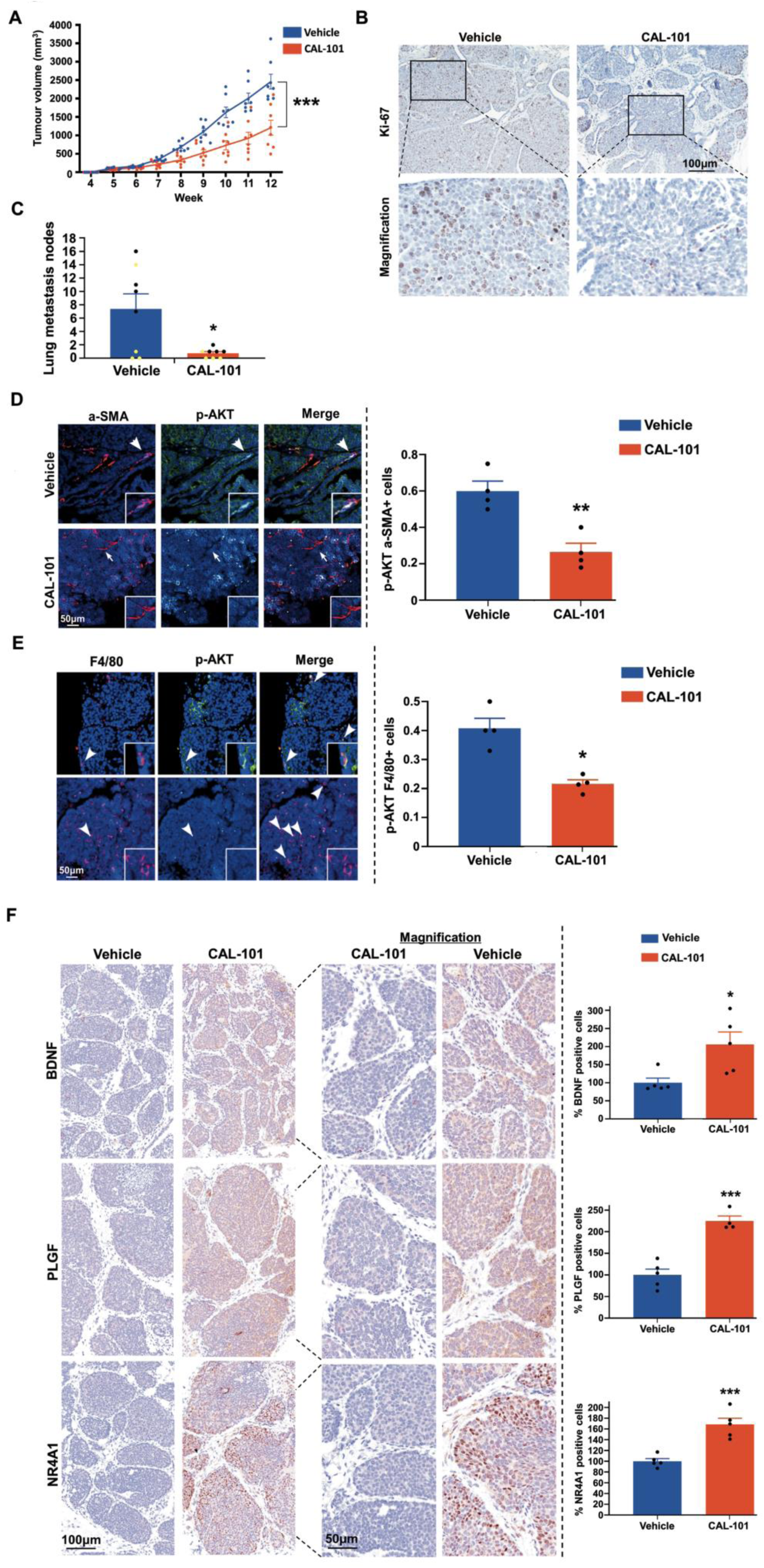
Effects of CAL-101 treatment on tumor growth of MMTV-PyMT transgenic mice. (**A**) Tumor volumes chart from MMTV-PyMT transgenic mice after vehicle or CAL-101 treatment (n=8 mice / group). Individual values for each mouse are displayed. Significance was calculated using unpaired t-test (week 12). Results are expressed as mean ± SEM; *** P < 0.001. **(B)** Representative images for immunohistochemical Ki-67 staining in the mammary tumor sections of MMTV-PyMT transgenic mice after vehicle or CAL-101 treatment. **(C)** Quantification of lung metastatic nodules in each group. Significance was calculated using unpaired t-test. Results are expressed as mean ± SEM; * P < 0.05. Yellow and black dots represent mice that were sacrificed at week 12 or week 15 respectively. **(D)** Left panel: Representative images of immunofluorescent staining for α-SMA and p-AKT (Thr308) in the mammary tumor sections of MMTV-PyMT transgenic mice after vehicle or CAL-101 treatment. Arrows indicate a-SMA^+^ fibroblasts. Higher-magnification images are shown at the bottom right corner. Right panel: Quantification of p-AKT (Thr308) immunofluorescent staining in tumor infiltrating a-SMA^+^ fibroblasts in the mammary tumors of MMTV-PyMT transgenic mice after vehicle or CAL-101 treatment. Significance was calculated using multiple t-tests. Results are expressed as mean ± SEM; ** P < 0.01 vs. vehicle treated tumors. (**E**) Left panel: Representative images of immunofluorescent staining for F4/80 and p-AKT (Thr308) in the mammary tumor sections of MMTV-PyMT transgenic mice after vehicle or CAL-101 treatment. Arrows indicate F4/80^+^ macrophages. Higher-magnification images are shown at the bottom right corner. Right panel: Quantification of p-AKT (Thr308) immunofluorescent staining in tumor infiltrating F4/80^+^ macrophages in the mammary tumors of MMTV-PyMT transgenic mice after vehicle or CAL-101 treatment. Significance was calculated using multiple t-tests. Results are expressed as mean ± SEM; * P < 0.05 vs. vehicle treated tumors. (**F**) Representative images of IHC staining of the mammary tumor sections of MMTV-PyMT transgenic mice stained for BDNF, PLGF and NR4A1, after vehicle or CAL-101 treatment. Significance was calculated using unpaired t-test. Results are expressed as mean ± SEM; * P < 0.05, *** P < 0.001 vs. vehicle treated tumors. Original magnification, 20×. Scale bar, 100 μm. The intensity of BDNF, PLGF and NR4A1 signals were evaluated in the vehicle treated tumor slides and then this value was compared with the signals from CAL-101 treated tumor slides results are expressed as percentage (%) of signal vs vehicle treated tumors.

### High fibroblast PIK3Cδ expression correlates with poor patient outcome in TNBC

Finally, we investigated the plausible role of fibroblast (a-SMA^+^) or tumor-expressed (epithelial cancer cells) PIK3Cδ in a clinical setting by analyzing a well-characterized TNBC patients’ cohort (n=179) (48, 49). The clinico-pathologic parameters are summarized in **Supplementary Table 7**. PIK3Cδ expression was variable in both tumor (H-score range 20-220) and surrounding cancer associated fibroblasts (5-100%) with high PIK3Cδ tumoral expression (H-Score >130 observed in 31/179 cases (17%) whereas fibroblasts showed high PIK3Cδ (>85%) in 44/179 (25%) of the cases (**Figure 7A and Supplementary Figure 19**). Analysis of the surrounding stromal (a-SMA^+^) PIK3Cδ expression, revealed PIK3Cδ as an independent prognostic factor for overall survival (OS; P=. 0.000285; **Figures 7B**) and disease free survival (DFS; P = 0.048; **Figure 7C**), indicative of fibroblast-PIK3Cδ’s involvement in TNBC progression. Furthermore, multivariate analyses demonstrated that PIK3Cδ in the surrounding cancer fibroblasts was also an independent prognostic factor for OS (P = 0.001), DFS (P = 0.044) as well as for age and nodal stage (**Supplementary Table 8**). Similar analyses revealed that, high PIK3Cδ protein levels in tumoral-PIK3Cδ were associated with a significantly shorter OS (P = 0.0004) and DFS (P = 0.009) (**Supplementary Figures 20A and 20B**), while tumoral-PIK3Cδ was an independent prognostic factor of age, nodal stage and lymphovascular invasion (LVI) status and for predicting OS (P = 0.006) and DFS (P = 0.028) (**Supplementary Table 9**). Interestingly, investigation of fibroblast-PIK3Cδ in an ERα^+^ patients’ cohort from Singapore (n=73; (P = 0.703) did not reveal any correlation with survival outcome (**Figure 7D and Supplementary Tables 10-12**).

**Figure 7:**
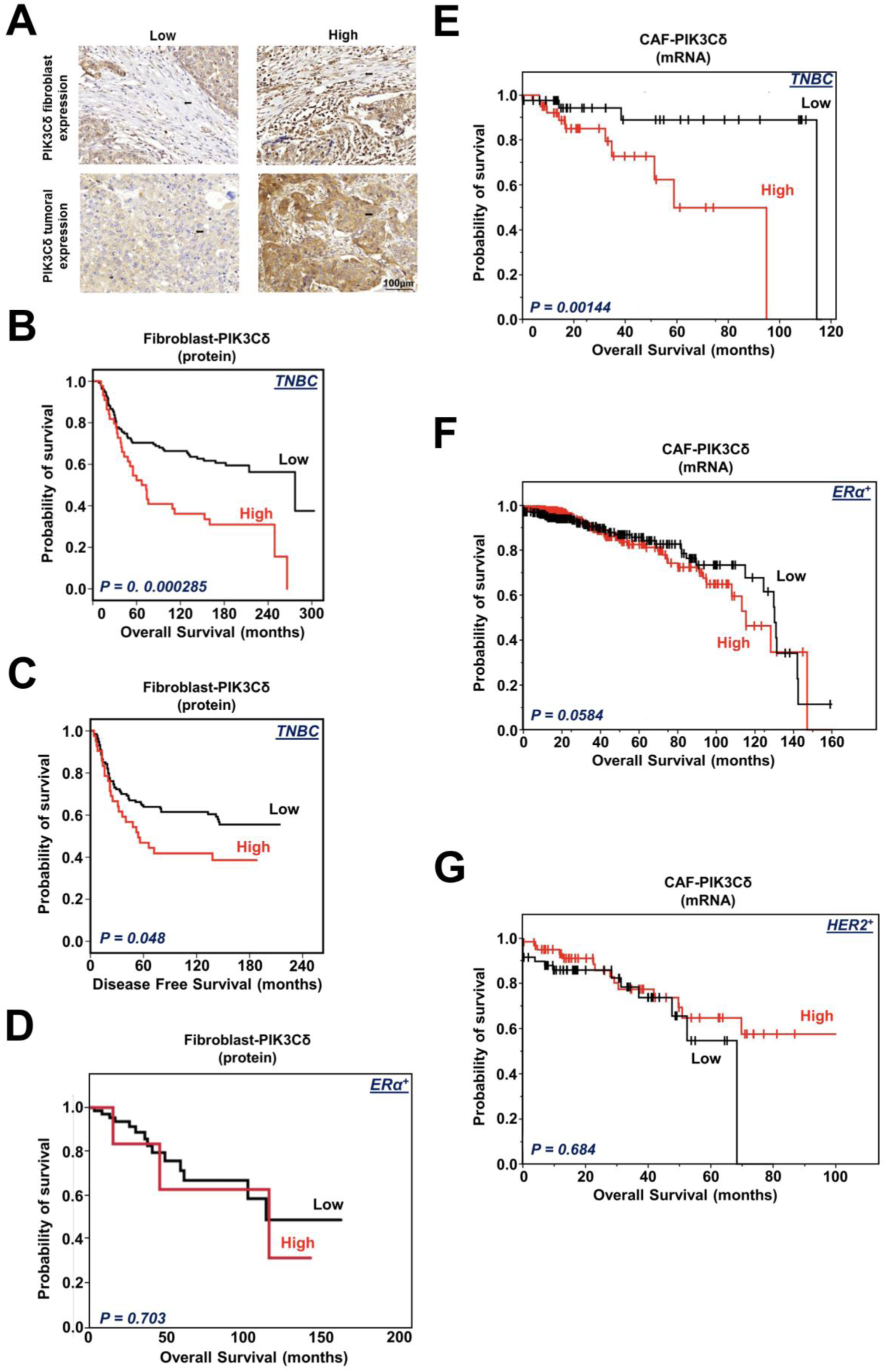
PIK3Cδ expression in fibroblast cells and association with patients’ survival. (**A**) Representative images of low and high PIK3Cδ expression in tumor or surrounding fibroblast cells (a-SMA^+^) are shown. Original magnification, × 20. Scale bar, 100 μm. (**B**) Kaplan-Meier plots showing the association between fibroblast-PIK3Cδ protein expression with OS (P = 0.000285) in TNBC patients. (**C**) Kaplan-Meier plots showing the association between fibroblast-PIK3Cδ protein expression with DFS (P = 0.048) in TNBC patients. (**D**) Kaplan-Meier plots showing the association between fibroblast-PIK3Cδ protein expression with OS (P = 0.703) in ERα^+^patients. (**E**) Kaplan-Meier plots showing the association between CAF-PIK3Cδ mRNA expression with OS (P = 0.001) in TNBC patients following deconvolution of bulk TCGA RNA-seq samples. (**F**) Kaplan-Meier plots showing the association between CAF-PIK3Cδ mRNA expression with OS (P = 0.058) in ERα^+^ patients following deconvolution of bulk TCGA RNA-seq samples. (**G**) Kaplan-Meier plots showing the association between CAF-PIK3Cδ mRNA expression with OS (P = 0.684) in HER2^+^ patients following deconvolution of bulk TCGA RNA-seq samples. * P < 0.05, ** P < 0.01, *** P < 0.001.

We also employed another approach to investigate the potential association of CAF-PIK3Cδ mRNA levels with survival outcomes of ERα^+^, HER2^+^ and TNBC subtypes, by de-convoluting bulk RNA-seq samples from TCGA BC data (as described in M&M). As illustrated in **Figure 7E**, TNBC patients (n=108) with high CAF-expressed PIK3Cδ levels had shorter OS compared to those with low CAF-expressed PIK3Cδ (P = 0.001), in agreement with our IHC data. Conversely, when studying the bulk tumors, there was no significant association between PIK3Cδ mRNA levels and TNBC patients’ survival outcome (P = 0.405; **Supplementary Figure 20C**), opposite to the IHC data, emphasizing the discrepancies that can arise when examining mRNA vs protein levels, which can lead to different conclusions. In addition, these results also highlight the importance of comprehensively analyzing the different cell types separately within the TME. Moreover, expression of CAF-PIK3Cδ mRNA levels were not predictive for survival neither for ERα^+^ (n=778, P = 0.0584; **Figures 7F**) -recapitulating the IHC data- nor for HER2^+^ (n=160, P = 0.684; **Figures 7G**) BC subtypes, further underlining the significance of fibroblast-expressed PIK3Cδ isoform in TNBC specifically.

We also analyzed the association between the mRNA expression levels of CAF-PIK3Cα, -β, and -γ isoforms and patients’ survival for all BC subtypes. Our results revealed that in TNBC high CAF-expressed PIK3Cα (P = 0.01; **Supplementary Figures 20D**) or CAF-expressed PIK3Cβ (P = 0.01; **Supplementary Figures 20E**) were correlated with shorter OS, while there was no association of either PIK3Cα or PIK3Cβ with ERα^+^ or HER2^+^ BC patients’ survival (ERα^+^ patients: PIK3Cα, P = 0.106; PIK3Cβ, P = 0.15. / HER2^+^ patients: PIK3Cα, P = 0.731; PIK3Cβ, P = 0.849). Regarding CAF-PIK3Cγ mRNA levels, there was no correlation with survival in neither TNBC (**Supplementary Figures 18F**) nor any of the other BC subtypes (ERα^+^ patients: PIK3Cγ, P = 0.137. / HER2^+^ patients: PIK3Cγ, P = 0.943). In conclusion, our results demonstrate the clinical significance of fibroblasts-expressed PIK3Cδ in TNBC.

## Discussion

TNBC represents an aggressive BC subtype where there remains an unmet clinical need; currently the recommended therapeutic approach in the neoadjuvant, adjuvant and metastatic settings are based on chemotherapeutics (most often platinum-, anthracycline or taxane -based) with recent data suggesting roles for antibody-drug conjugates and immunotherapies (50–53). However, fewer than 30% of women with metastatic TNBC will survive 5 years after diagnosis (54). Sequencing and other ‘omics’ have revealed an unexpected level of heterogeneity in TNBCs and led to identification of potential actionable targets (52).

However, translational research and clinical trials usually focus on targeting epithelial cancer cells. This is likely to diminish the contribution of reciprocal interactions between malignant and stromal cells that creates a local microenvironment, which fosters tumor growth and also influences responses to treatment (55). Yet, the prognostic and predictive significance of gene/protein expression signatures of the surrounding stroma have been well-documented and could represent unexplored ground within the TME which could then be used to improve therapies and outcomes.

PKs are involved in almost every aspect of cell activity and perturbation of their signaling can contribute to most human diseases including malignancies (56–59). Pharmaceutical intervention targeting aberrant kinase signaling represents the major therapeutic approach but although targeted therapies against PKs have improved the clinical outcome of patients, resistance to these treatments develops (60), emphasizing the need for the identification of new druggable targets.

Surprisingly, despite extensive research describing deregulated kinase activity in cancer cells, there has been no thorough and comprehensive investigation so far about how kinases expressed in stromal cells can influence tumor growth and malignant progression. Therefore, in this study, we focused on fibroblasts, the main stromal component in the TME, whose multiplex role in BC initiation, progression and therapy-resistance has been well described (7, 13, 14). We performed a kinome siRNA screening in two different fibroblast cell lines, aiming to identify kinases responsible for stroma-tumor cross-talk. Our siRNA screening/3D co-culturing model was linked to an invasion readout assay that could be performed easier and more reliably by using TNBC cell lines, considering their invasive potential, compared to the non-invasive and less aggressive ER^+^ luminal BC cells. However, our findings do not rule out the possibility that these targets (kinases) can also be linked to other BC subtypes. Ultimately this study aimed to identify targets that are associated with an aggressive phenotype, invasion being the clearest readout. Nevertheless, the original screening is not designed to identify a mechanism of action and therefore the target’s effects could be diverse once studied further (e.g. in an *in vivo* setting).

Considering the limited available therapeutic options for TNBC, we focused on this subtype since the identification of new putative druggable targets for TNBC is fundamental. Amongst a subset of differently fibroblast-expressed kinases that were able to modulate TNBC progression (positively or negatively), PIK3Cδ was one of the prominent hits. Despite the involvement of PI3K activity in tumor-stroma interactions (15), still the possibility of using PI3K inhibitors on fibroblasts has not been considered to date.

Given its almost exclusive expression in fibroblasts, this target (PIK3Cδ) could not have been identified by focusing solely on TNBC cells, further supporting that the contribution of the TME in cancer development and progression needs to be studied in detail. Using a variety of 2D and 3D co-culturing models we determined that fibroblast-expressed PIK3Cδ is integral in TNBC progression. We validated our findings using genomic approaches (loss and gain of function experiments) and we also assessed the effects of the chemical inhibition of PIK3Cδ, using a highly selective FDA approved PIK3Cδ inhibitor (CAL-101; Idelalisib) (61), confirming that the catalytic activity of fibroblast PIK3Cδ is required for its paracrine effects on TNBC cells. Mechanistically, using an integrated analysis of the fibroblast PIK3Cδ-regulated secretome and its paracrine mediated transcriptomic changes in TNBC cells, we identified secreted factors and genes that represent key signaling pathways contributing towards the observed PIK3Cδ-induced tumor promoting phenotype. We focused our attention on the link between the overexpression of fibroblast-secreted factors, including PLGF and BDNF, and the upregulation of NR4A1 transcription factor in TNBC epithelial cells, after inhibition of fibroblast-PIK3Cδ. Intriguingly, NR4A1 has been recently reported to act as a tumor suppressor implicated in TNBC proliferation, viability, migration and invasion (37). Our results herein support a model where inhibition of fibroblast-expressed PIK3Cδ impedes TNBC progression, by promoting the secretion of PLGF, BDNF and other factors, which in turn lead to the paracrine upregulation of NR4A1 in TNBC cells (**Figure 4I**). The existence and elucidation of additional direct as well as reciprocal signaling pathways originated from cancer cells towards fibroblasts, which could potentially affect PIK3Cδ expression and ultimately contribute to this phenotype, merits further investigation.

To examine the effects of fibroblast-expressed PIK3Cδ on TNBC growth *in vivo*, we initially attempted to generate stable PIK3Cδ knock-out (KO) HMF and MRC5 cell lines and compare their involvement in tumor growth vs PIK3Cδ wild-type fibroblasts. However, PIK3Cδ-KO clones exhibited a relative slow growing rate and considering also the fact that fibroblasts can easily differentiate (62), led us to the alternative option of pharmacologically (CAL-101) inhibiting PIK3Cδ. CAL-101 had no effect when used as a treatment on MDA-MB-231 tumor growth, similarly to what we observed in our *in vitro* (cell-based) proliferation data when MDA-MB-231 cells were directly treated with CAL-101. Moreover, CAL-101 was administered orally in immune-deficient mice which, combined with the lack of any MDA-MB-231 tumor inhibition, suggests that there were systemic responses initiated from other sub-populations of cells within the TME. This result, along with a recent report, where the authors showed that pharmacological inhibition of PIK3Cδ impedes *in vivo* tumor growth by targeting cancer cells and macrophages, further supports the stromal involvement of PIK3Cδ in BC and the potential use of PIK3Cδ inhibitors in a clinical setting.

In our immunocompromised xenograft model, we initially verified that the co-injection of fibroblasts (MDA-MB-231+MRC5) had an additive effect in tumor formation, when compared to MDA-MB-231 cancer cells alone. More importantly, we observed a decrease in MDA-MB-231+MRC5 tumors following daily treatment with CAL-101. Considering that the only variable between the two mouse models was the introduction of fibroblasts, it is clear that the anti-tumor effects of CAL-101 were conferred via their actions on fibroblasts. Noteworthy, the tumor growth reduction that was observed on Day 21 between MDA-MB-231+MRC5 CAL-101-treated tumors vs MDA-MB-231+MRC5 vehicle-treated ones was borderline non-significant despite the 23.65% median reduction (it is worth mentioning that alternative statistical tests gave a significant P value). We attributed this to the progressive population dilution and decreased viability of human fibroblast cells (MRC5) as the tumor size increases, causing a reduction in relative potency of CAL-101 (as the dosage was left unchanged) on fibroblast PIK3Cδ and its paracrine consequences. Moreover, our results in MMTV-PyMT transgenic mice also revealed a significant reduction in primary tumor growth as well as in metastasis following treatment with CAL-101. The downregulation of PIK3Cδ’s activity in fibroblasts, apart from macrophages, implies of a prospective additive, immune-independent mechanism of action of PIK3Cδ inhibitors for cancer treatment. In addition, as fibroblasts have been reported to influence a number of other immune cells, namely monocytes and macrophages (63, 64), the existence of additional PIK3Cδ-mediated paracrine signaling effects between different cell types could not be ruled out.

The translational significance of fibroblast-expressed PIK3Cδ was validated in a TNBC cohort, where we revealed PIK3Cδ as a prognostic factor for outcomes (OS and DFS), providing strong evidence for the use of PIK3Cδ inhibitors in this setting in clinical trials. Interestingly, PIK3Cδ was also expressed in the cancer cell population of patients, possibly as a result of inflammatory processes, since it has been reported that PIK3Cδ can be activated by pro-inflammatory mediators (65). This can explain the low/undetectable protein levels of PIK3Cδ in our tested BC cell lines as well as in our animal models, considering the short period of the *in vivo* experiments. In light of new evidence of the existence of distinct TNBC subtypes (66, 67), an even more comprehensive profiling of TNBC patients can reveal a specific subgroup where stromal/tumoral PIK3Cδ can epitomize a successful treatment strategy.

In conclusion, this study highlights the so-far undescribed tumor promoting role of fibroblast-expressed PIK3Cδ in BC. Although our work predominantly focused on TNBC, fibroblasts represent the major cellular components within the TME in most cancers, therefore the involvement of PIK3Cδ in other BC subtypes and malignancies should be explored. Moreover, considering that local invasion and metastasis are the main causes of death for most types of cancer (including BC), this discovery also opens new potential therapeutic paths and supports the rationale for using PIK3Cδ inhibitors (as single or combined therapy) for the treatment of solid tumors where irregular activation of stromal PIK3Cδ occurs independently of the immunological landscape.

## Acknowledgments

We would like to Margarita Andreadou for her help with the histology analysis. We would also like to thank the Nottingham Health Science Biobank for the provision of tissue samples. The authors also wish to acknowledge the role of the Breast Cancer Now Tissue Bank in collecting and making available the samples used in the generation of this publication. We thank Michael Toss for his invaluable assistance with histopathology and Rosy Favicchio for the useful and stimulating discussions. We would like to thank Enrico Frabetti for his endless patience and support.

## Materials and methods

### Reagents

CAL101 (Idelalisib), AS252424, PI-103 and Leniolisib, were purchased from Selleckem and resuspended in DMSO (Sigma). PLGF (#100-06) and BDNF (#450-02) were purchased from Peprotech. Cytosporone B (#C2997) was purchased by Sigma and resuspended in DMSO. PIK3Cδ (#34050), AURKA (#14475), AKT total (#4691), AKT-Ser473 (#4060) and AKT-Thr308 (#13038), total PDPK1(#3062), Integrin β1 (#9699), S100A4, (#13018) PDGFR (#28E1), Caveolin1 (#3267), antibodies were purchased from Cell Signaling; NR4A1 (#STJ94578), mTOR total (#STJ113771) and mTOR-Ser2448 (# STJ27512) and PDPK1-Ser241 (# STJ29323) were purchased from St Johns Laboratory. FAP (#ab54651), α-SMA #ab5694 (α-smooth muscle actin) were purchased from Abcam. PIK3Cδ overexpressing plasmid (pCMV6-PIK3Cδ-AC-GFP) and the empty vector (pCMV6-AC-GFP) were purchased from Origene. All other reagents were purchase from Thermofisher Scientific.

### Cell lines and primary samples

MDA-MB-231 were purchased by ATCC, maintained in DMEM low glucose supplemented with 10% FBS and 10 u/ml Penicillin/Streptomycin, incubated at 37 °C with 5% CO_2_. BT-549 were purchased by ATCC, maintained in RPMI 1640 supplemented with 10% FBS and 10 u/ml Penicillin/Streptomycin, incubated at 37 °C with 5% CO_2_. HMF (Human adult Mammary Fibroblast) were purchased from ZenBio, maintained in E-MEM supplemented with 10% FBS and 10 u/ml Penicillin/Streptomycin and 30 ng/ml of EGF, incubated at 37 °C with 5% CO_2_. MRC5 were purchased from ATCC, maintained in E-MEM supplemented with 10% FBS and 10 u/ml Penicillin/Streptomycin and 30 ng/ml of EGF, incubated at 37 °C with 5% CO_2_. Matching primary fibroblasts from tumor (< 5cm) or from a distal site (> 5cm) of TNBC patients, were kindly provided by Breast Cancer Now tissue-bank at Bart’s Cancer Institute, Queen Mary University, London (patients’ information are provided in **Supplementary Table 8**). Primary fibroblasts were grown in DMEM/F12 medium, supplemented with 10μg/ml apo-transferrin, 5μg/ml insulin, 30 ng/ml of EGF, 0.5μg/ml hydrocortisone, 5 µg/ml Amphotericin B, 10% FBS and 10 u/ml Penicillin/Streptomycin.

### siRNA transfection

HMF and MRC5 fibroblasts were transfected with a pool of 3 siRNAs targeting each of the 710 human kinases (Silencer Human Kinase siRNA Library, #A30079 Thermofisher). Fibroblasts (1250 cells/well) were seeded in a 96 well plate. 50 nM siRNA/well were used for transfection using 0.8 µl/well of Hi-Perfect (Qiagen) following manufacturer’s instructions. For NR4A1 silencing, the following siRNAs were used: #s144919 and #s41752 (Thermofisher). In addition, non-targeting scrambled control (#AM4611, Thermofisher) was used. Briefly, 400.000 MDA=MB-231 cells were resuspended in 20 µl of complete buffer SE with the addition of 300 nM siRNA. Silencing was then confirmed by real-time qRT-PCR. For silencing, the following siRNAs were used: PLGF #s10399 and BDNF #s1964 and #s1963 (Thermofisher). In addition, non-targeting scrambled control (#AM4611, Thermofisher) was used. Briefly, 400.000 MRC5 cells were resuspended in 20 µl of complete buffer SE with the addition of 300 nM siRNA, and electroporated using 4D-Nucleofector following manufacturer instruction.

### 3D co-culture and invasion assay

24 hours after transfection, 2 µl of 0.025% trypsin in 7µl of PBS-EDTA was added to each of the fibroblasts transfected wells. Trypsin action was stopped by adding 30 µl of medium containing 1250 MDA-MB231 cells. Cell suspension was then transferred to an Ultra-low attachment 96 well plate (corning). Plates were centrifuge for 3 minutes at 300xg and incubated at 37 °C for 72 hours. After spheroids formation, 30 µl of Growth factor reduced Matrigel was added to each well, plates were incubated for 2 hours at 37 °C, followed by adding 20% FBS medium to each well to promote invasion. Spheroids pictures were taken after 3 and 6 days using EVOS FL microscope (Thermofisher) and the images were analyzed with ImageJ software. The results were expressed as changes in the overall spheroid surface (Δ=Surface day 6 - Surface day 3).

The Δ-value of each silenced kinase (Δ_K_) was compared with the Δ-value of the control (Δ_CT_ of the non-targeting/scrambled siRNA), at different time-points, to obtain a Δ_Ratio_ (Δ_Ratio_=Δ_CT_/Δ_K_). Kinases with a Δ_Ratio_ ≤ 0.5 (50% less invasion vs CT) were considered as ‘invasion promoting kinases’, (as their silencing led to a reduction of invasion), while kinases with a Δ_Ratio_ > 2 (100% more invasion vs CT) were considered as ‘invasion inhibiting kinases’, (as their silencing led to an increase of invasion). Based on our cut-off values/criteria, we classified kinases with Δ_Ratio_ between 0.5 and 2 as ‘uninfluential on the invasiveness of MDA-MB-231 cells. The Δ_Ratio_ values were used to calculate the Z-Scores based on the formula: z = (x – μ)/σ (μ: Δ_Ratio_ mean of 710 kinases; σ: standard deviation (SD); x: Δ_Ratio_ value for each kinase).

### Screening validation

MRC5 cells (50.000/well) were seeded in a 24well plate. Using Fugene HD (Promega), cells were transfected with 0.5 µg/well of pLKO.1-puro Vector empty or with plasmids containing the target shRNA sequence for the indicated kinases (Mission Library, Sigma). 48h after transfection, cells were counted for 3D co-culture with MDA-MB-231 and subsequent invasion assays were performed as described above. Part of the transfected cells were used to determine the efficacy of silencing using Cells To CT kit (Thermofisher), following manufacturer’s instruction for Real-time PCR assay (qRT-PCR), using specific primers for each silenced kinase (QuantiTect, #249900, Qiagen).

### Lipophilic tracer staining and Real-time 3D invasion assay

SP-DiOC18(3) or Dil Stain (Thermofisher) were added to 10^6^ cells in serum-free medium to a final concertation of 5 µM. Cells were then incubated at 37°C for 1h, and centrifuged for 5 minutes at 300xg and washed twice with PBS. Finally, cells were resuspended in fresh complete medium and used for further assays.

### Western Blotting

Cells were dissolved in RIPA Buffer (Sigma), as previously described (68). Protein concentration was measured by BCA Protein Assay Reagent Kit (Pierce, Rockford, IL, USA), as described (69). Proteins were fractionated on 10% SDS-PAGE, and transferred by electrophoresis to Nitrocellulose Transfer Membrane (GE). Membranes were incubated with several primary antibodies. Horseradish peroxidase-conjugated antibody anti-mouse IgG and anti-rabbit IgG (Dako) were used to detect immunoreactive bands and binding was revealed using enhanced chemiluminescence (Pierce). The blots were then stripped and used for further blotting with anti-GAPDH antibody (Thermofisher #AM4300). Unless otherwise specified, displayed western blots are representative images of at least 2 independent experiments. Where indicated densitometric analysis of detected bands were performed by measuring the peak height of the bands using ImageJ and normalized using loading control.

### 2D Invasion Assay

Reduced growth factors Matrigel (BD Biosciences) was polymerized in 8 µm pore cell inserts (Sarstedt) prior to the addition of cells. MDA-MD-231 or BT549 (5×10^4^) cells were seeded into the insert containing Matrigel in serum free media. Fibroblast, pre-treated with CAL101 or DMSO for 24h were used as an attractant in the bottom chamber, and cells were allowed to migrate for additional 24 h, the inserts were then removed and migrated cells were fixed with PFA 4% and then stained with crystal violet and counted providing a quantitative value for migrated cells across the membrane.

### Real-time invasion assay

MDA-MB-231 and BT-549 TNBC cell lines were seeded in cell inserts with 0.4 µm pores, to be co-cultured with fibroblasts pre-treated with 10 µM CAL-101 or vehicle only. 24h after co-culture, TNBCs were collected and counted and used for real-time invasion assay. Cell invasion was monitored in real time using the xCELLigence RTCA technology as previously described (70). CIM-16-well plates, provided with interdigitated gold microelectrodes on bottom side of a filter membrane interposed between a lower and an upper compartment, were used. The lower chamber was filled with 10 % serum-medium, cells (2×10^4^ cells/well) were seeded on filters in serum-free medium. Microelectrodes detect impedance changes which are proportional to the number of invading cells and are expressed as cell index. Invasion was monitored in real-time for at least 30h. Each experiment was performed at least twice in quadruplicate.

### Conditioned medium preparation

HMF and MRC5 were seeded in 10 cm plates and treated with either vehicle or with 10 µM CAL-101 or transfected to silence or overexpress PIK3Cδ, in serum free for 24h in order to obtain the conditioned medium (CM). Medium collected was centrifuged for 30’ at 1800 x g at 4°C and then passed through a 0.22µm filter, aliquoted and store at -80°C for further assays.

### PIK3Cδ overexpression and silencing

MRC5 cells were transfected with pCMV-PIK3Cδ-AC-GFP (Origene) or empty vector using 4-D electroporator (LONZA), following manufacturer instruction. Briefly, 400.000 cells were resuspended in 20 µl of complete buffer SE (Lonza), and 0.4 µg of plasmid DNA were added prior electroporation, cells were then seeded in warm complete medium. Similarly, for PIK3Cδ siRNAs (s10531, s224263, s10530 Thermofisher) or scramble control (AM4611, Thermofisher), 400.000 MRC5 were resuspended in 20 µl of complete buffer SE with the addition on 300 nM siRNA. For short hairpin RNA (shRNA) silencing, pLKO.1-puro Vector empty or containing the target shRNA sequence for PIKCδ (Mission Library, Sigma) were transfected using 4-D electroporator (LONZA), following manufacturer instruction. Briefly, 400.000 cells were resuspended in 20 µl of complete buffer SE (Lonza), and 0.4 µg of plasmid DNA were added prior electroporation, cells were then seeded in warm complete medium. Overexpression and silencing were then confirmed by real-time qRT-PCR.

### RNA extraction and Real-time qRT-PCR

Total RNA was extracted from the cell lines using PureLink RNA Mini Kit following the manufacturer’s instructions (Invitrogen). All RNA samples were subjected to DNase I treatment. The RNA samples were subjected to RT using SuperScript IV VILO following the manufacturer’s instructions (Invitrogen). Real-time quantitative RT-PCR (qRT-PCR) was performed to assess the expression of human PIK3CD (Hs00192399_m1) using the TaqMan gene expression assay (Applied Biosystems) while GAPDH was used as reference (Hs99999905_m1). For RNAseq validation, Real-time PCR assay (qRT-PCR), was performed using specific primers for each silenced kinase (**Supplementary Table 4**). The samples were run in triplicate on an Applied Biosystems STEPONE thermal cycler and analyzed with the SDS 1.9 Software (Applied Biosystems), by applying the method described by Pfaffl and previously applied (71, 72).

### Cell viability

Cell viability assay was performed as previously described (69) (36). Briefly, cells were seeded in 5.000 for well in a 96 well plate and treated with the indicated compound and time points and cell viability was evaluated by Cell Titer-Glo 2.0 (Promega) according to manufacturer instruction. 6 replicates in 3 independent experiments were analyzed and results were expressed as percentage mean ± SEM vs vehicle treated cells.

### Statistical analysis for cell based assay

Statistical analysis was performed using GraphPad Prism software. Unless otherwise specified, all the graphs displayed represent a distribution of all recorded measures in the different experiments normalized for the control(s).

### Secretome analysis

Supernatants of CAL-101 and Vehicle (DMSO)-treated fibroblasts were collected after 24 h incubation in serum free MEM. Supernatants were dialyzed and after quantification, secreted proteins were biotin-labelled and hybridized onto human L1000 glass slide arrays (RayBiotech; https://www.raybiotech.com/human-l-1000-array-glass-slide-2). Detected fluorescence signals were then normalized to positive controls.

### RNA Extraction, Library Construction and RNA-Sequencing

Cell pellets were obtained from MDA-MB-231 co-cultured in 6-well cell inserts with MRC5 treated with CAL-101 10 µM or vehicle only. All conditions were analyzed using 2 biological replicates. RNA was extracted as described before (36). The RNA samples were then quantified using a Qubit 2.0 (Life Technologies, CA, USA) fluorimeter from Invitrogen with a high sense RNA kit. The quality of RNA was assessed and confirmed using the RNA Pico 6000 kit. The high-quality RNA were then used for library preparation using the RNA hyper prep kit with riboerase from the KAPA Biosystems (Cat. No. KK8560) according to the manufacturer’s recommendation and were fragmented at 85°C for 4.5 minutes. The libraries were amplified using 8 cycles of PCR and were ligated with the NEXTflex-96 DNA barcodes (Bio Scientific). The library was quantified using the Qubit DNA high sense kit, and its quality was confirmed using Bioanalyzer (DNA high sense kit). Libraries were then pooled and diluted to a final concentration of 2nM. Pooled and diluted library were denatured along with 1% PhiX spike in as per recommendation from Illumina. Library was loaded onto the NextSeq 550 v2 mid-output 150 cycle kit.

### Raw data processing, alignment analysis, and identification of differentially expressed genes

We first obtained the high-quality clean reads by trimming the adapter sequences and removing reads that contained poly-N or were of low-quality from the raw data using the fastX tool kit (v 0.0.14) (http://hannonlab.cshl.edu/fastx_toolkit/license.html). The quality of reads were then confirmed using the fastqc tool kit (v 0.11.5) (http://www.bioinformatics.babraham.ac.uk/projects/fastqc/). The downstream analyzes were conducted using this high-quality clean read. Briefly, the high-quality reads were mapped to the ENSEMBL built GRCH37 using the STAR aligner (v2.5.3a) (73) with the ENCODE options as described in the STAR manual. We then estimated the read counts for each gene using the summarize Overlaps function provided with the R package DESeq2 (74). Consequently, we performed differentially expressed genes (DEGs) using DESeq2 non-interaction model and the Benjamini-Hochberg corrected *P* value (*P*_adj_ ≤ 0.05) and Log2 fold change ≥ |0.5| were used as the threshold to screen significance of DEGs. The pathway analyzes for the secretome data and the RNAseq data combined were performed using the Ingenuity Pathway (https://www.qiagenbio-informatics.com/products/ingenuity-pathway-analysis).

### Functional annotation of secreted proteins

Analyzes to identify canonical pathways effected by CAL-101 treatment of the HMF and MRC5 cells was conducted by using secreted proteins differentially expressed in comparison to CAL-101 treatment and the Ingenuity Pathway Analysis tool (https://www.qiagenbio-informatics.com/products/ingenuity-pathway-analysis).

### Xenograft mouse model

Animals were obtained from Charles Rivers (Italy). All mice were kept in the animal facilities of BSRC “Alexander Fleming” under specific pathogen-free conditions. The experiment was performed with mice aged 8 weeks. All procedures were approved by the Prefecture of Attica (license #3554/2018) in accordance to national legislation and the European Union Directive 63/2010. The study consisted of 4 experimental groups each containing 10 female NOD SCID mice after randomization. On day 0, 1×10^6^ MDA-MB-231-Luc cells or 1×10^6^ MDA-MB-231-Luc with 1.5×10^6^ MRC5 cells, respectively, in 200μl Matrigel:PBS 1:1 were injected in the mammary fat pad of 20 animals per condition. Primary tumor sizes were measured twice weekly (Monday and Friday) by calipering. Tumor volumes were measured every 3 or 4 days using a calliper and calculated using the formula: W2xL/2 (L= length and W= the perpendicular width of the tumor. On day 7, after mean tumor volumes had reached approx. 100-200 mm^3^, animals were randomized into 4 groups per condition, each containing 8 animals. On the same day, treatment with 2 mg/kg CAL-101 orally was initiated for Groups 2 (MDA-MB-231+CAL-101) and 4 (MDA-MB-231+MRC5 +CAL-101), whereas animals of Groups 1 (MDA-MB-231) and 3 (MDA-MB-231+MRC5) were kept untreated (vehicle; 30% PEG 400 +0.5% Tween 80). Animals of Groups 2 and 4 were treated for 10days. Mice were sacrificed on day 28. Primary tumor tissues of all animals were photographed and collected and wet weights and volumes determined accordingly. Fifteen minutes prior to bioluminescent imaging, all mice were intraperitoneally injected (150 mg/kg) with a 30 mg/ml solution of D-luciferin (Vivo Glo Luciferin, P1043, Promega) in PBS. Quantitative imaging of bioluminescence was performed, using the In-Vivo Xtreme Imaging System (Bruker).

### Histology and immunohistochemistry of xenograft tissues

At the end of the study, one half of every tumor was fixed in formalin 10% (pH = 6.9–7.1) O/N at 4°C before processing for paraffin embedding and the other half was embedded in Optimun Cutting Temperature compound (VWR chemicals). At least 2 serial sections of 5 μm were stained with: hematoxylin and eosin (H&E) for general histology. For Ki67 staining, paraffin embedded tissues sections were deparaffinized, hydrated, and treated with boiling Citrate buffer pH 6.0 under microwaves for 20 min. Sections were blocked in 2% BSA. Primary monoclonal antibody Ki67 (SP6, Thermofisher) and HRP-conjugated secondary antibodies (Southern Biotech) were incubated in diluent for 1–24 h. Visualization was performed with DAB (Vector) and counterstained with hematoxylin; Photomicrographs were acquired using a Nikon ECLIPSE E200 microscope equipped with a Nikon Digital Sight DS-5M digital camera. For immunofluorescent staining, 4μm frozen sections were fixed in 4% paraformaldehyde or cold acetone for 10 minutes, permeabilized with 0.2% Triton X-100 for 30 min and blocked with 5% BSA for 1 hour. Primary antibodies (anti-fibroblasts clone TE-7, Millipore and phospho AKT/4060 Cell Signaling) were diluted in 3%BSA and incubated overnight at 4°C. Then, samples were washed with PBS-0.05% Tween-20 and incubated with fluorescent-conjugated secondary antibody (Alexa 488-conjugated mouse and rabbit antibody; Invitrogen) for 1hour. All samples were counterstained with DAPI. Photomicrographs were acquired using Leica Fluorescent Microscope (Leica DM 2000).

### MMTV-PyMT transgenic mouse model

Four-week-old MMTV-PyMT transgenic mice under pathogen-free conditions were administered with either 10 mg/kg body weight of the selective PIK3Cδ inhibitor CAL-101 (#S2226, Selleckchem) or the control vehicle by daily oral gavage for 6 weeks. At the age of 10 weeks, mice (n=4 / group) were harvested and the mammary tumors, lungs and livers were fixed in 4% paraformaldehyde and embedded in paraffin.

### TCGA BC dataset deconvolution pipeline

We estimated the specific expression of different PIK3C isoforms in fibroblasts in different BC subtypes (including ERα+, HER2+ and TNBC) RNA-seq samples from TCGA (https://portal.gdc.cancer.gov/projects/TCGA-BRCA) by applying a recently developed deconvolution algorithm CDSeq (https://github.com/kkang7/CDSeq) to the bulk RNA-seq samples of 1051 BC patients. First, we downloaded the RNA-seq HTSeq read count data for those samples from the TCGA data portal (https://portal.gdc.cancer.gov); we performed no additional normalization. Next, we extracted the read counts value for 45 known breast-cancer-associated fibroblast (CAF) marker genes (compiled from Le Bleu et. al. (75)) and 547 immune marker genes for various immune subtypes based on a reference profile developed by CIBERSORT (76) (**Supplementary Table 13**). Next, we used CDSeq to estimate the proportions of CAFs in each of the 1051 samples and the corresponding cell-type-specific gene expression profiles (GEPs). We set the number of cell types to 20 (which should be large enough to detect the major cell types in the data). CDSeq identified four CAFs in which the CAF marker genes had 1.5-fold or higher expression compared to expression of immune marker genes. Then, for each BC sample, we estimated CAF-expressed PIK3C-isoforms’ expression levels by multiplying the PIK3C-isoform expression level in four CDSeq-estimated CAFs expressions with their estimated proportions. Finally, we analyzed the association between the different BC subtype patients’ survival time and their estimated CAFs-expressed PIK3C-isoforms’levels. Samples were grouped into “High CAF-PIK3C isoform’ if their CAFs-expressed PIK3C isoform levels were above the 60th percentile of CAFs-expressed PIK3C isoform levels of all samples, and into “Low CAF-PIK3C isoform” if their CAFs-expressed PIK3C isoform levels were below the 40th percentile. For the survival analysis, we employed a MATLAB function MatSurv (https://github.com/aebergl/MatSurv) which generates a Kaplan-Meier plot and calculates the estimated Hazard Ratio (HR) and associated p-value (log-rank test).

### Immunohistochemistry and immunofluorescence staining of transgenic tissues

Paraffin-embedded samples were sectioned at 4 μm thickness. Antigen retrieval was performed by incubation of the slides in a pressure cooker for 5 min in 0.01 M citrate buffer, pH 6.0, and subsequent treatment with 3% hydrogen peroxide for 5 min. Slides were incubated overnight at 4 °C with antibodies to the following: α-SMA (Abcam, ab21027, 1:00), F4/80 (Biolegend, 123101, 1:20), PIK3Cδ (Santa Cruz, sc-55589, 1:30), pAkt-S473 (Cell Signaling Technology, CST, 4060, 1:100), pAkt-Thr308 (Millipore, 07-1398, 1:100), E-cadherin (CST, 14472S, 1:100), Vimentin (CST, 5741S, 1:100), CD31 (Abcam, ab9498, 1:100), Ki67 (Abcam, ab15580, 1:50). Immunohistochemical staining was performed according to the manufacturer’s instructions. For immunofluorescence, specimens were incubated with Alexa Fluor secondary antibodies (Thermo Fisher Scientific, A-31571, A-31570, A-31572, and A-31573) for 1 hr at room temperature. DAPI was then used for counterstaining the nuclei and images were obtained by laser scanning confocal microscopy (LSM780, Zeiss).

### Clinical specimens

This study was conducted on n=179 TNBC cases with (n=86) or without (n=93) Lymphovascular Invasion (LVI) from patients presenting at Nottingham City Hospital (48, 49). Treatment and outcome data, including BC-overall survival (OS) and Disease-Free survival (DFS) was maintained on a prospective basis. OS was defined as the duration (in months) from the date of primary surgery to the patients’ survival at the end of follow up time. DFS was defined as the duration, in months, from primary surgical treatment to the occurrence of first local, regional, or distant metastasis.

The sample set of the ERα^+^ cohort (n=73) were newly diagnosed BC patients presenting at the National University Cancer Institute, Singapore who were treated with primary or neoadjuvant chemotherapy or endocrine therapy before surgery. Baseline tumor biopsies were taken before anti-cancer therapy and analyzed. All patients in this study provided written informed consent to be enrolled; the studies were approved by the local ethics review committee.

### Immunohistochemistry of clinical samples

PIK3Cδ protein expression was evaluated by immunohistochemistry (IHC) using the rabbit PIK3Cδ antibody (PA5-15250; ThermoFisher Scientific) in full-face FFPE breast cancer tissue sections using either the Novolink Max Polymer Detection system (Leica, Newcastle, UK) for the Nottingham TNBC cohort, or the Leica Bond Max automated system (Leica Biosystems, Nussloch GmbH) using the bond polymer refine detection kit (DS9800, Leica Biosystems). The antibody was incubated for 1 hour at a concentration of 1:100. PIK3Cδ cytoplasmic immunoreactivity in the invasive tumors cells was assessed using the modified Histo-score (H score) method (77). Surrounding fibroblasts were identified by IHC expression of α-SMA (Abcam, ab5694) in serial FFPE sections. The antibody was incubated for 1 hour incubated using a 1:500 concentration. As the intensity of PIK3Cδ was not variable in inter-tumoral fibroblasts, it was assessed as a percentage of positively-stained cells.

### Statistical analysis of clinical samples

IBM SPSS 22.0 (Chicago, IL, USA) software was used for statistical analysis. The H-scores of tumoral PIK3Cδ expression did not follow a normal distribution and expression of PIK3Cδ protein was dichotomized using cut points derived from prediction of patient survival using X-tile (http://tissuearray.org; Yale University), with OS as the endpoint. Kaplan–Meier and log-rank analysis were used to assess OS and DMFS. Multivariate Cox Regression analysis with adjustment of covariates was fitted to test independence from standard prognostic factors in breast cancer (tumor grade, tumor size, lymph nodal stage, age, and LVI status) was used to identify independent prognostic value. Spearman’s correlation coefficient was carried out to examine the association between PIK3Cδ in the tumor and surrounding fibroblasts. P value ≤ 0.05 was considered significant.

### Ethics approval

This work obtained ethics approval by the North West – Greater Manchester Central Research Ethics Committee under the title; Nottingham Health Science Biobank (NHSB), reference number 15/NW/0685.

## Supplementary Figure Legends

**Supplementary Figure 1: CAF markers in primary and immortalized fibroblast cell lines and expression of PIK3Cδ in fibroblasts and breast cancer cell lines**. **(A)** Western blotting of CAF markers in HMF, MRC5 and in matching fibroblasts obtained from four TNBC patients (CAFs<5cm; NFs >5cm distance from the tumor site). GADPH was used as loading control. **(B)** PIK3Cδ expression in fibroblasts (primary and immortalized cell lines). GAPDH was used as loading control. **(C)**. Comparison of PIK3Cδ expression in breast cancer cells and primary fibroblasts. Tubulin was used as loading control.

**Supplementary Figure 2: 3D spheroid formation and invasion of fluorescently labelled cells**. Representative image showing the distribution of fluorescently labelled fibroblast and TNBC cells. During spheroids formation, fibroblast (green; SP-DiOC18) and TNBC cells (red; Dil Stain) form a compact agglomerate, making it difficult to distinguish between the two cell types. However, during the invasion step (addition of Matrigel), cells distribute differently, with fibroblasts localized at the core of the spheroids, while TNBC cells leading the invasion into the Matrigel.

**Supplementary Figure 3: Distribution of siRNA effects on MDA-MB-231 invasion**. Graphs depicting the Δ_Ratio_ siRNA screening values and distribution plots for different kinases in (A) HMF and (B) MRC5 cells. P-values were calculated using Student’s t-test and were further processed using the Bonferroni correction method. Lower panels show a magnification of the graph for Kinases with Δ_Ratio_<0.5. The overlapping hits (PIK3Cδ and AURKA) are highlighted in blue in both graphs.

**Supplementary Figure 4: mRNA expression levels of PIK3Cδ in different cell lines/types and examination of TNBC expression of PIK3Cδ following co-culturing with fibroblasts**. **(A)** qRT-PCR of PIK3Cδ expression levels in BC, lymphoid and primary or immortalized fibroblast cell lines. **(B)** RNAseq data, courtesy of Human Protein Atlas (www.proteinatlas.org) showing the expression of PIK3Cδ in various lymphoid, myeloid, fibroblast and BC cell lines.**(C)** Western blotting of PIK3Cδ expression in MDA-MB-231 and BT-549 cells, following co-culturing with fibroblast cells (MRC5 or HMF) for a period of 9 days. GADPH was used as loading control. **(D)** Upper panel: Western blotting of PIK3Cδ expression in primary CAFs isolated from human TNBC samples. Lower panel: Effects of co-culturing CM isolated from primary CAFs on the expression of PIK3Cδ in BT-549 cells (for a period of 6 days). GADPH was used as loading control.

**Supplementary Figure 5: Effects of silencing or overexpressing of fibroblast PIK3Cδ in TNBC invasion**. The effects of PIK3Cδ silencing in HMF (pool of 3 different siRNAs) on **(A)** MDA-MB-231 or **(B)** BT-549 invasion were assessed following the 3D experimental co-culture procedure described in Figure 1 (n=3 independent experiments, with at least 4 technical replicates). Significance was calculated using unpaired t-test. Results are expressed as mean ± SEM; * p<0.05, *** P < 0.001 vs. control-siRNA transfected fibroblasts. **(C)** The knockdown efficacy of the PIK3Cδ siRNAs in HMF was verified by qRT-PCR. The effects of PIK3Cδ overexpression in HMF, using pCMV6-AC-PIK3Cδ-GFP plasmid, on **(D)** MDA-MB-231 or **(E)** BT-549 invasion were assessed following the same experimental procedure described in Figure 1 (n=3 independent experiments, with at least 4 technical replicates). Significance was calculated using unpaired t-test. Results are expressed as mean ± SEM; * P < 0.05 vs. control-siRNA transfected fibroblasts. **(F)** The overexpression of PIK3Cδ in HMF was verified by qRT-PCR. **(G)** HMF and MRC5 cells were either transfected with pCMV6-AC-PIK3Cδ-GFP plasmid or silenced with PIK3Cδ siRNA for 24h and the CM from these treatments was collected. MDA-MB-231 were seeded in a 96 well plate (3000 cell/well), treated with the aforementioned CM for 48h and cell viability was assessed by CellTiter-Glo assay using a Glow-max reader (n=3 independent experiments, with 2 technical replicates). Significance was calculated using one-way ANOVA. Results are expressed as mean ± SEM.

**Supplementary Figure 6: HT-RNAi screening validation**. **(A)** Genomic inhibition validation. Different shRNA plasmids targeting randomly selected kinases were used to transfect MRC5. 48h after transfection, fibroblasts were co-cultured with MDA-MB-231 cells and a 3D invasion assay was performed. Upper panel: Heatmap displaying the results of the shRNA inhibition compared to the original screening (siRNA) with regards to the invasion potential of MDA-MB-231. Lower panel: Representative pictures of the 3D invasion assay for certain kinases are shown. **(B)** qRT-PCR results demonstrating the silencing efficiency of the selected shRNAs plasmids against the respective kinases. **(C)** Pharmacological inhibition validation. MRC5 were pre-treated with 10 µM of the indicated compounds for 24h and then co-cultured in 3D with MDA-MB-231. Upper panel: Heatmap displaying the results of the chemical inhibition compared to the original screening (siRNA) with regards to the invasion potential of MDA-MB-231. Lower panel: Representative pictures of the 3D invasion assay for certain kinases are shown. **(D)** HMF and MRC5 cells were transfected with either siRNA or shRNA plasmids targeting PIK3Cδ or AURKA respectively and the cell viability was monitored 48h later using CellTiter-Glo assay. (n=3 independent experiments, with at least 2 technical replicates). Significance was calculated using two-way ANOVA. Results are expressed as mean ± SEM; ** P < 0.01, **** P < 0.0001 vs. control transfected fibroblasts. **(E)** HMF and MRC5 cells were transfected with pCMV6-AC-PIK3Cδ-GFP plasmid and the cell viability was monitored 48h later using CellTiter-Glo assay. (n=3 independent experiments, with at least 2 technical replicates). Significance was calculated using two-way ANOVA. Results are expressed as mean ± SEM.

**Supplementary Figure 7: Effect of CAL-101 inhibitor on fibroblast cell lines**. **(A)** Western blotting of downstream targets of PIK3Cδ following treatment of HMF and MRC5 cells with increased concentrations of CAL-101. Each experiment was performed in triplicate. **(B)** HMF and MRC5 cells were treated with 5μΜ and 10μΜ of CAL-101 and the cell viability was monitored 48h later using CellTiter-Glo assay (n=3 independent experiments, with at least 4 technical replicates). Significance was calculated using two-way ANOVA test. Results are expressed as mean ± SEM; **** P < 0.0001 vs. vehicle (DMSO) treated cells.

**Supplementary Figure 8: Effects of chemical inhibition of PIK3Cδ on BT-549 cell invasion**. **(A)** 3D invasion assay: HMF (left panel) and MRC5 (right panel) were pre-treated with either vehicle (DMSO) or with 1, 5 and 10 µM of CAL-101 for 24h. Following, fibroblasts were co-cultured with BT-549 as 3D spheroids and the invasion potential was measured. Representative pictures of the 3D invasion assay at different time points are shown (n=3 independent experiments, with at least 4 technical replicates). Significance was calculated using unpaired t-test. Results are expressed as mean ± SEM; * P < 0.05 vs. DMSO treated fibroblasts. **(B)** 2D invasion assay: HMF (left panel) and MRC5 (right panel) were pre-treated with either vehicle (DMSO) or with 1, 5 and 10 µM of CAL-101 for 24h and were then seeded on the lower chamber of a transwell insert. BT-549 cells were seeded on the Matrigel-coated upper chamber of the transwell insert (pore size: 8 μm allowing cell-crossing) and were co-cultured with the fibroblasts. After 24h of co-culture, migrated BT-549 cells were fixed and stained with crystal violet and counted using an inverted microscope (n=3 independent experiments, with at least 6 technical replicates). Significance was calculated using one-way ANOVA test. Data are expressed as mean ± SEM; ** P < 0.01, **** P < 0.0001 vs. DMSO treated fibroblasts. **(C)** 2D real-time invasion assay: HMF (left panel) and MRC5 (right panel) were pre-treated with either vehicle (DMSO) or with 10 µM of CAL-101 for 24h and were then seeded on the lower chamber of a transwell insert. BT-549 cells were seeded on the upper chamber of the transwell insert (pore size: 0.4 μm not allowing cell-crossing) and were co-cultured with the fibroblasts. After 24h of co-culture, BT-549 were moved to CIM-Plates and their relative invasion rate was monitored using the xCELLigence Real-Time Cell Analysis (RTCA) instrument (n=3 independent experiments). Significance was calculated using two-way Anova test. Results are expressed as mean ± SEM; *** P < 0.001, **** P < 0.0001 vs. DMSO treated fibroblasts. **(D)** 2D CM invasion assay: HMF (left panel) and MRC5 (right panel) were treated with vehicle only or with 10 µM CAL101 in serum free for 24h in order to obtain the CM. BT-549 were then incubated with HMF or MRC5 CM for 2D invasion assays (n=3 independent experiments). Significance was calculated using unpaired t-test. Data are expressed as mean ± SEM; * P < 0.05, ** P < 0.01 vs. DMSO treated fibroblasts’ CM. **(E)** 2D invasion assay: TNBC patients’ CAFs were pre-treated with either vehicle (DMSO) or with 10 µM of CAL-101 for 24h and were then seeded on the lower chamber of a transwell insert. BT-549 cells were seeded on the Matrigel-coated upper chamber of the transwell insert (pore size: 8 μm allowing cell-crossing) and were co-cultured with the TNBC CAFs. After 24h of co-culture, migrated BT-549 cells were fixed and stained with crystal violet and counted using an inverted microscope. (*n*=3 independent experiments, with at least 4 technical replicates). Significance was calculated using unpaired t-test. Data are expressed as mean ± SEM; **** P < 0.0001 vs. DMSO treated fibroblasts.

**Supplementary Figure 9: Recovery of PIK3Cδ mRNA levels after silencing with shRNA or siRNA (*data corresponding to* Figure 2F).** MRC5 were transfected with control or PIK3Cδ siRNA (left panel) or control or PIK3Cδ shRNA (right panel). Silenced MRC5 were transfected 24h later with the pCMV6-AC-PIK3Cδ-GFP plasmid. PIK3Cδ expression levels were monitored during the course of the experiment. Significance was calculated using unpaired t-tests. Results are expressed as mean ± SEM; ****P < 0.01 vs. control or vs. PIK3Cδ silenced cells.

**Supplementary Figure 10: Effects of various PI3K inhibitors on TNBC cell invasion and fibroblasts’ cell viability**. **(A)** Table showing the selectivity of specific inhibitors against different PI3K isoforms. **(B)** 2D invasion assay: MRC5 and HMF cells were pre-treated with either vehicle (DMSO) or with different concentrations of various PIK3 inhibitors for 24h and were then seeded on the lower chamber of a transwell insert. MDA-MB-231 and BT-549 cells were seeded on the Matrigel-coated upper chamber of the transwell insert (pore size: 8 μm allowing cell-crossing) and were co-cultured with the fibroblasts. After 24h of co-culture, migrated MDA-MB-231 cells were fixed and stained with crystal violet and counted using an inverted microscope (n=2 independent experiments, with at least 5 technical replicates). Significance was calculated using one-way ANOVA test. Data are expressed as mean ± SEM; * P < 0.05, *** P < 0.001, **** P < 0.0001 vs. DMSO treated fibroblasts. **(C)** MRC5 cells and HMF were treated with different concentrations of various PIK3 inhibitors and the cell viability was monitored 48h later using CellTiter-Glo assay. (n=2 independent experiments, with at least 2 technical replicates) Significance was calculated using two-way ANOVA test. Results are expressed as mean ± SEM.

**Supplementary Figure 11: Principal component analysis (PCA) plot and cluster analysis of RNA-seq samples**. **(A)** PCA performed using DESeq2 rLog-normalized RNAseq data. Loadings for principal components 1 (PC1) and PC2 are reported in the graph. **(B)** Hierarchical clustering analyses performed using DESeq2 rLog-normalized RNAseq data. Color code from white to dark blue refers to the distance metric used for clustering, with dark blue referring to maximum of correlation values.

**Supplementary Figure 12: Fibroblast-PIK3Cδ / cancer epithelial cells-NR4A1 paracrine signaling axis promotes TNBC invasion**. **(A)** Western blotting of NR4A1 expression in a panel of BC cell lines. GADPH was used as loading control. **(B)** 2D invasion assay: BT-549 cells were seeded on the Matrigel-coated upper chamber of the transwell insert (pore size: 8 μm allowing cell-crossing) and were treated with either vehicle (DMSO) or Cytosporone B (5 µM). After 24h, migrated BT-549 cells were fixed and stained with crystal violet and counted using an inverted microscope (n=3 independent experiments, with at least 4 technical replicates). Significance was calculated using unpaired t-test. Results are expressed as mean ± SEM; **** P < 0.0001 vs. DMSO treated cells. **(C)** HMF cells were treated with CAL-101 or DMSO. NR4A1-silenced (siRNA) MDA-MB-231 or control-siRNA cells were seeded on the Matrigel-coated upper chamber of a transwell insert (pore size: 8 μm allowing cell-crossing) and were co-cultured with fibroblasts. After 24h of co-culture, migrated MDA-MB-231 cells were fixed and stained with crystal violet and counted using an inverted microscope. (n=3 independent experiments, with at least 3 technical replicates). Significance was calculated using two-way ANOVA test. Results are expressed as mean ± SEM; ** P < 0.01, ***P<0.001, **** P < 0.0001 vs. samples indicated in the graph.

**Supplementary Figure 13: Schematic of integrated secretome and RNAseq analysis**.

**Supplementary Figure 14: Effects of BDNF or PLGF treatment on NR4A1 expression and invasion of BT-549 cells**. **(A)** qRT-PCR of *NR4A1* expression levels in BT-549 cells following treatment with PLGF and BDNF (or control PBS/BSA (0.1%)). **(B)** 2D invasion assay: BT-549 cells were seeded on the Matrigel-coated upper chamber of the transwell insert (pore size: 8 μm allowing cell-crossing) and were treated with PLGF or BDNF (or control PBS/BSA (0.1%)). After 24h, migrated BT-549 cells were fixed and stained with crystal violet and counted using an inverted microscope (n=3 independent experiments, with at least 4 technical replicates). Significance was calculated using unpaired t-test. Results are expressed as mean ± SEM; ** P < 0.01, *** P < 0.001, **** P < 0.0001 vs. vehicle treated cells.

**Supplementary Figure 15: Effects of BDNF or PLGF silencing on CAL-101 mediated TNBC cell invasion**. MRC5 cells transfected with siRNA for BDNF and/or PLGF were pre-treated with either vehicle (DMSO) or 10 µM of CAL-101 for 24h. MDA-MB-231 cells were seeded on the Matrigel-coated upper chamber of a transwell insert (pore size: 8 μm allowing cell-crossing) and were co-cultured with the fibroblasts. After 24h of co-culture, migrated MDA-MB-231 cells were fixed, stained with crystal violet and counted using an inverted microscope (n=3 independent experiments, with at least 3 technical replicates). Significance was calculated using two-way ANOVA test. Results are expressed as mean ± SEM; * P < 0.01 vs. DMSO treated control cells.

**Supplementary Figure 16: Effects of CAL-101 treatment on MDA-MB-231 cell viability**. MDA-MB-231 cells were treated with either vehicle (DMSO) or increasing concentrations of CAL-101 and cell viability was monitored after 24h, 48h and 72h using CellTiter-Glo assay. (n=3 independent experiments, with at least 5 technical replicates). Significance was calculated using two-way ANOVA test. Results are expressed as mean ± SEM; * P < 0.05, ** P < 0.01 vs. vehicle treated cells.

**Supplementary Figure 17: Expression of CD68+ cells in the immunocompromised orthotopic BC xenograft model**. Representative images of immunofluorescent staining of MDA-MB-231/MRC5 tumor cryosections for CD68 (red) and p-AKT (Thr308) (green) after vehicle or CAL-101 treatment. Original magnification, 40x. Scale bar, 50μm. The quantification of p-AKT (Thr308) immunofluorescent staining in tumor infiltrating CD68 macrophages were very low (white arrows) without difference between the groups. Statistics were not performed.

**Supplementary Figure 18: Effects of CAL-101 treatment on tumor growth of MMTV-PyMT transgenic mice**. **(A)** Tumor volumes chart from MMTV-PyMT transgenic mice after vehicle or CAL-101 treatment Individual values for each mouse are displayed. Significance was calculated using unpaired t-test (week 15). Results are expressed as mean ± SEM; * P < 0.05. **(B)** Images of tumors, lungs and livers of MMTV-PyMT transgenic mice after vehicle or CAL-101 treatment are shown. Yellow arrows indicate gross appearance of metastatic lung tumor nodes in different mice from both treatment groups. **(C)** Representative images of lung tissues and hematoxylin and eosin staining from MMTV-PyMT mice that were treated with either vehicle or CAL-101. (Scale bar, 5mm). **(D)** Upper panel: Representative images of immunofluorescent staining for α-SMA and total PIK3Cδ in the mammary tumor sections of MMTV-PyMT transgenic mice after vehicle or CAL-101 treatment. Arrows indicate a-SMA^+^ fibroblasts. Higher-magnification images are shown at the bottom right corner. Lower panel: Quantification of p-total PIK3Cδ immunofluorescent staining in tumor infiltrating a-SMA^+^ fibroblasts in the mammary tumors of MMTV-PyMT transgenic mice after vehicle or CAL-101 treatment. Significance was calculated using multiple t-tests. Results are expressed as mean ± SEM. **(E)** Upper panel: Representative images of immunofluorescent staining for F4/80 and total PIK3Cδ in the mammary tumor sections of MMTV-PyMT transgenic mice after vehicle or CAL-101 treatment. Arrows indicate F4/80^+^ macrophages. Higher-magnification images are shown at the bottom right corner. Lower panel: Quantification of total PIK3Cδ immunofluorescent staining in tumor infiltrating F4/80^+^ macrophages in the mammary tumors of MMTV-PyMT transgenic mice after vehicle or CAL-101 treatment. Significance was calculated using multiple t-tests. Results are expressed as mean ± SEM. (F) Western blotting of PIK3Cδ expression in CAFs and cancer cells isolated MMTV-PyMT tumors. The expression of PIK3Cδ expression following co-culturing with CM isolated from CAFs for a period of 6 days was also examined. GADPH was used as loading control.

**Supplementary Figure 19: PIK3Cδ expression in tumor and fibroblast cells**. Representative images of low and high PIK3Cδ expression in tumor or surrounding fibroblast cells (a-SMA+) are shown. **(A)** Low PIK3Cδ expression in tumoural and intra-tumoural fibroblasts. **(B)** α-SMA positive expression in same field for quantifying PIK3Cδ expression in intra-tumoral fibroblasts. **(C)** High PIK3Cδ expression in tumoral and intra-tumoral fibroblasts. **(D)** α-SMA positive expression in same field for quantifying PIK3Cδ expression in intra-tumoral fibroblasts. Original magnifications: ×2, x5, 10x. Scale bars: 200, 500, 1000 μm. The red arrows represent the fibroblast expression of PIK3Cδ, while the black arrows represent the tumoral expression of PIK3Cδ.

**Supplementary Figure 20: Association of tumoral-PIK3Cδ expression with patients’ survival**. **(A)** Kaplan-Meier plots showing the association between tumoral-PIK3Cδ protein expression with OS (P = 0.004) in TNBC patients. **(B)** Kaplan-Meier plots showing the association between tumoral-PIK3Cδ protein expression with DFS (P = 0.009) in TNBC patients. **(C)** Kaplan-Meier plots showing the association between tumoral-PIK3Cδ mRNA expression with OS (P = 0.405) in TNBC patients following deconvolution of bulk TCGA RNA-seq samples. **(D)** Kaplan-Meier plots showing the association between CAF-PIK3Cα mRNA expression with OS (P = 0.014) in TNBC patients following deconvolution of bulk TCGA RNA-seq samples. **(E)** Kaplan-Meier plots showing the association between CAF-PIK3Cβ mRNA expression with OS (P = 0.0123) in TNBC patients following deconvolution of bulk TCGA RNA-seq samples. **(F)** Kaplan-Meier plots showing the association between CAF-PIK3Cγ mRNA expression with OS (P = 0.493) in TNBC patients following deconvolution of bulk TCGA RNA-seq samples. * P < 0.05, ** P < 0.01.

## Supplementary Table Legends

**Supplementary Table 1. TNBC patients’ clinical information from whom the primary fibroblasts were obtained from**. Primary fibroblasts were obtained from TNBC patients’ specimens. Patients’ characteristics including age, tumor type, tumor grade, ER/HER2 status, ER/HER2 score, response to treatment and survival status.

**Supplementary Table 2. siRNA Kinome screening results in HMF and MRC5**. **^A & J^** Kinase symbol; **^B,C & K^** Delta Ratio (replicate 1 or 2); **^D^** Average of Delta Ratio; **^E^** Standard deviation calculated from the two Delta Ratios in columns B and C; **^F & L^** P-Value. **^G & M^** P value corrected (Bonferroni correction). **^H & N^** Z score (average).

**Supplementary Table 3. Secretome report showing differentially expressed secreted A & M B,C & N,O proteins in CAL-101 treated HMF and MRC5 cells**. **^A&M^** RefSeq ID; **^B,C & N,O^** Ratio of Chemiluminescence signal measured in DMSO or CAL-101 treated cells (replicate 1 & 2). **^D,E & P,Q^** Normalized value were calculated according to manufacturer’s recommendation using value from positive control (replicate 1 & 2). **^F,G,H,I & R,S,T,U^** Log2 values for the normalized ratio DMSO/DMSO and DMSO/CAL-101. **^J & V^** Average of the Log2 normalized value of DMSO/CAL101 replicate. **^K & W^** P-value: Two-sample Student T-Test value to determine whether the DMSO/DMSO and DMSO/CAL-101 are likely to have come from the same two underlying populations that have the same mean.

**Supplementary Table 4. DESeq2 report showing differentially expressed genes for comparison model**. MDA-MB-231 cells co-cultured with CAL-101 treated vs DMSO treated MRC5 cells. **^A^** Ensembl ID; **^B^** Base Mean: Mean of the normalized read counts across all the samples; **^C^** Log2 fold change: Log2 fold change is the effect size estimate that measures how much gene’s expression have changed due treatment with doxorubicin treatment in comparison to treatment with DMSO; **^D^** LfcSE: Standard error estimate for the Log2 fold change estimate; **^E^** Stat: Wald-statistics to test whether the data provides sufficient evidence to conclude if the particular effect size estimate value is really different from zero; **^F^** P value: indicates the probability that a fold change as strong as the observed one, or even stronger, would be seen under DMSO treatment; **^G^** P adjusted: Benjamini-Hochberg (BH) corrected P values. BH method calculates an adjusted P value for each gene which answers the following question: if we called significant all genes with a P value less than or equal to this gene’s P value threshold, what would be the fraction of false positives among them; **^H^** Symbol: Gene symbol; **^I^** Entrez: entrez id of each gene.

**Supplementary Table 5. Sequences of primers used for the qRT-PCR validation**.

**Supplementary Table 6. Table showing functional annotation of pathways enriched for CAL-101 regulated secreted proteins. Table displaying the IPA predicted functional categories for pathways enriched with secreted proteins commonly upregulated by CAL-101 treatment of HMF and MRC5 cells.**

**Supplementary Table 7. Clinico-pathologic parameters of Nottingham series’ TNBC patients’ cohort.**

**Supplementary Table 8. Association of fibroblast-PIK3Cδ protein expression with TNBC patients’ outcome.**

**Supplementary Table 9. Association of tumoral-PIK3Cδ protein expression with TNBC patients’ outcome.**

**Supplementary Table 10. Clinico-pathologic parameters of ERα+ patients’ cohort from Singapore.**

**Supplementary Table 11. Association of fibroblast-PIK3Cδ protein expression with ERα+ patients’ outcome.**

**Supplementary Table 12. Association of tumoral-PIK3Cδ protein expression with ERα+ patients’ outcome.**

**Supplementary Table 13. List of BC CAF- and immune marker-genes.**

