## Supplementary Figures 1-20 for "Global kinome silencing combined with 3D invasion screening of the tumor microenvironment identifies fibroblast-expressed PIK3Cδ involvement in triple-negative breast cancer progression"

### Supplementary Figure 1

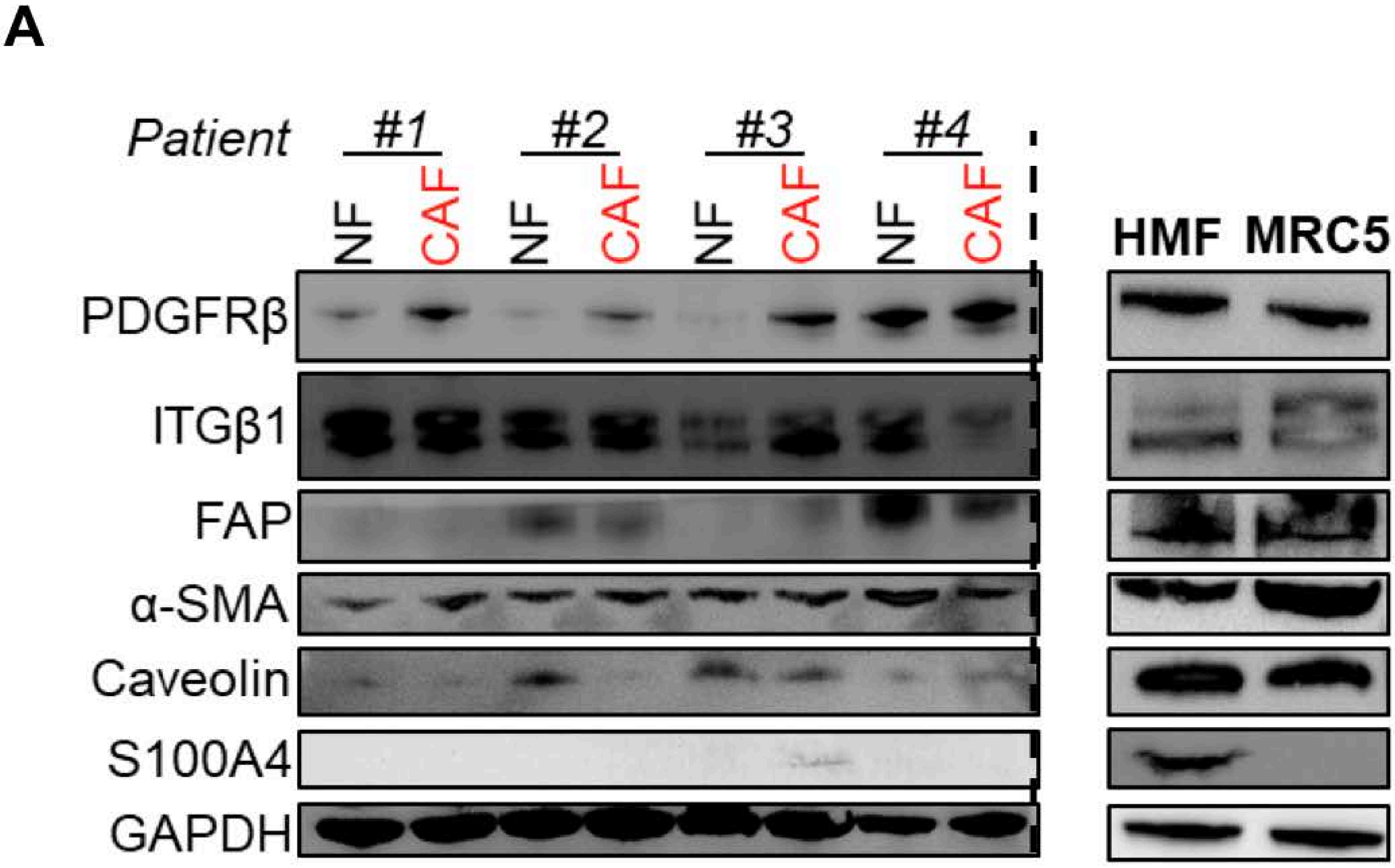


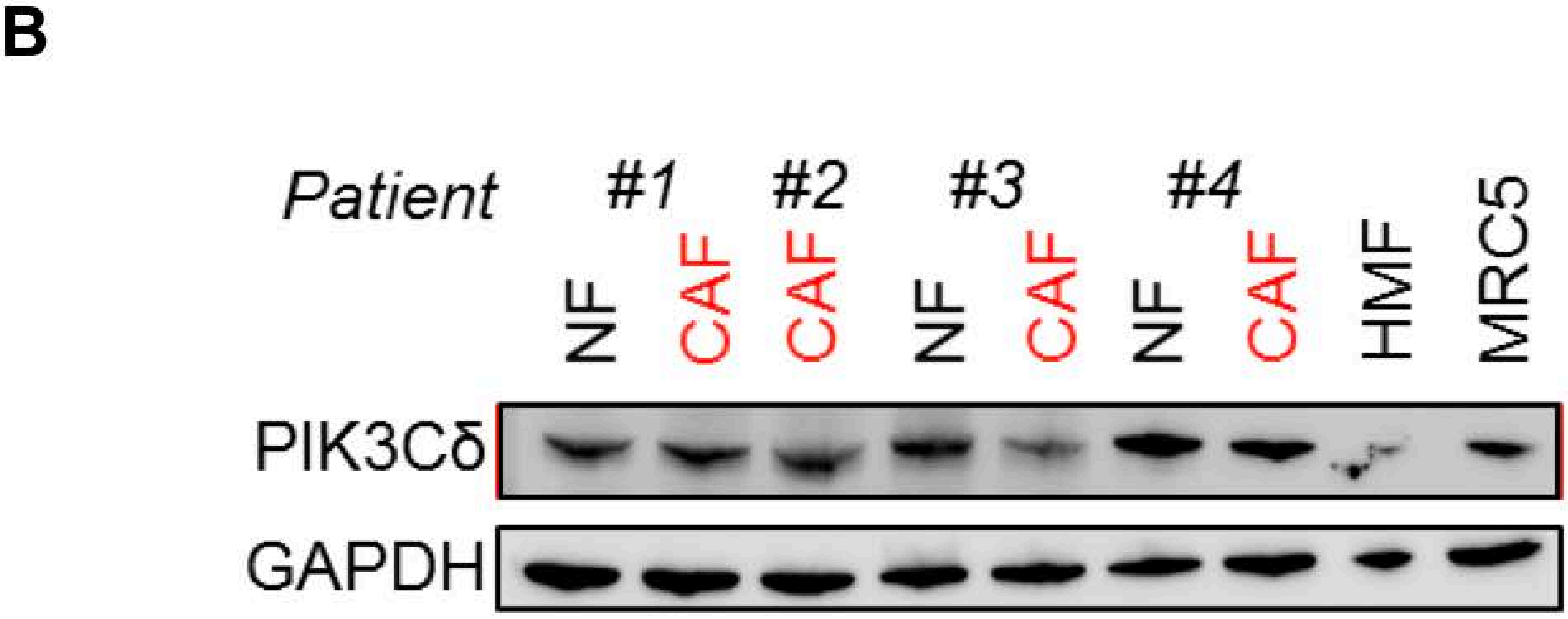


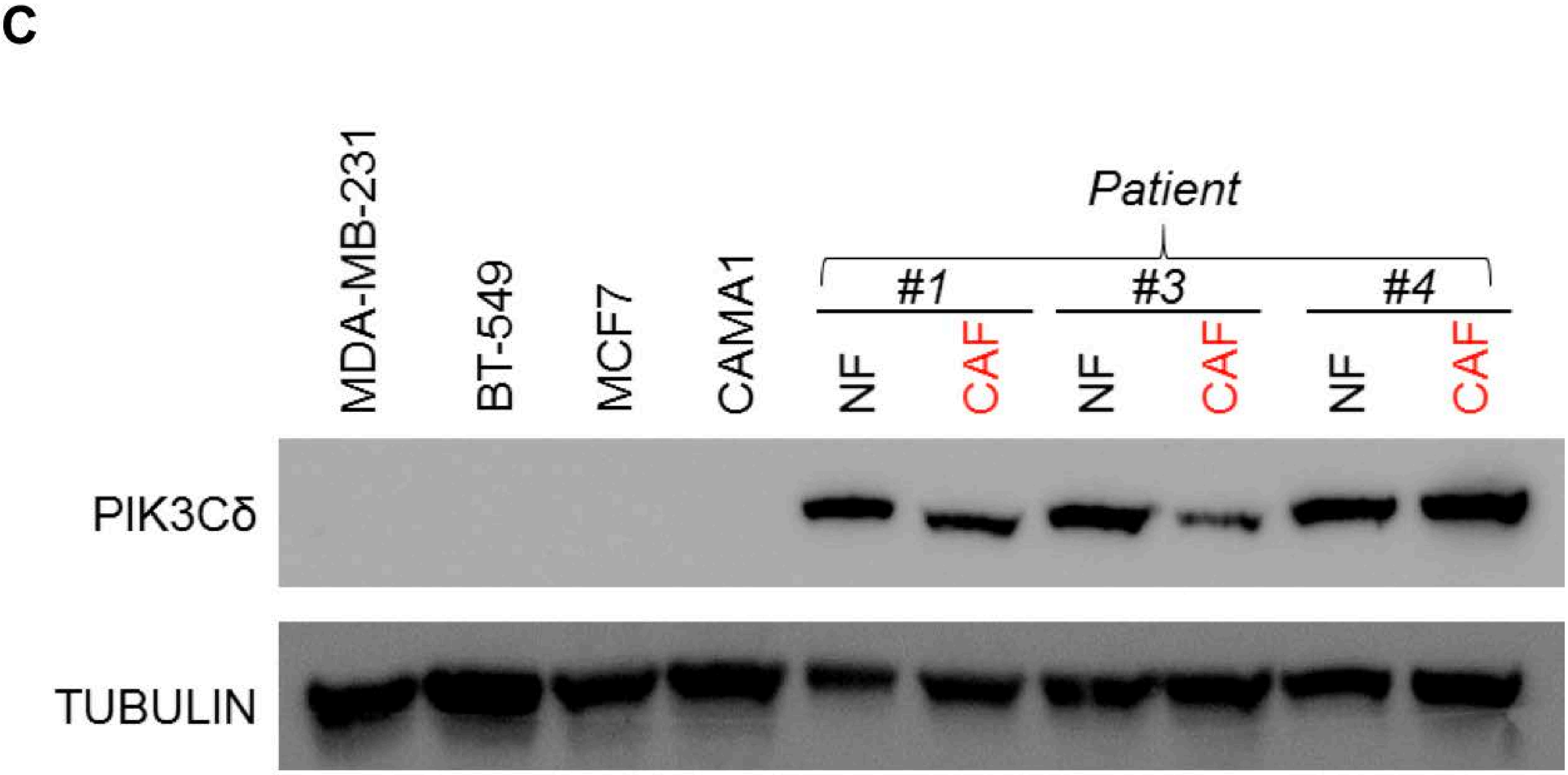

### Supplementary Figure 2

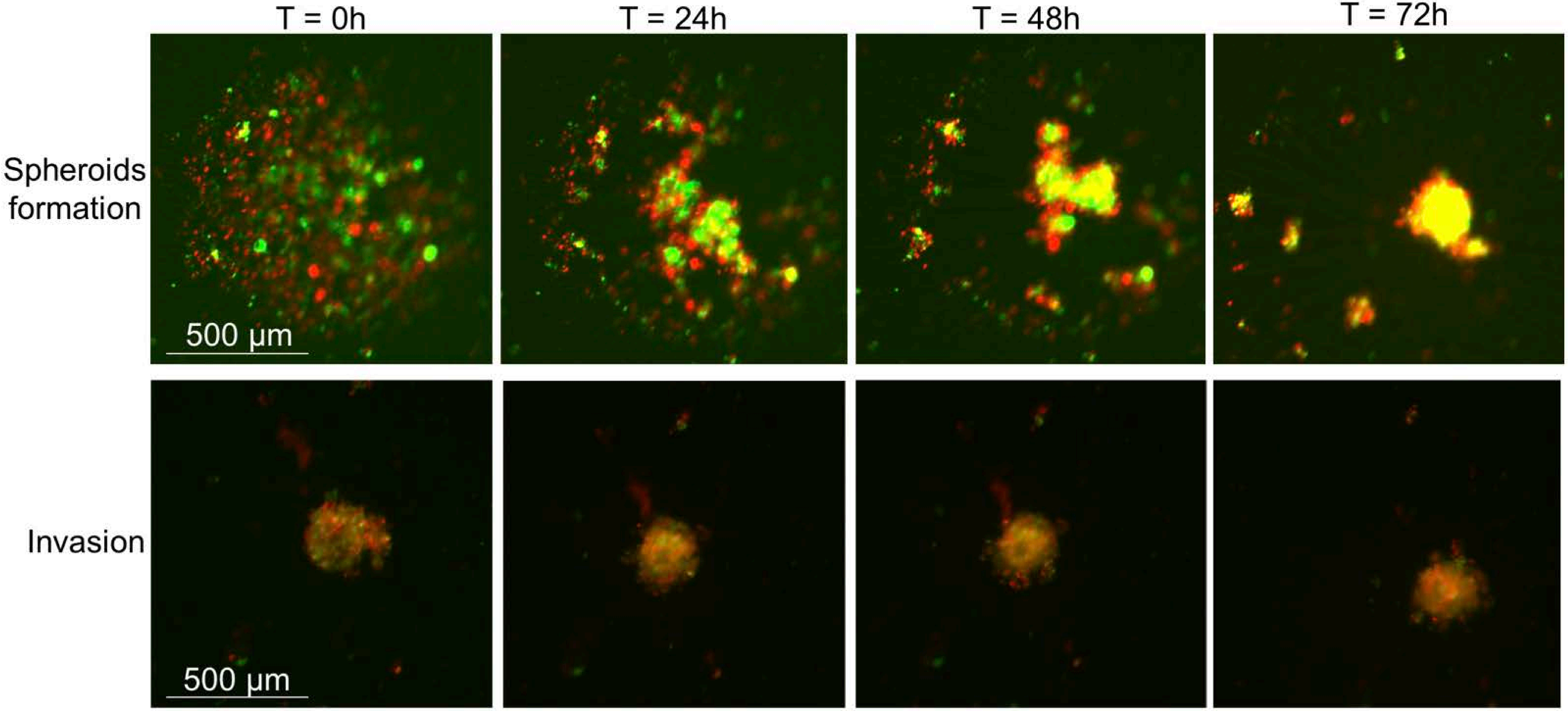

### Supplementary Figure 3

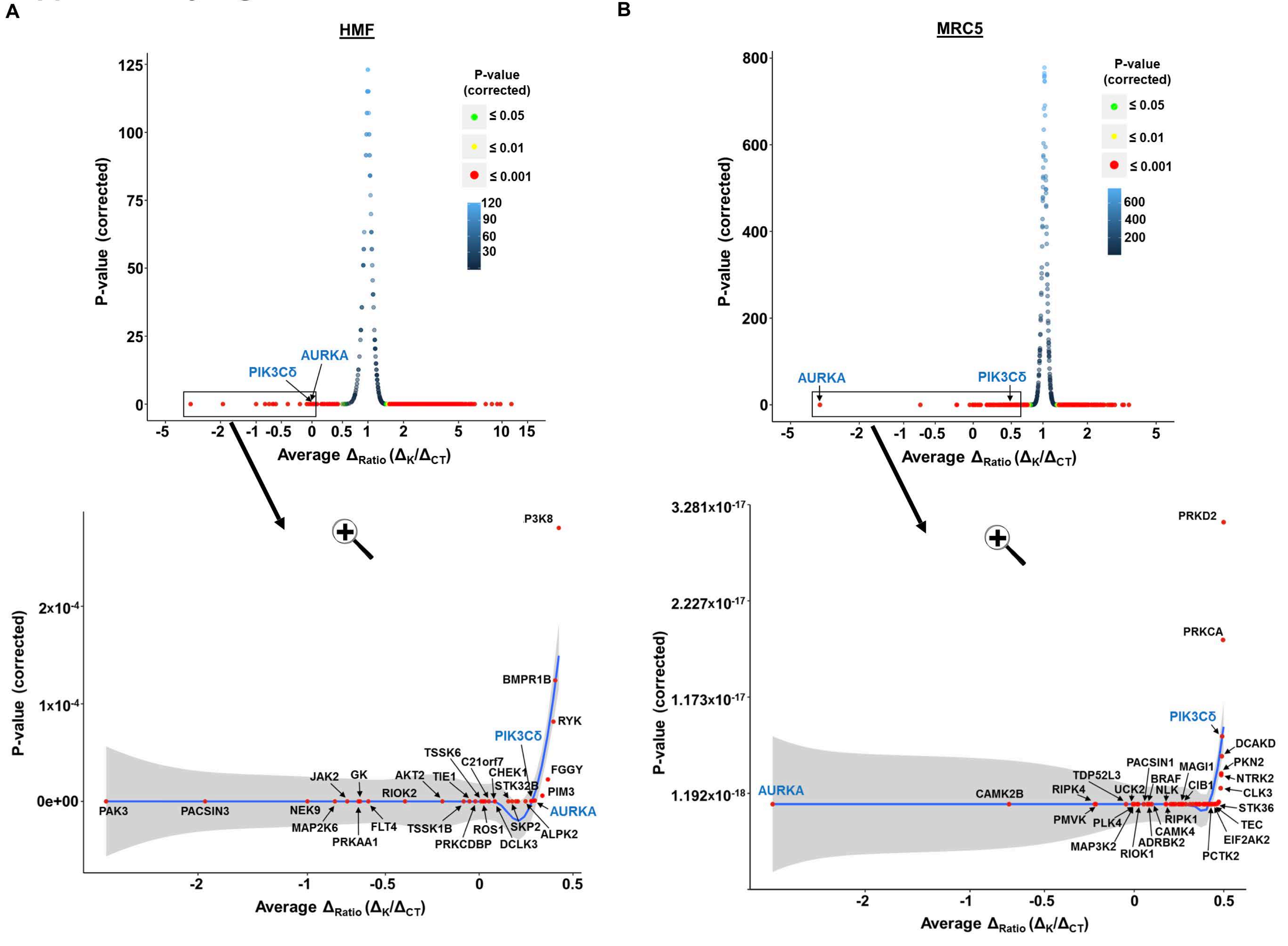

### Supplementary Figure 4

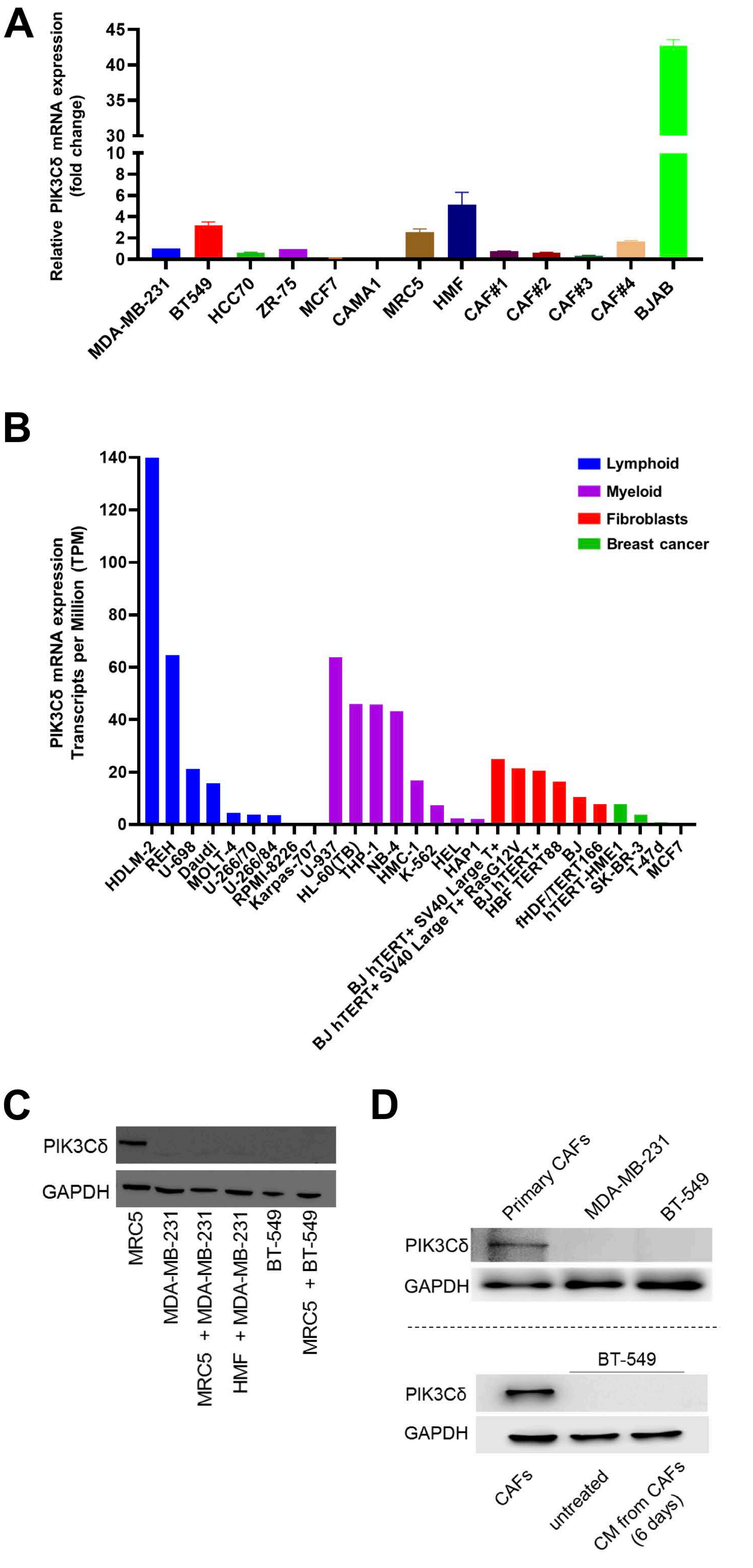

### Supplementary Figure 5

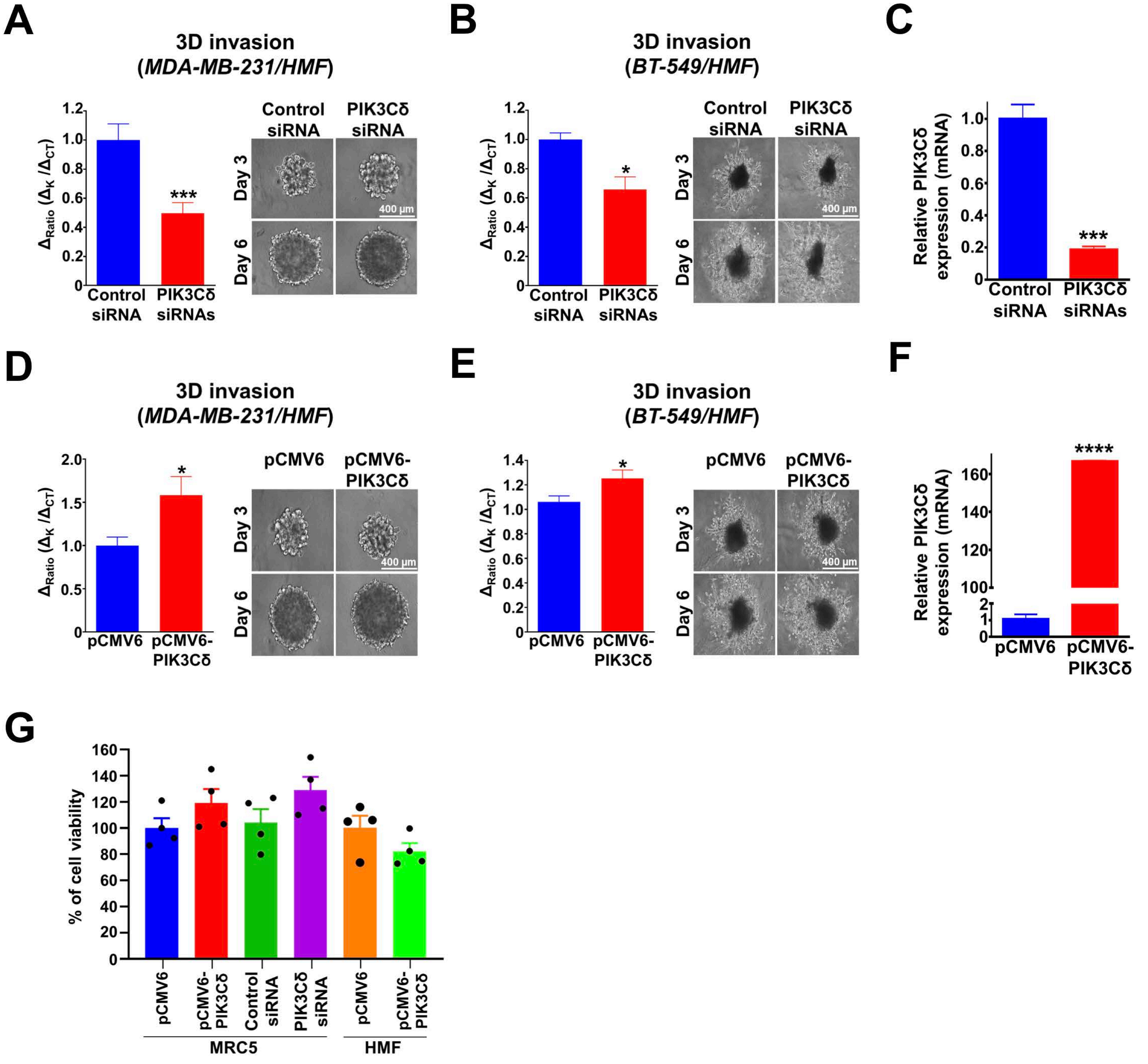

### Supplementary Figure 6

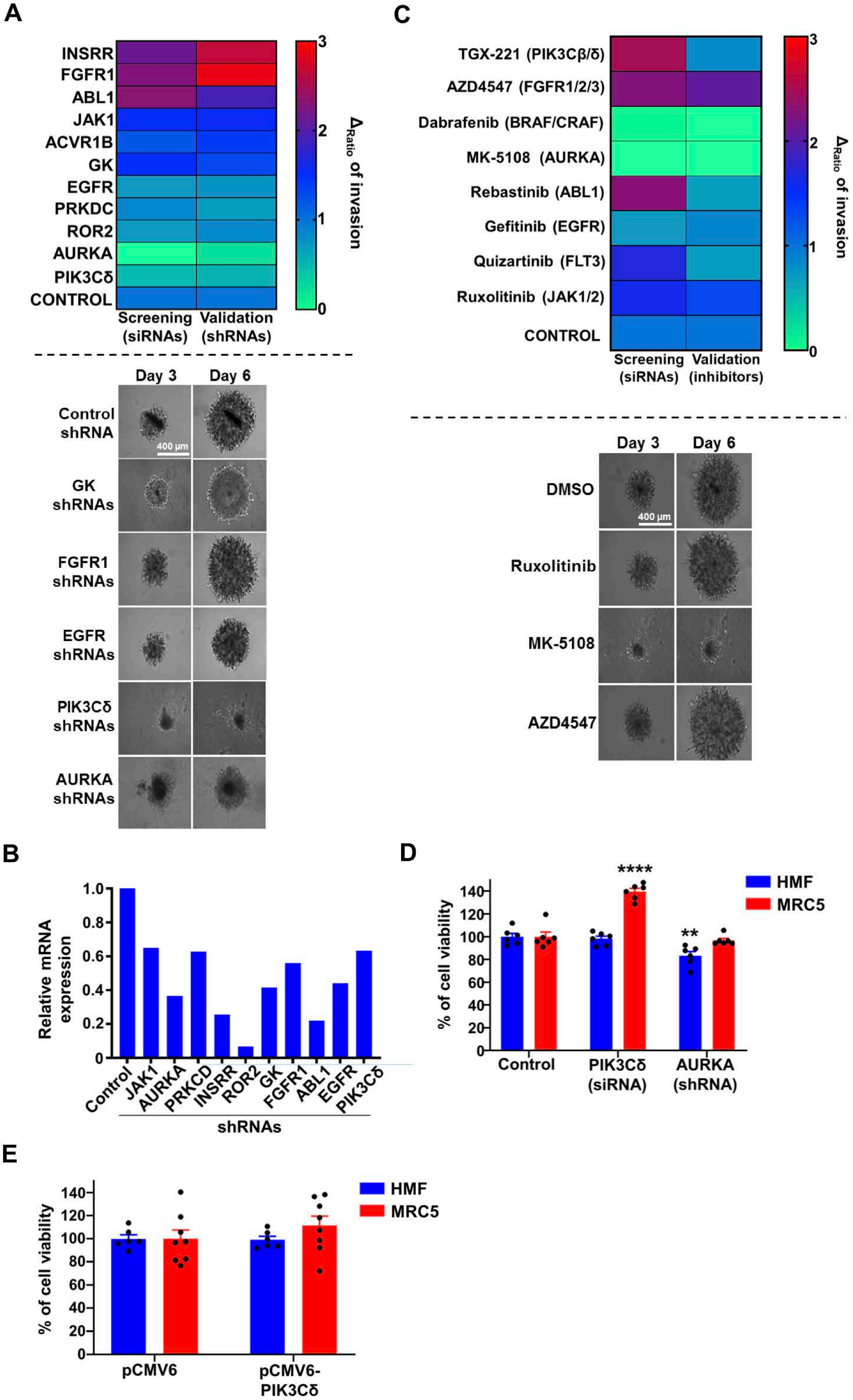

### Supplementary Figure 7

A

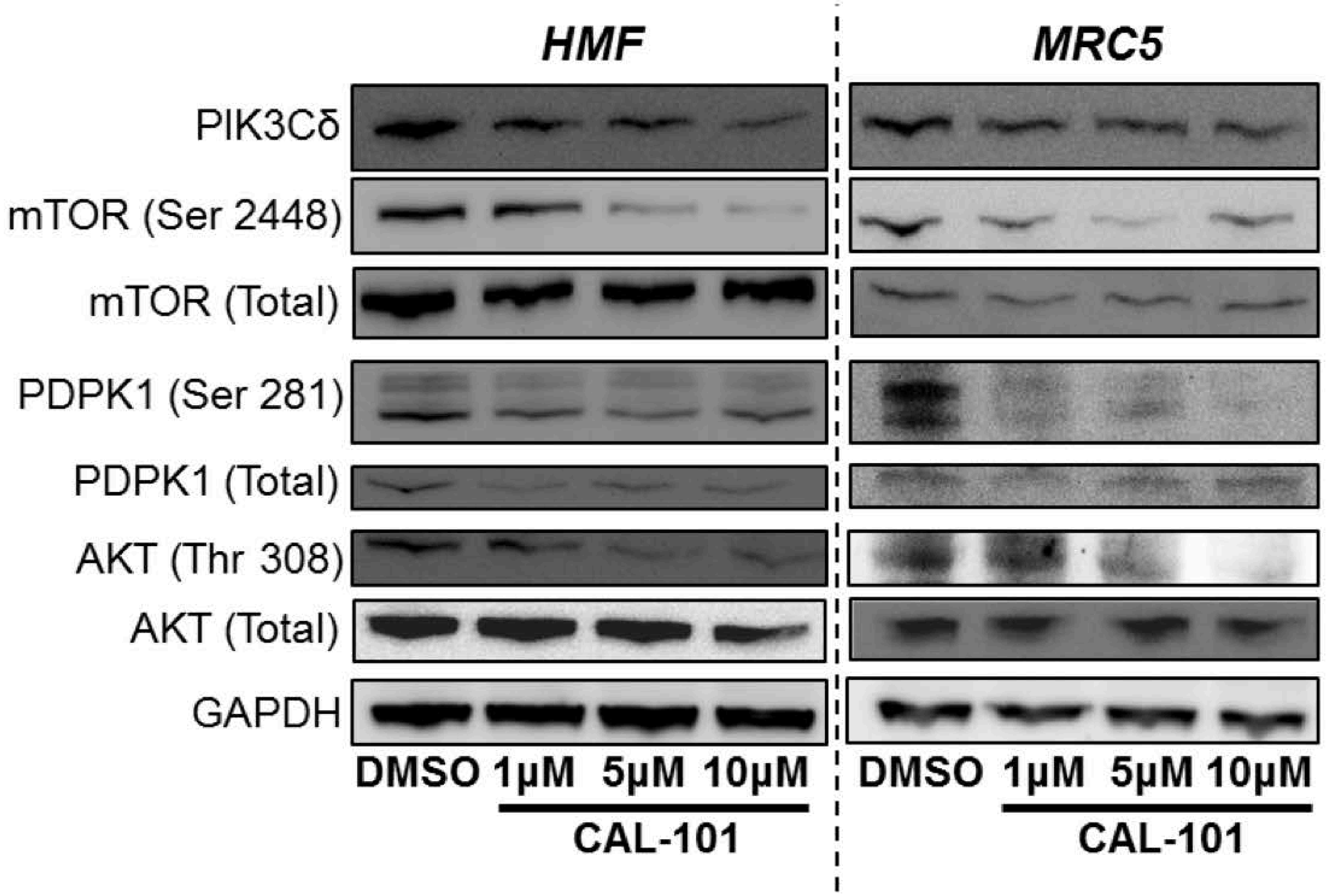


B

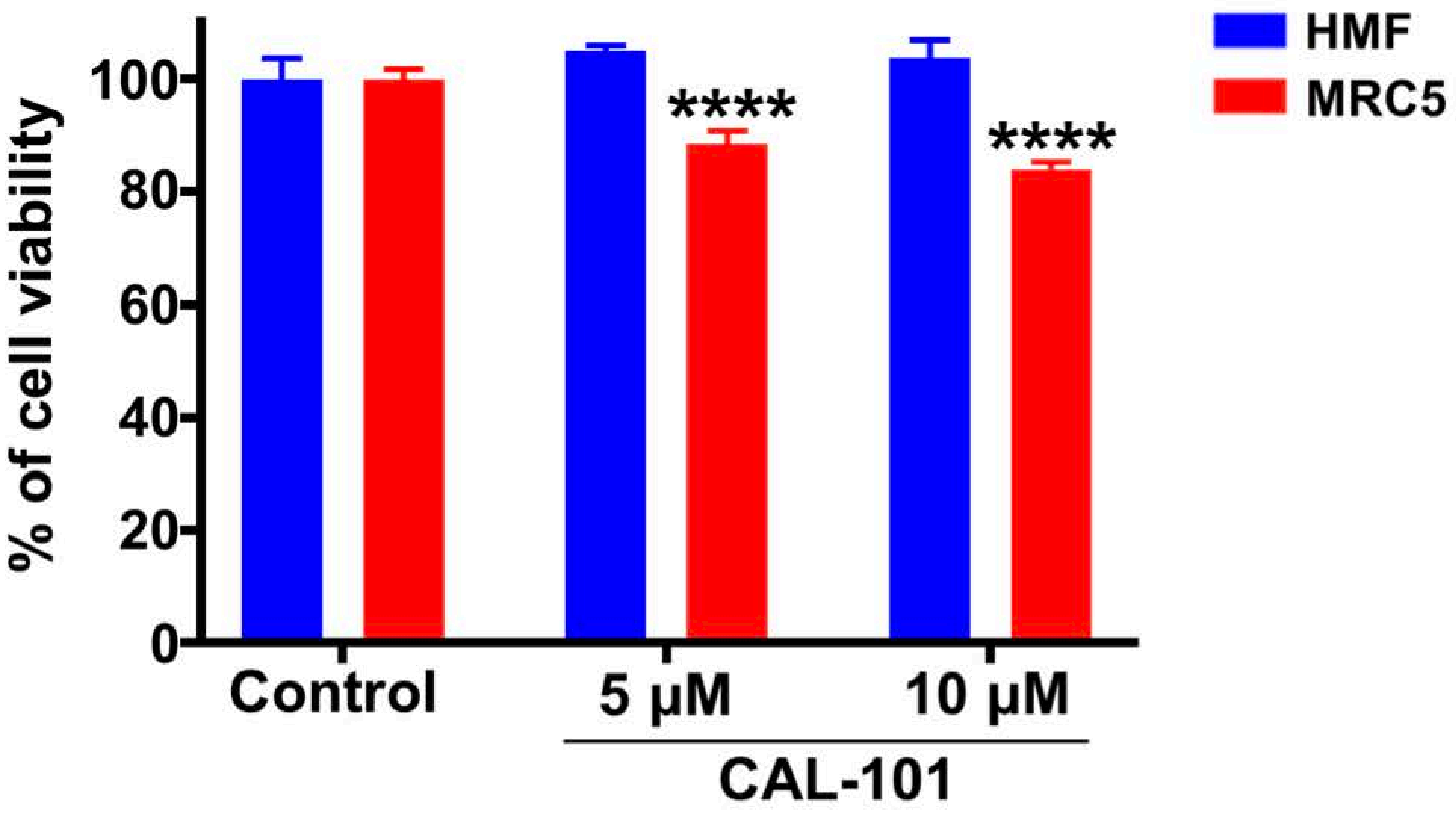

### Supplementary Figure 8

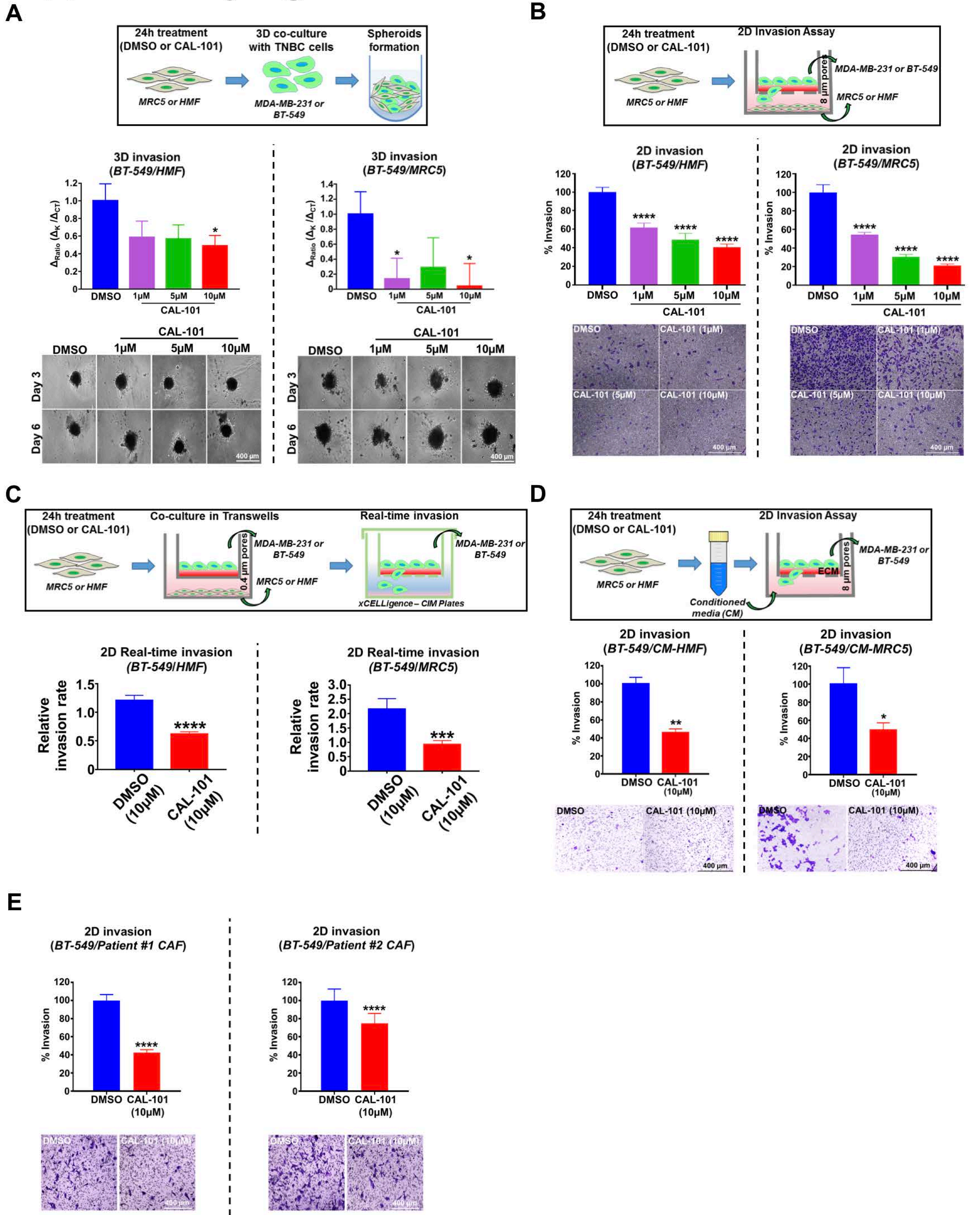

### Supplementary Figure 9

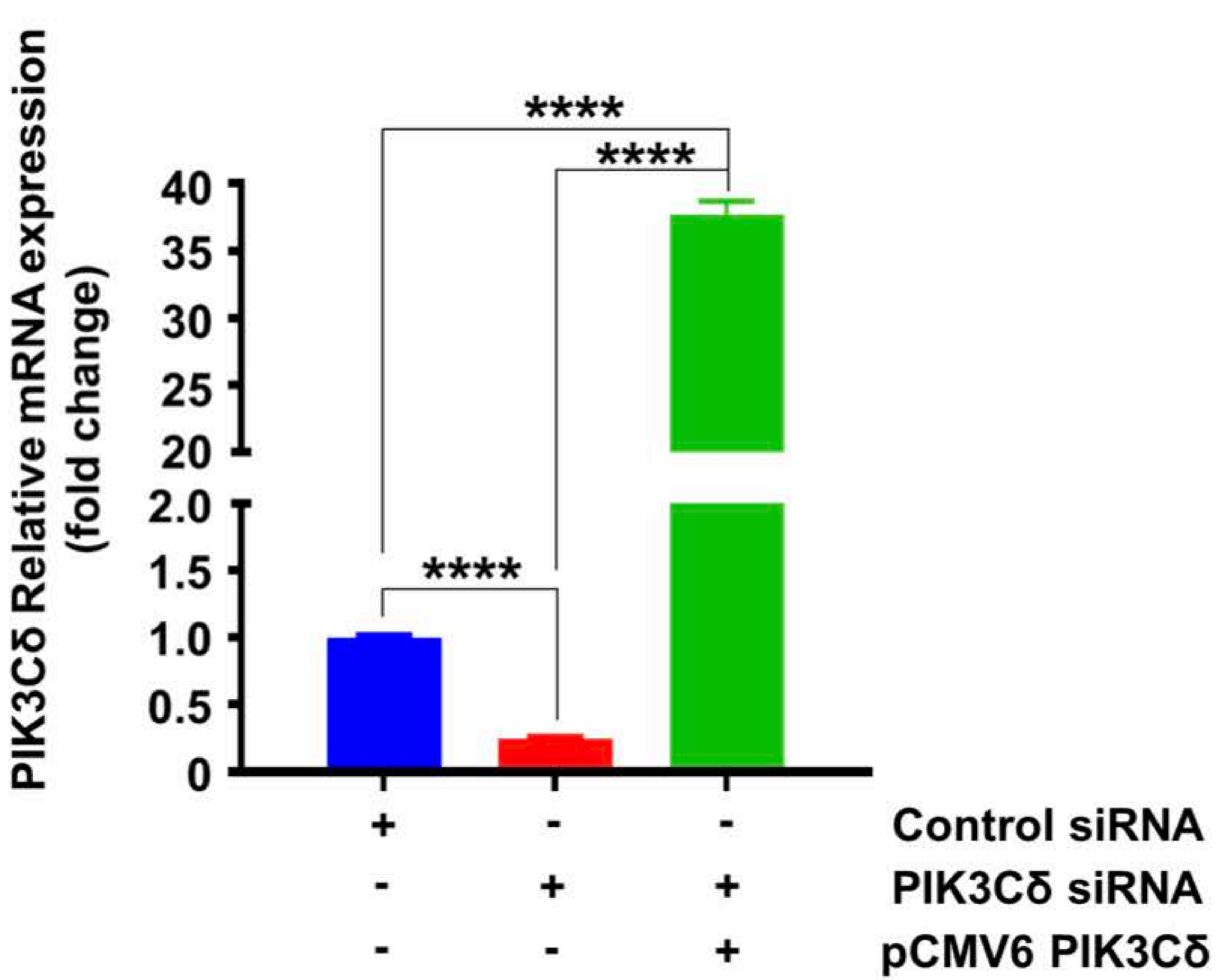


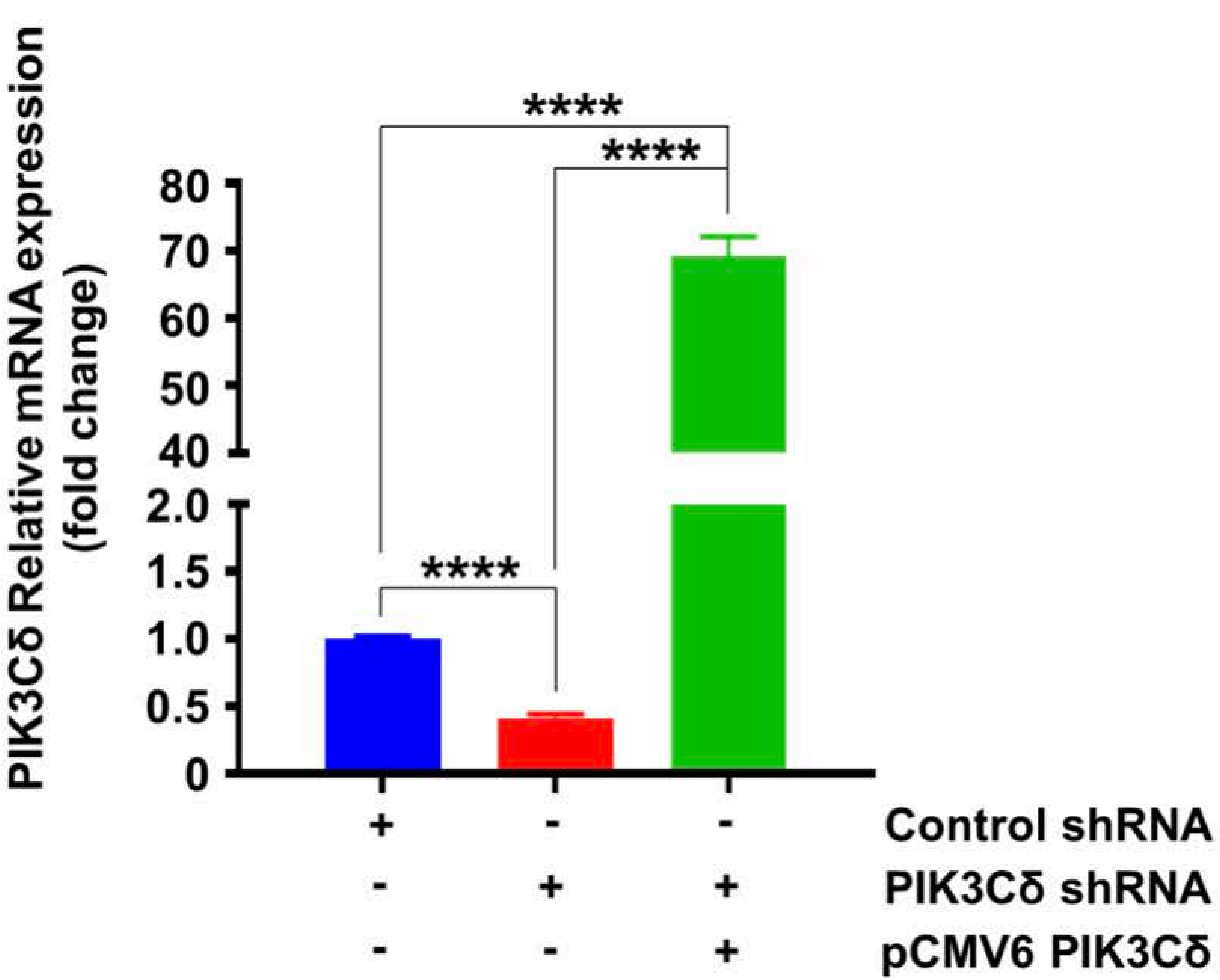

### Supplementary Figure 10

**A**

| *PIK3C isoform* | PIK3 inhibitor | | | | |
|---|---|---|---|---|---|
| | **AS252424** | **PI-103** | **CAL-101** | **Leniolisib (CDZ 173)** | **NVP-BEZ235** |
| **α** | + | ++++ | | + | ++++ |
| **β** | | ++++ | | + | ++ |
| **γ** | ++ | +++ | ++ | + | ++++ |
| **δ** | | ++++ | ++++ | ++++ | +++ |

**B**

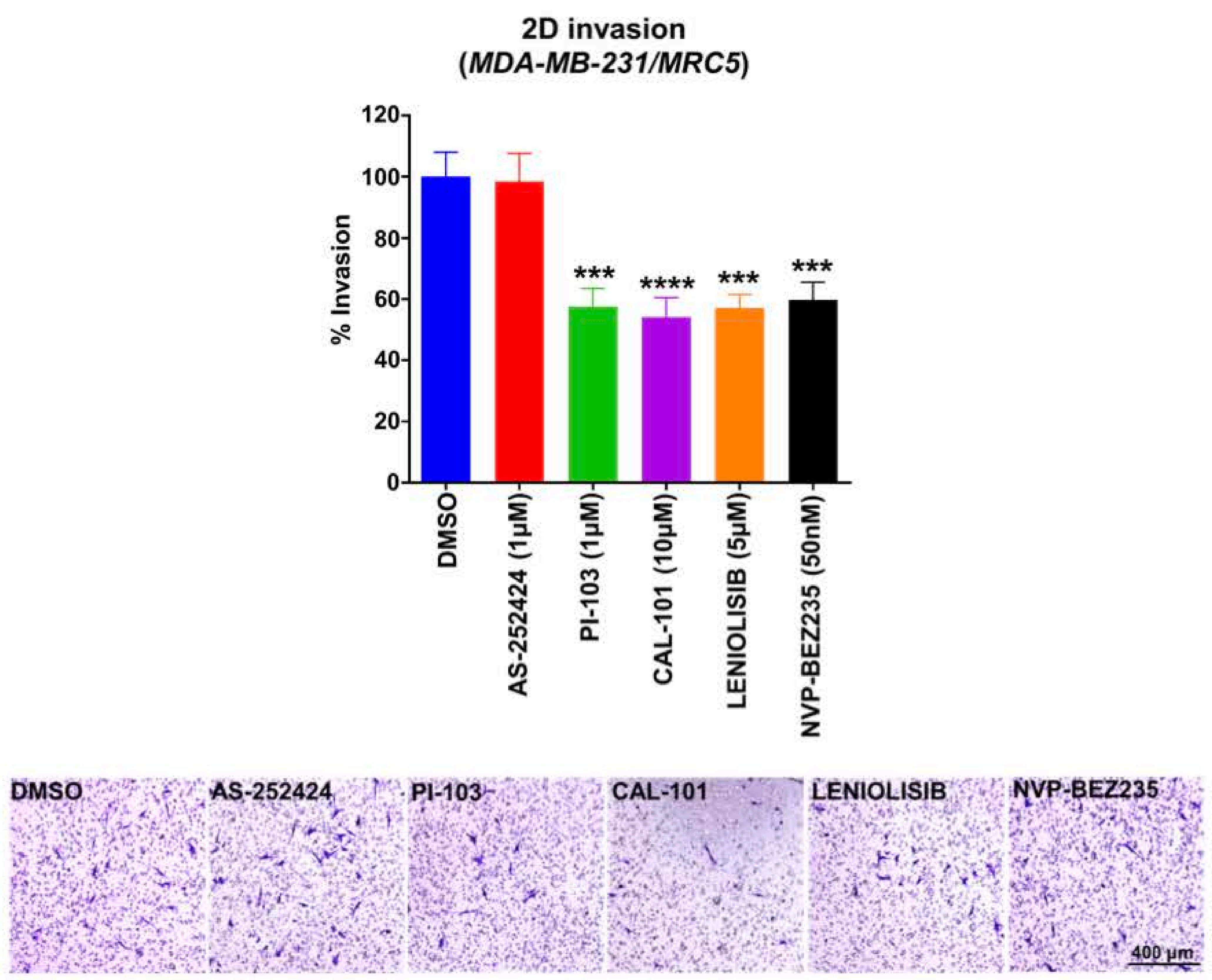


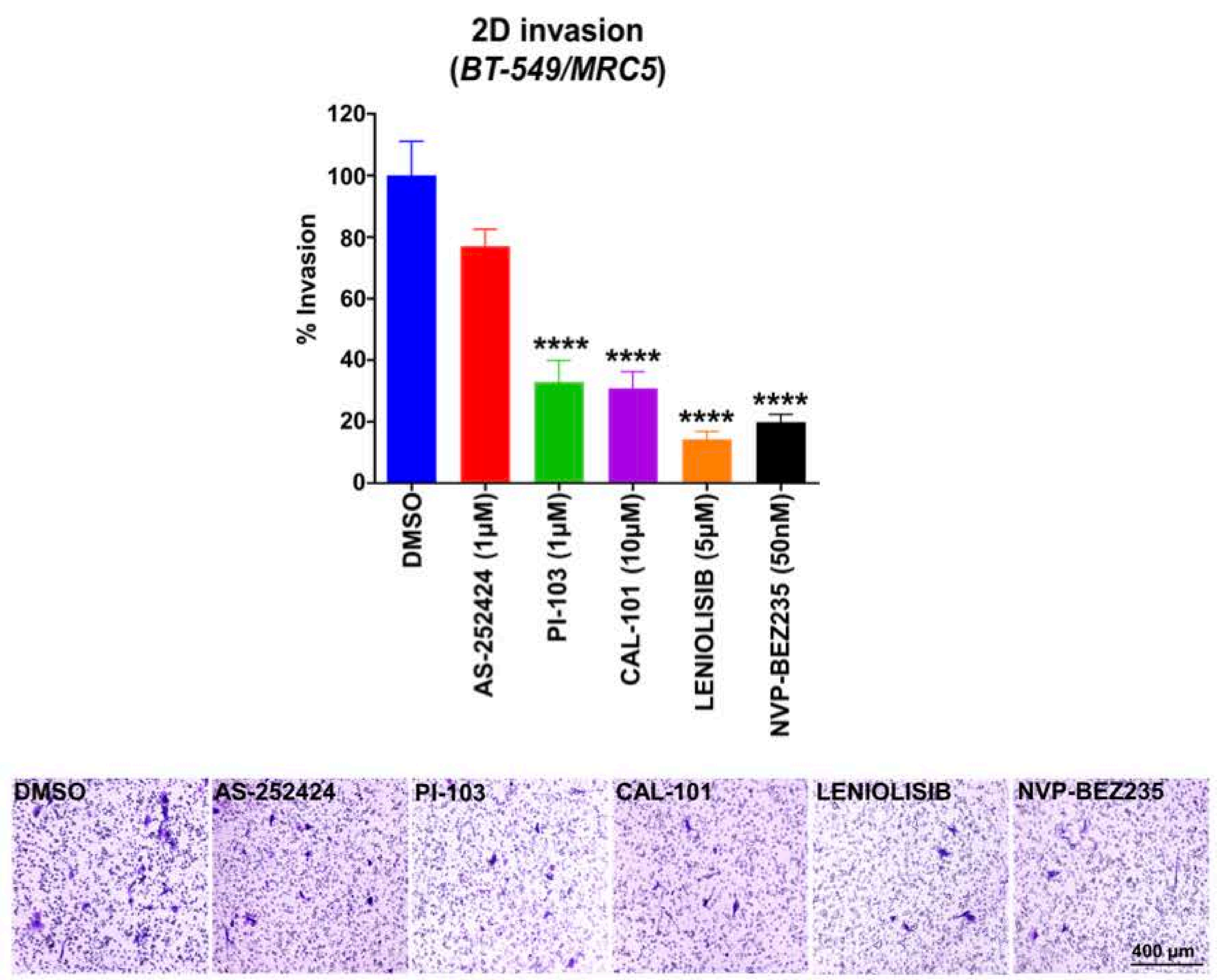


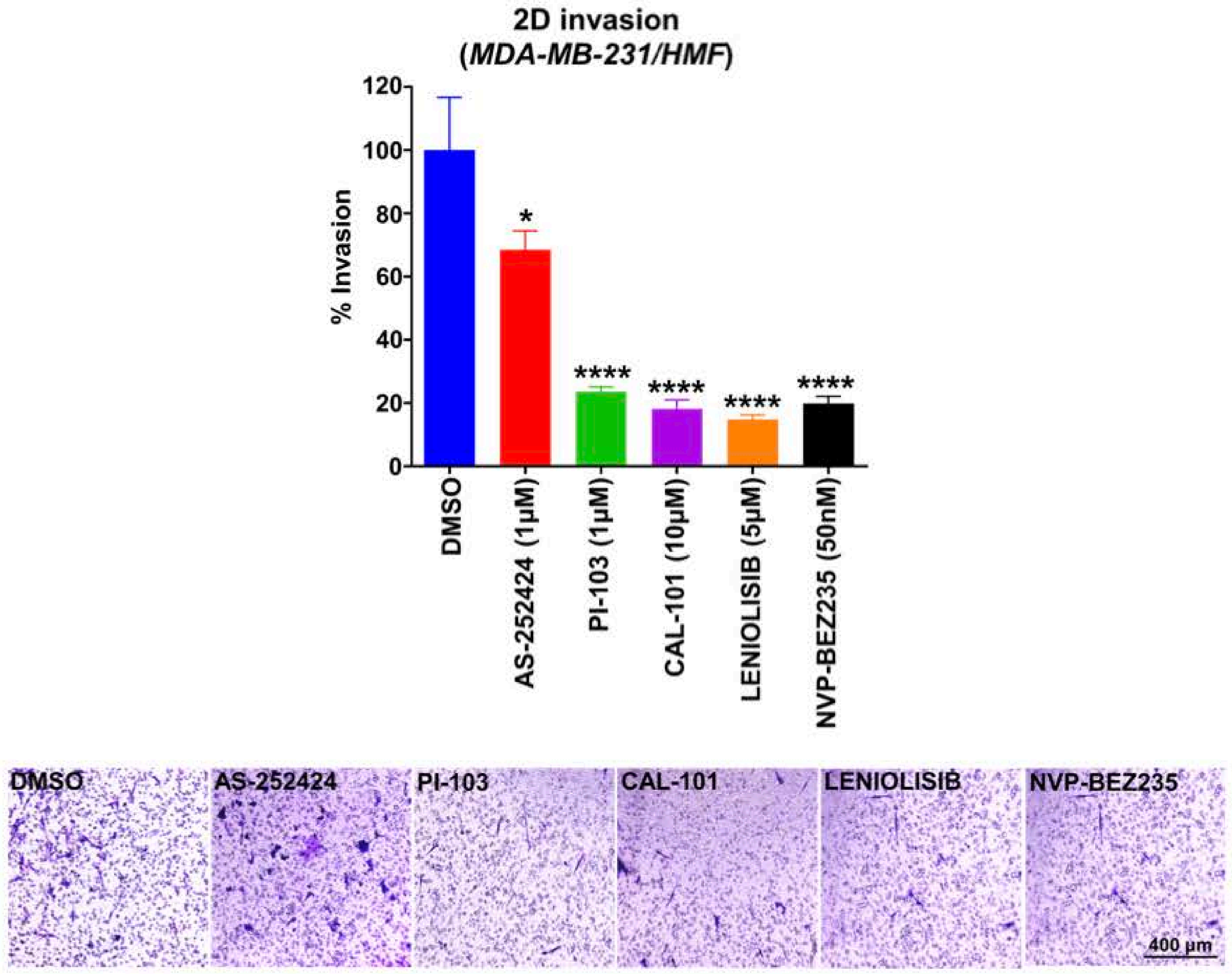


**C**

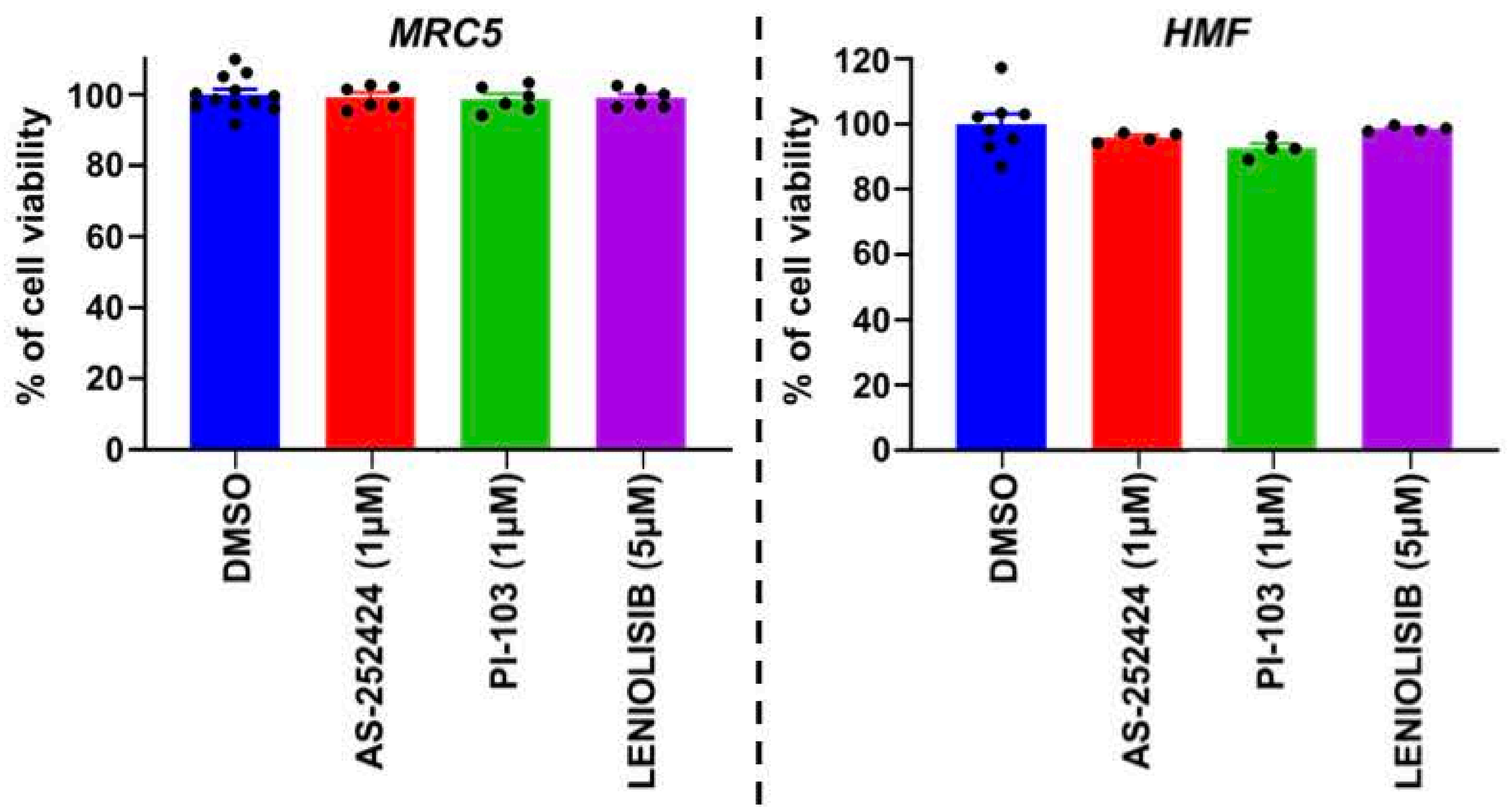

### Supplementary Figure 11

A

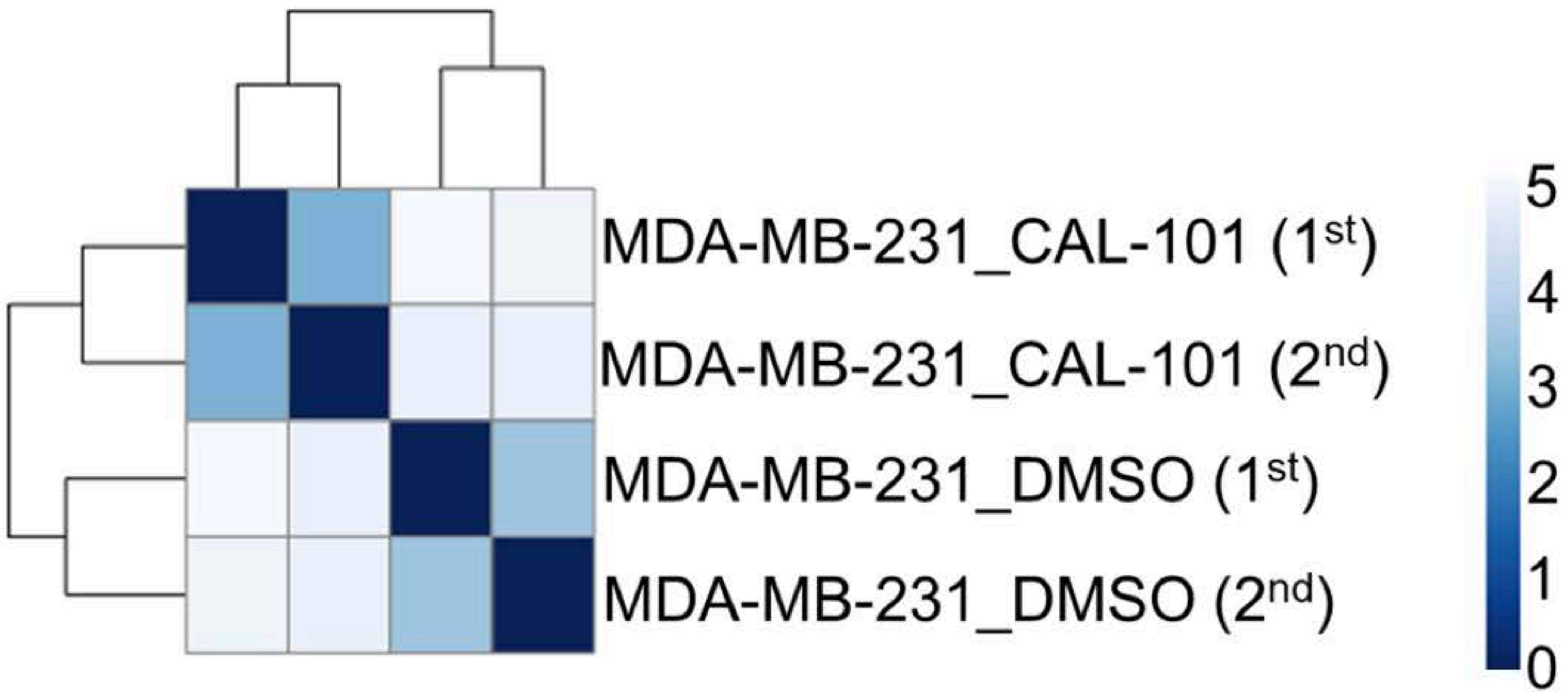


B

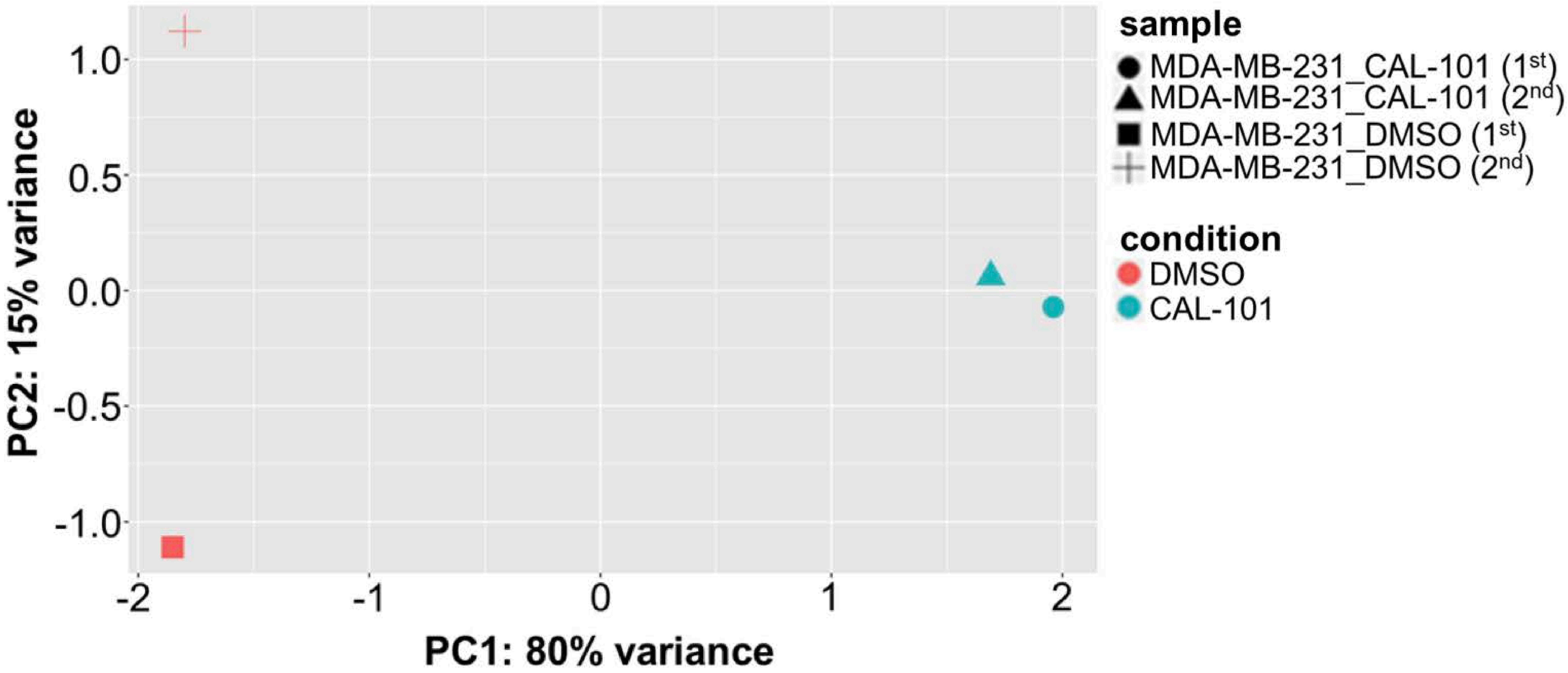

### Supplementary Figure 12

A

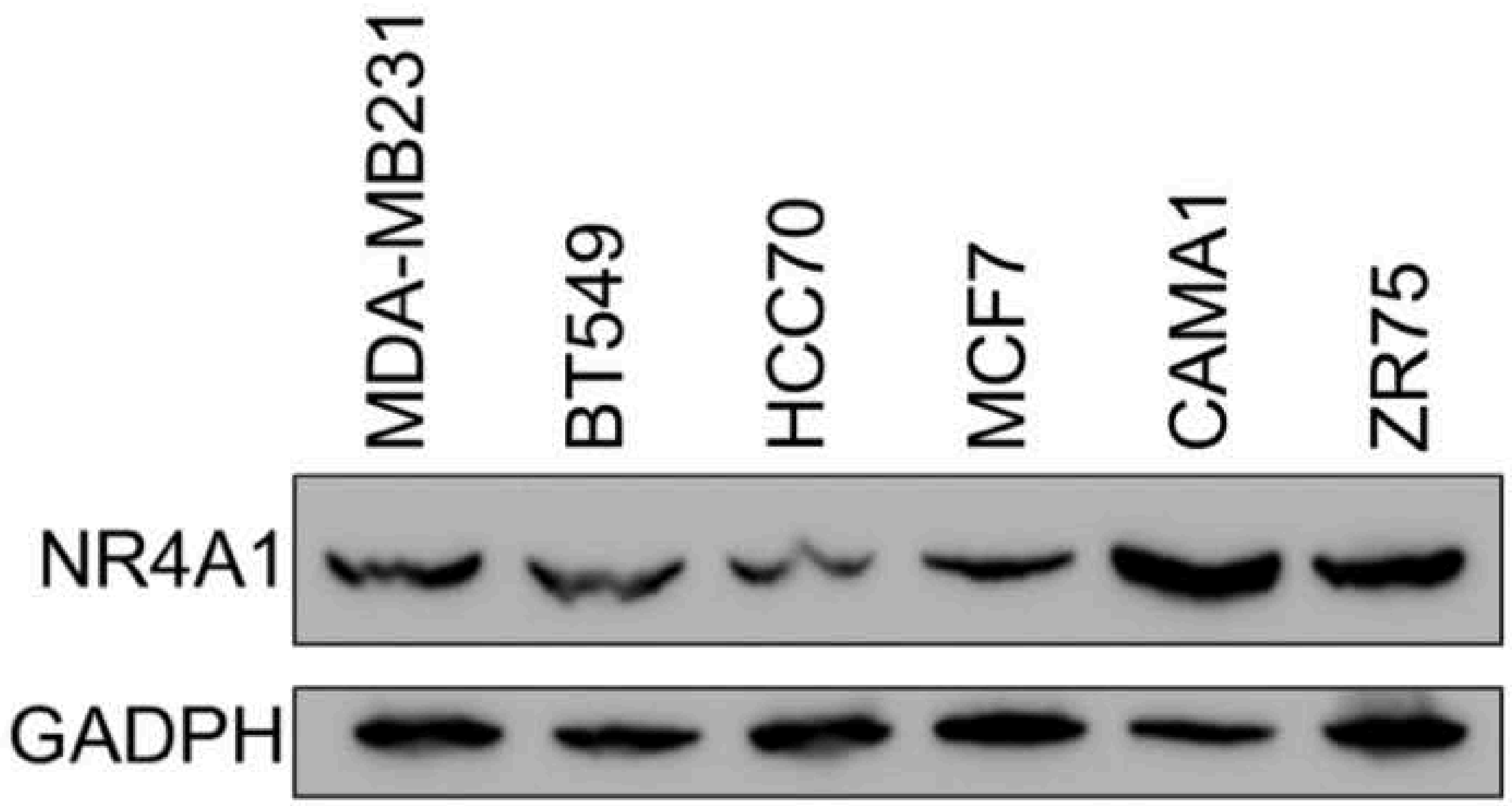


B

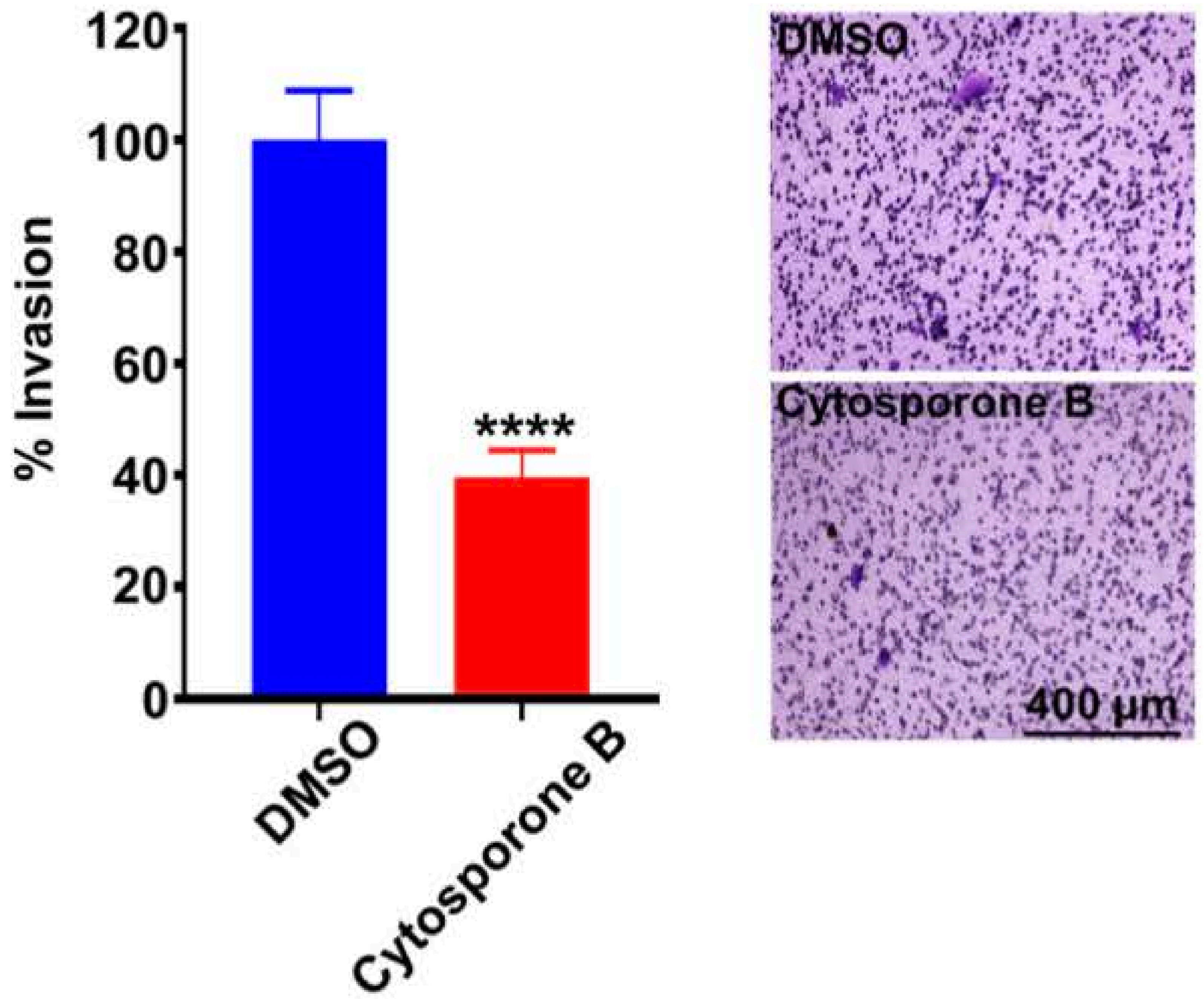


C

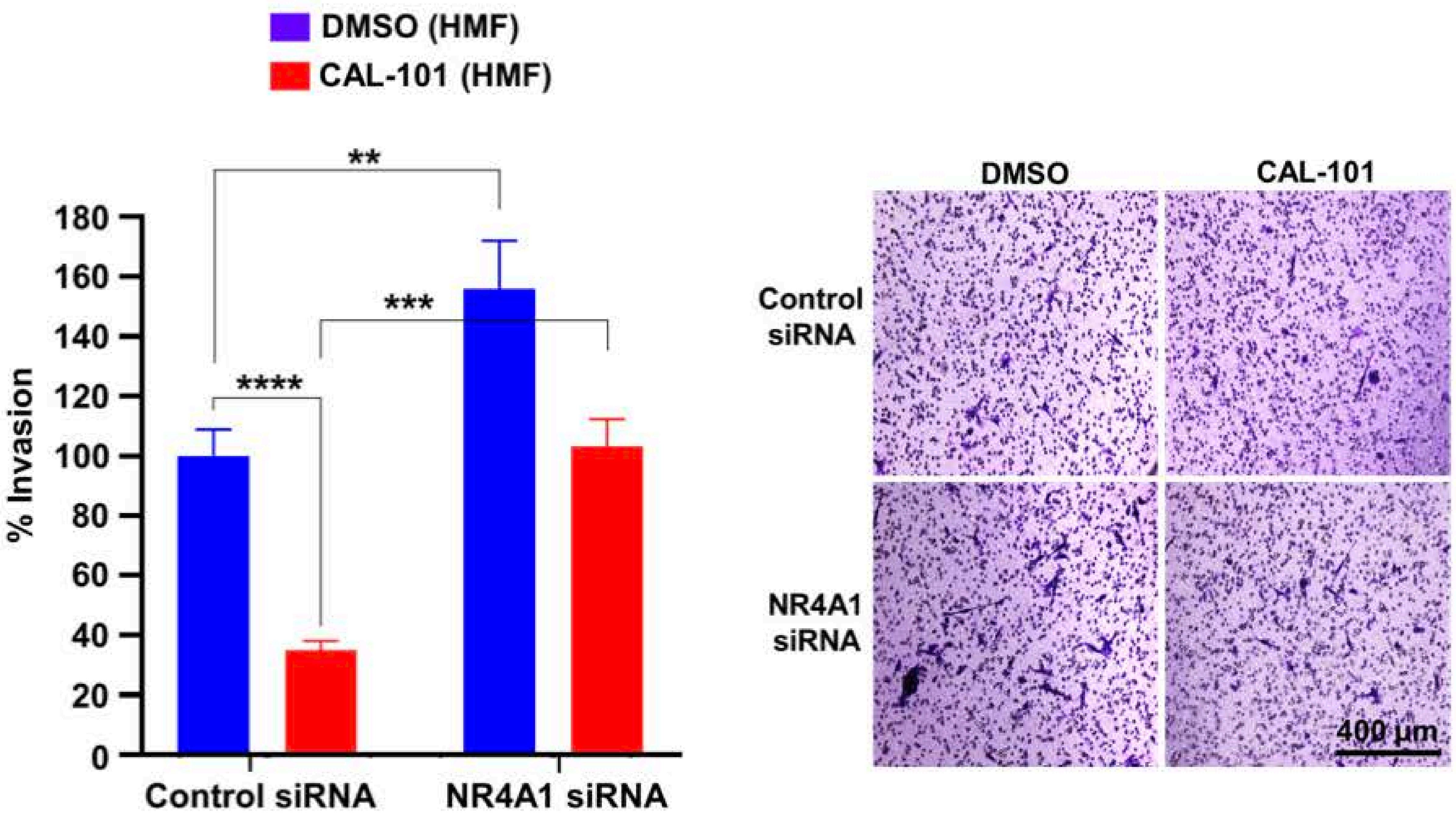

### Supplementary Figure 13

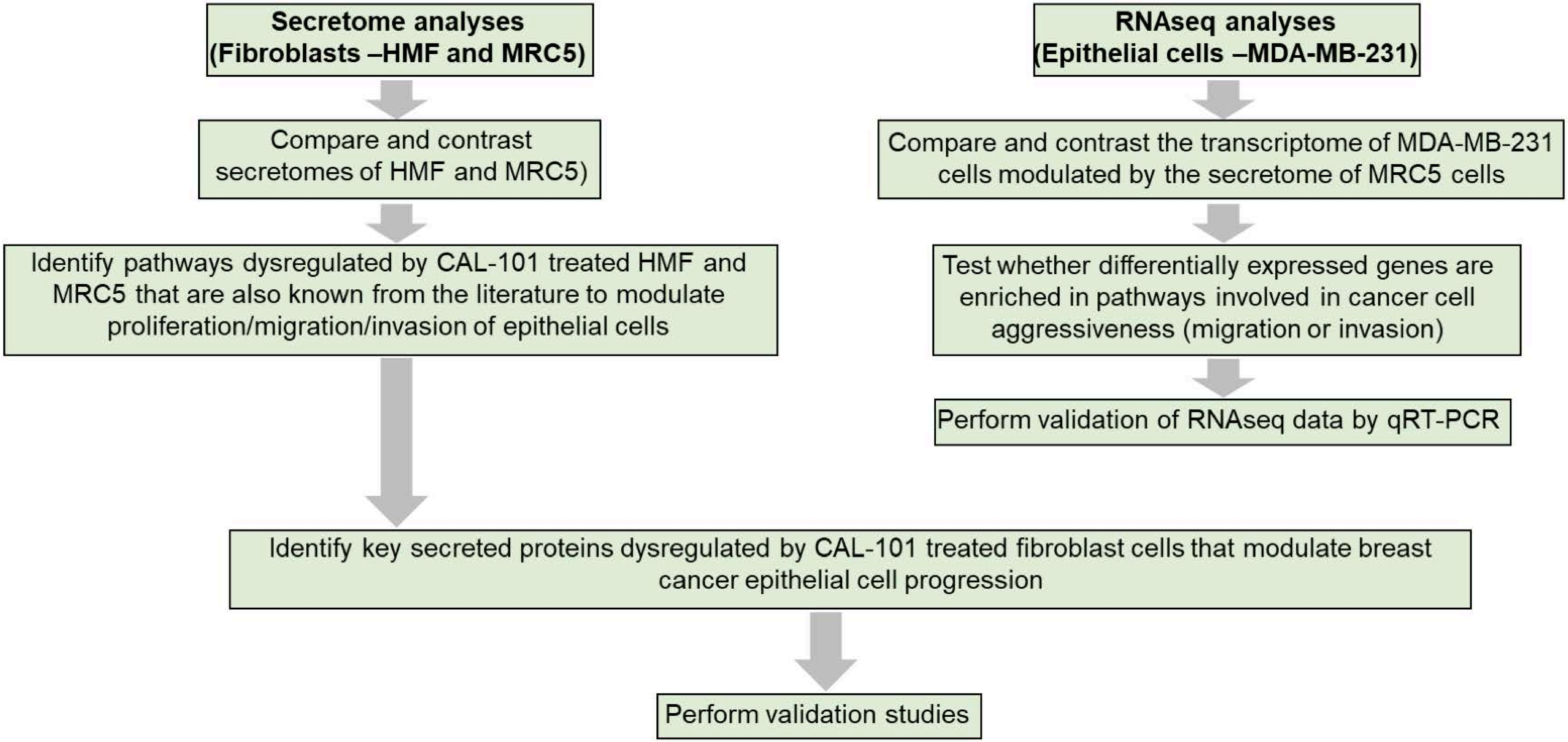

### Supplementary Figure 14

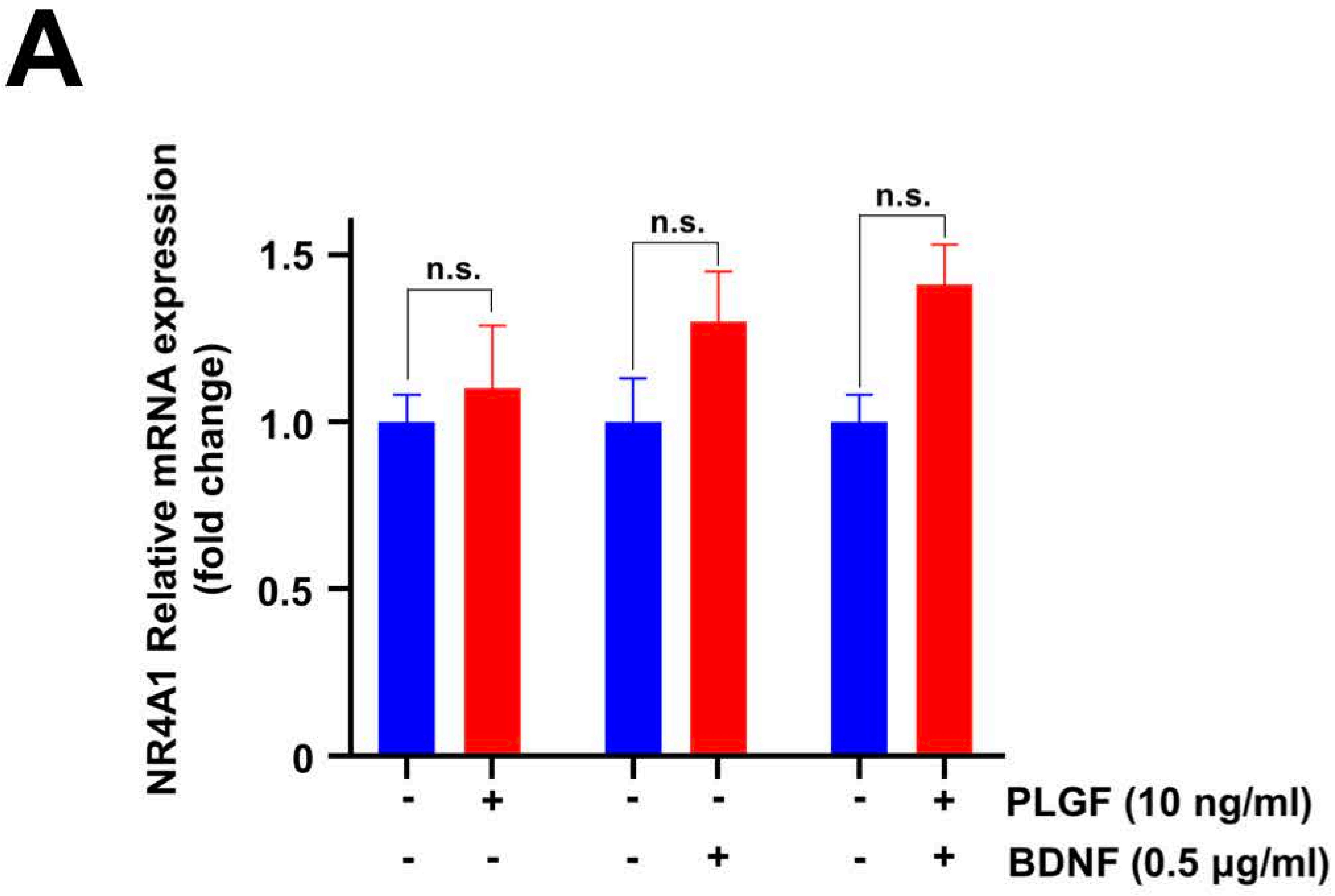


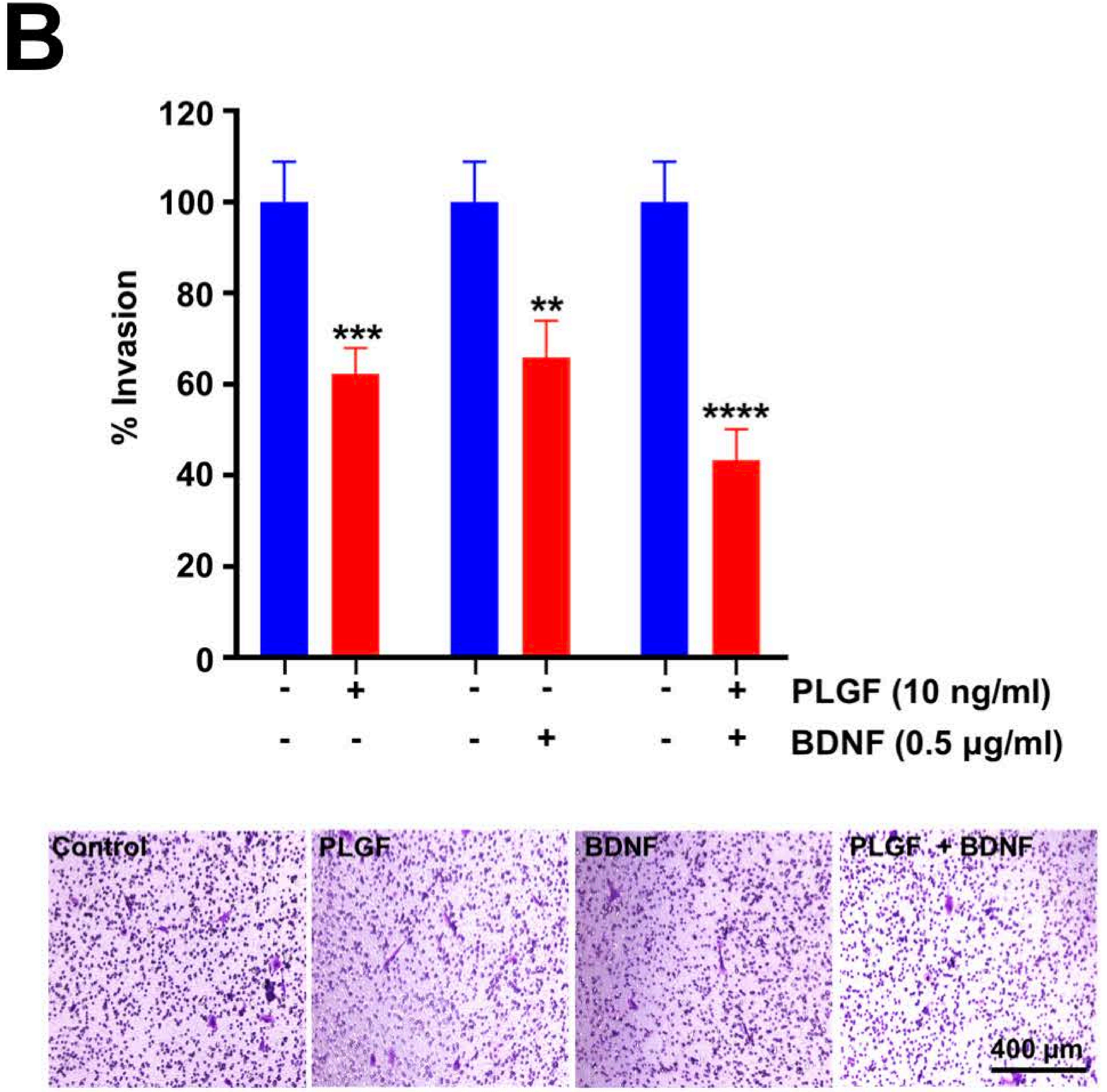

### Supplementary Figure 15

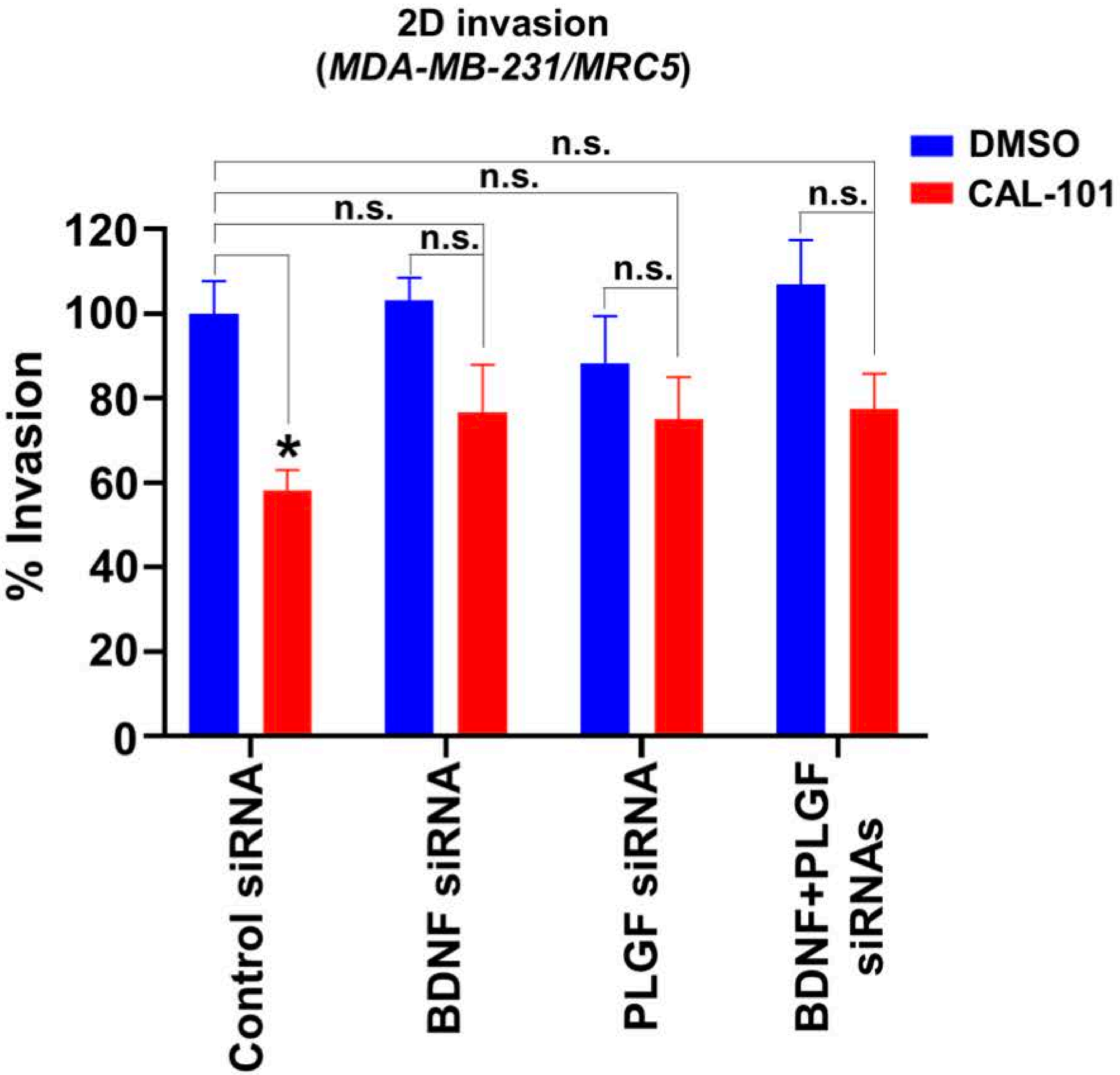


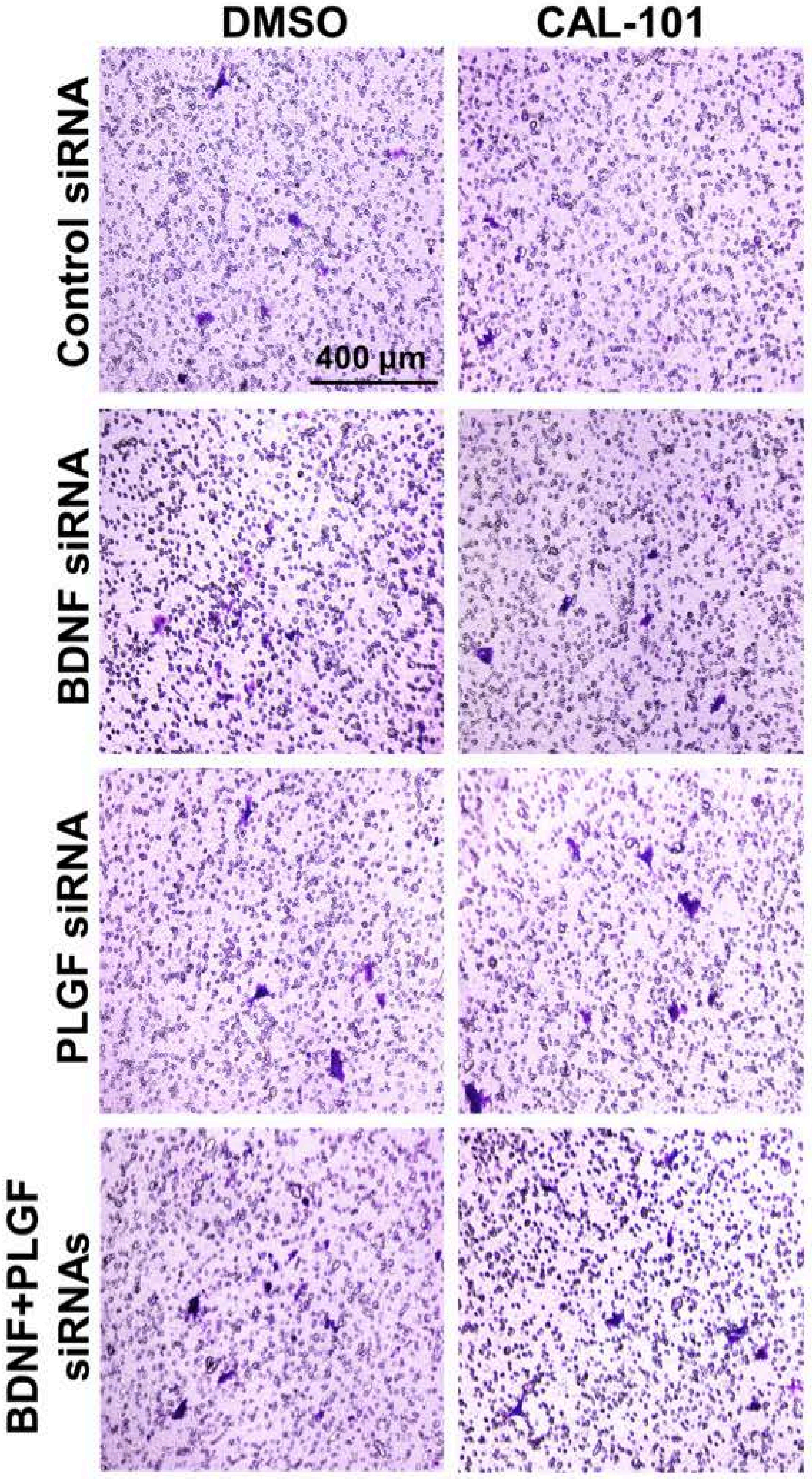

### Supplementary Figure 16

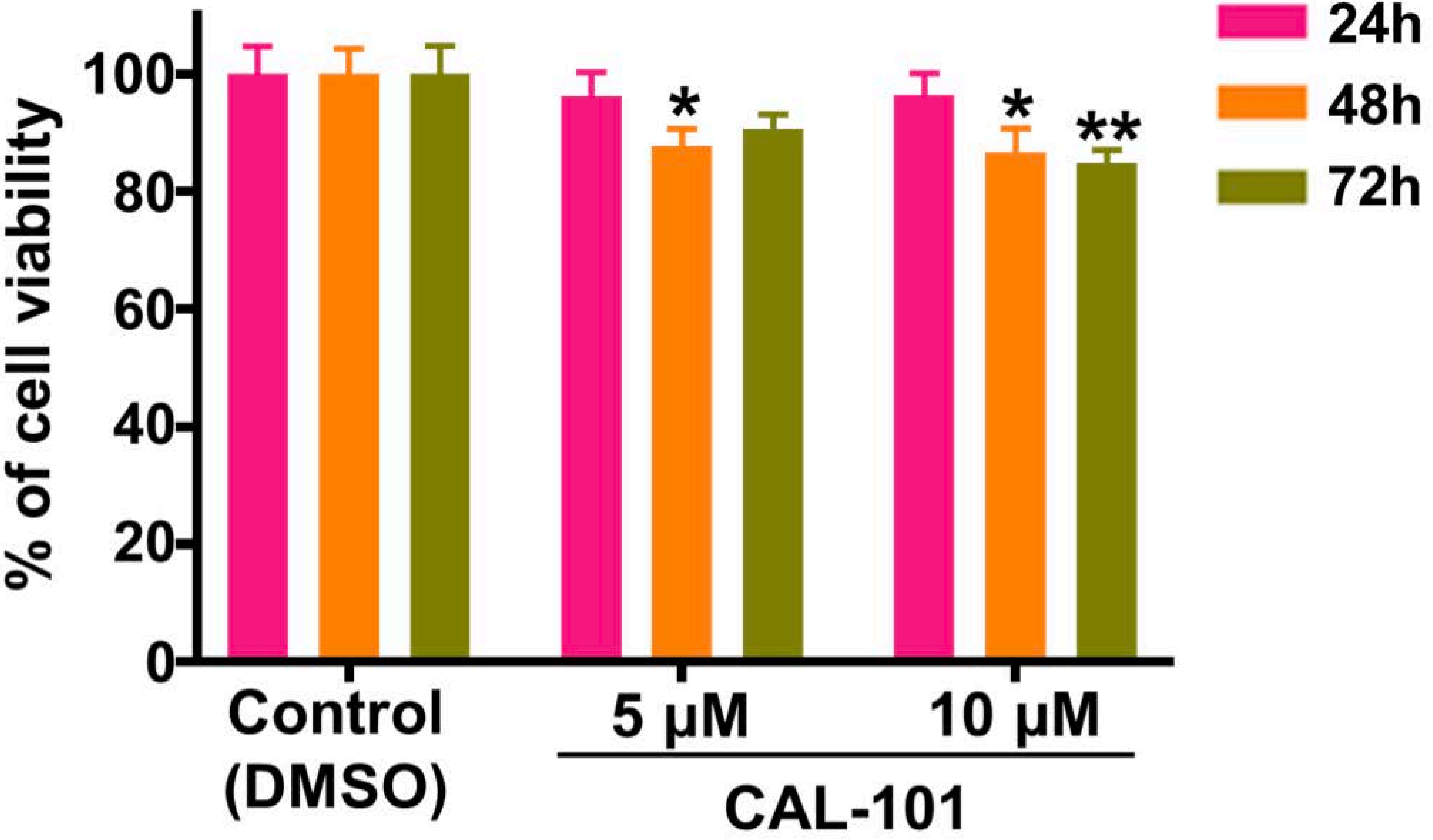

### Supplementary Figure 17

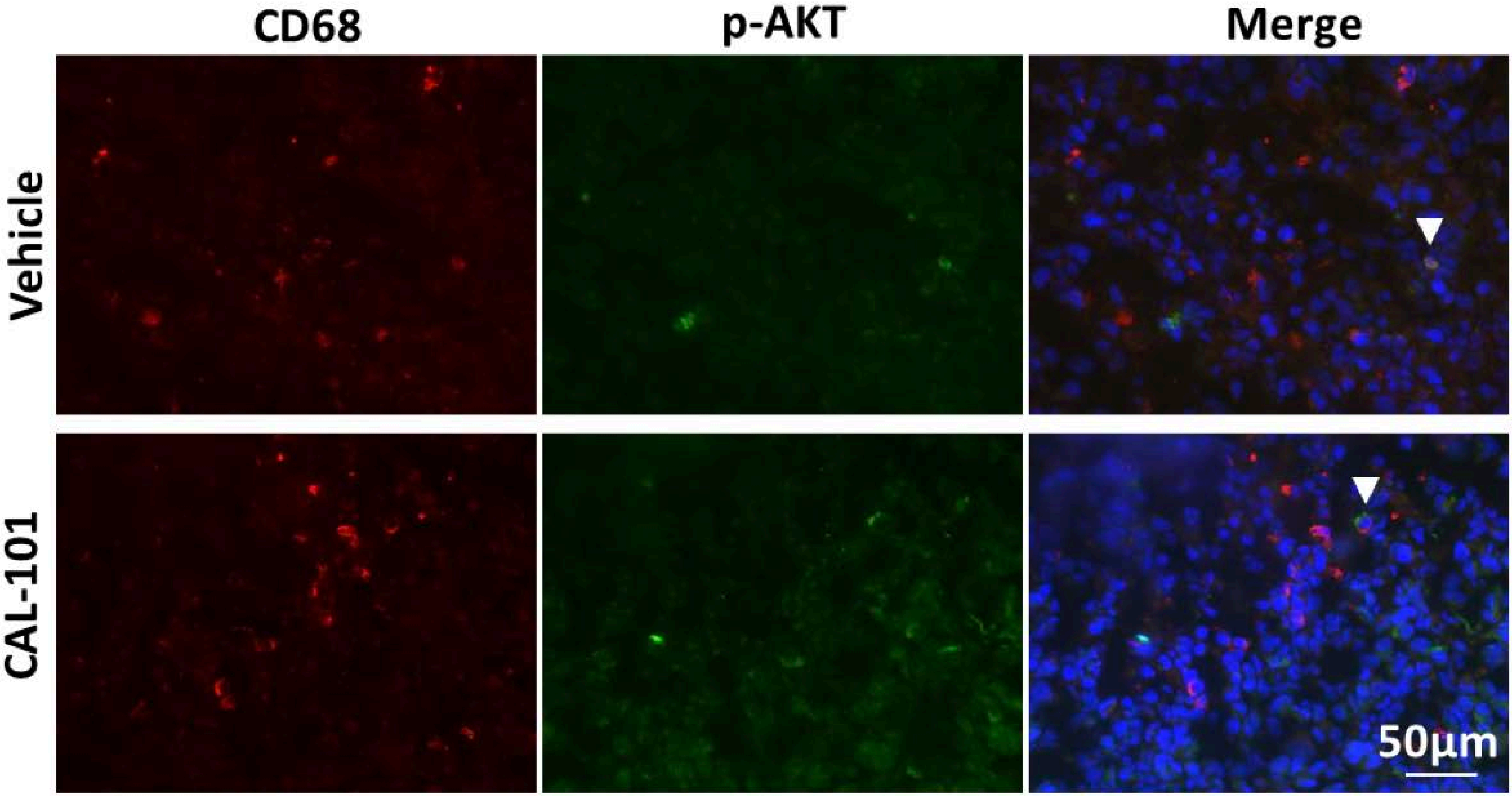

### Supplementary Figure 18

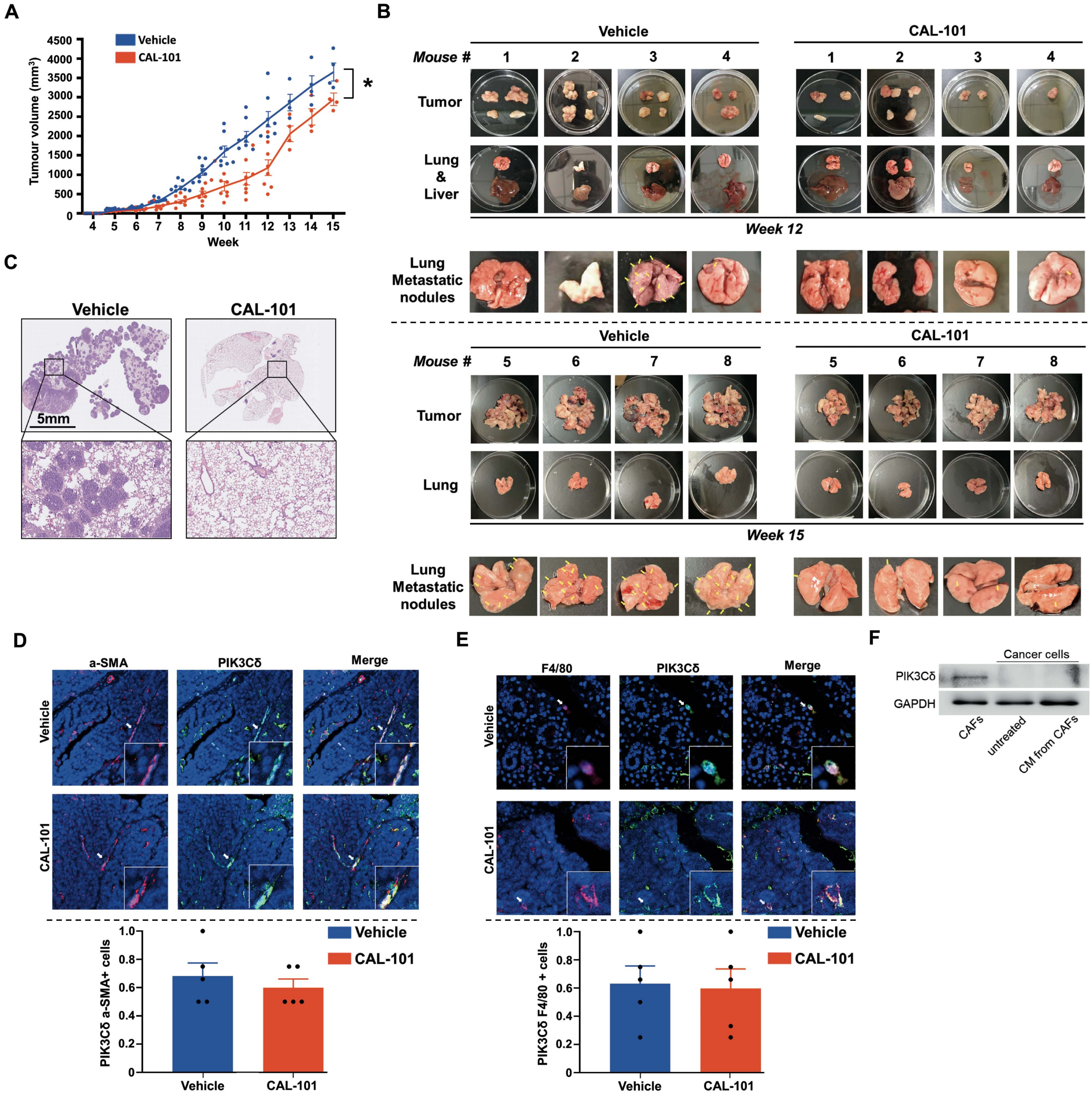

Supplementary Figure 19

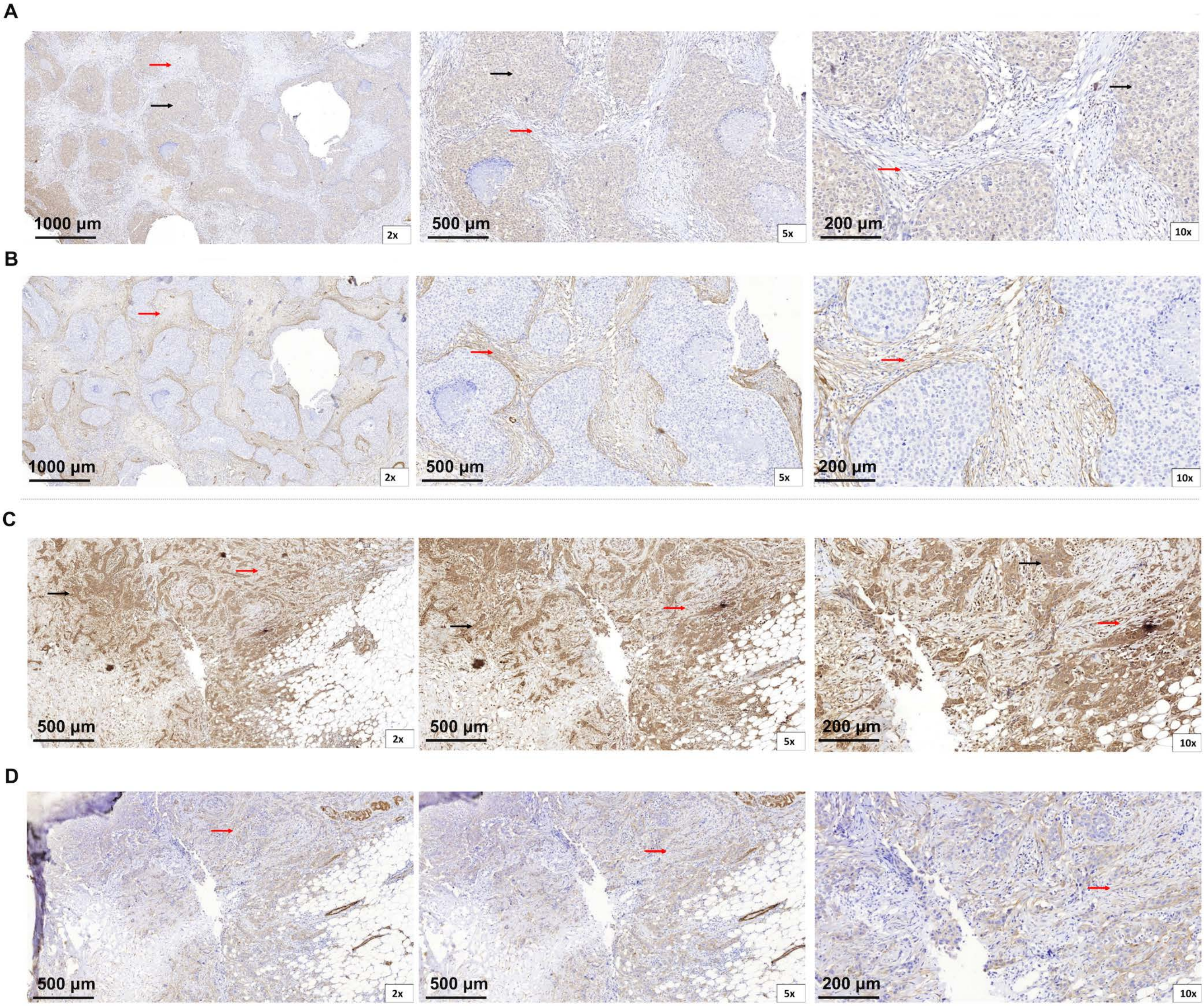

### Supplementary Figure 20

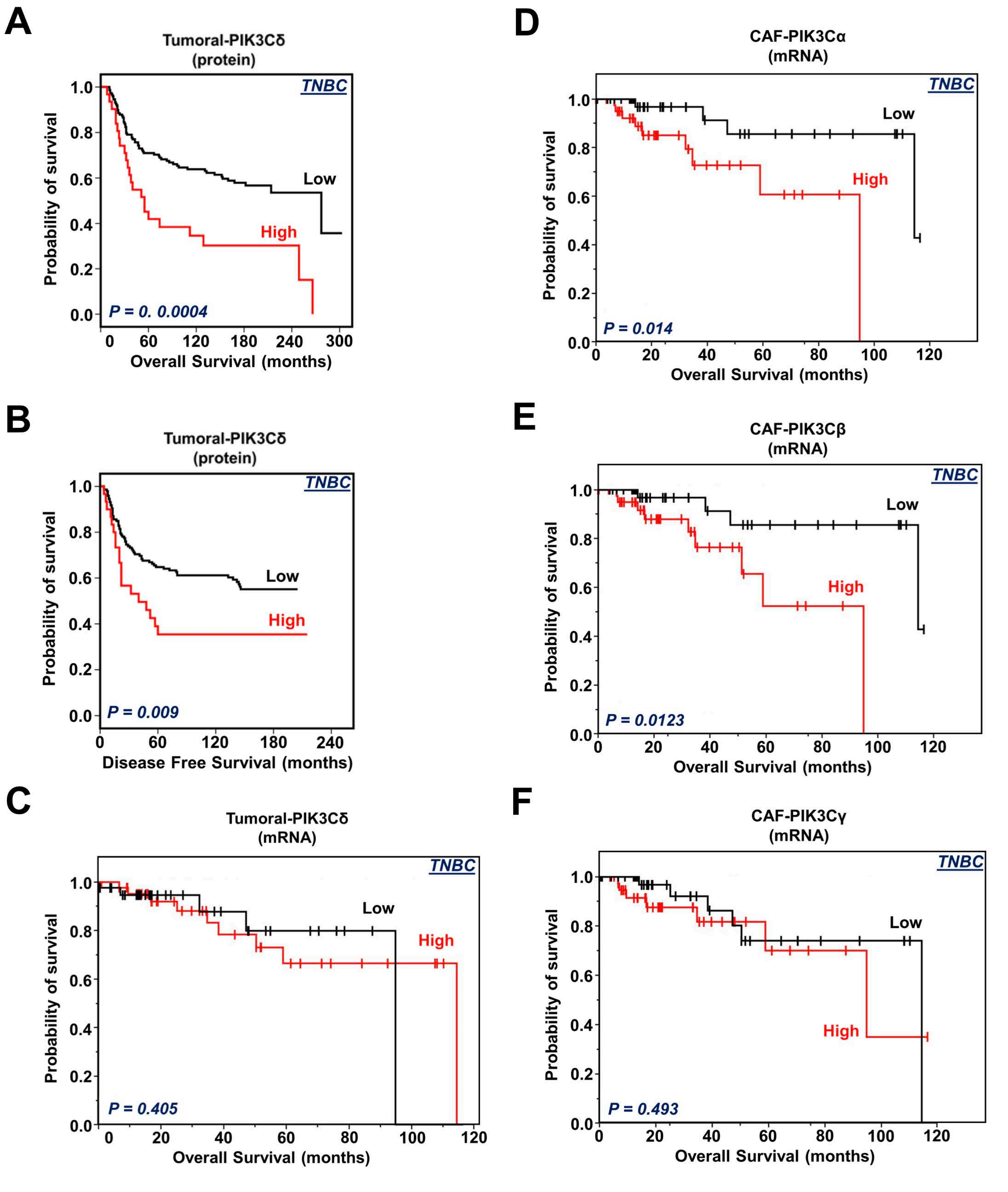
